# Total testosterone is not associated with muscle mass, function or exercise adaptations in pre-menopausal females

**DOI:** 10.1101/2024.05.12.593786

**Authors:** Sarah E. Alexander, Briana Gatto, Olivia E. Knowles, Ross M. Williams, Kinga N. Fiebig, Paul Jansons, Paul Della Gatta, Andrew Garnham, Nir Eynon, Glenn D. Wadley, Brad Aisbett, Danielle Hiam, Séverine Lamon

## Abstract

**Key points:**

- Bioavailable testosterone was positively related to exercise-induced muscle hypertrophy in pre-menopausal females.
- Nuclear localisation of the androgen receptor was positively related to muscle mass in pre-menopausal females before resistance training, but not with resistance-training induced hypertrophy.
- Total testosterone is not related to muscle mass or strength in pre-menopausal females.
- Testosterone treatment induced androgen receptor nuclear translocation, but did not induce mTOR signalling in myocytes from pre-menopausal females.

Testosterone, the major androgen hormone, influences the reproductive and non-reproductive systems in males and females via binding to the androgen receptor (AR). Both circulating endogenous testosterone and muscle AR protein content are positively associated with muscle mass and strength in males, but there is no such evidence in females. Here, we tested whether circulating testosterone levels were associated with muscle mass, function, or the muscle anabolic response to resistance training in pre-menopausal females.

Twenty-seven pre-menopausal, untrained females (aged 23.5 ± 4.8) underwent a 12-week resistance training program. Muscle strength, size, power and plasma and urine androgen hormone levels were measured. Skeletal muscle biopsies were collected before and after the training program to quantify the effect of resistance training on AR protein and mRNA content, and nuclear localisation. Primary muscle cell lines were cultured from a subset (n=6) of the participants’ biopsies and treated with testosterone to investigate its effect on myotube diameter, markers of muscle protein synthesis and AR cellular localisation.

Total testosterone was not associated with muscle mass or strength at baseline or with the changes in muscle mass and strength that occurred in response to resistance training. *In vitro,* supra-physiological doses of testosterone increased myocyte diameter, but this did not occur via the Akt/mTOR pathway as previously suggested. Instead, we show a marked increase in AR nuclear localisation with testosterone administration. In conclusion, we found that, bioavailable testosterone and the proportion of nuclear-localised AR, but not total testosterone, with skeletal muscle mass and strength in pre-menopausal females.

## 1. Introduction

The maintenance of skeletal muscle mass and function is an essential component of health and ageing (McLeod *et al*., 2016; Tieland *et al*., 2018). Muscle mass and function are also performance-determining factors in many sporting disciplines that rely on speed, power or strength, including sprinting (Barbieri *et al*., 2017) or weightlifting (Zaras *et al*., 2020). Skeletal muscle dynamically reacts and adapts to external stimuli such as mechanical loading and unloading, or internal stimuli such as the hormonal milieu (Schiaffino *et al*., 2013). Testosterone is an androgen (i.e., “male making”) sex hormone with anabolic properties. Females typically exhibit testosterone concentrations that are 10-fold lower than male concentrations (0.5-2.5 nmol·L^-1^ and 10-30 nmol·L^-1^, respectively) (Burger, 2002). The majority of testosterone circulates bound to carrier proteins, sex hormone binding globulin (SHBG; approximately 45%) or albumin (approximately 50%) (Burger, 2002). Only a small fraction of testosterone (approximately 3-5%) is “free” and unbound. When bound to SHBG, testosterone is not biologically active (Krakowsky & Grober, 2015). In contrast, testosterone is weakly bound to albumin and can easily dissociate. Therefore, albumin-bound and free testosterone are considered “bioavailable” and can enter target cells and bind to the androgen receptor (AR) (Burger, 2002). The AR is ubiquitously expressed and, as such, testosterone plays a role in many tissues throughout the body, including skeletal muscle.

The AR exists either in the cytosol of cells bound to chaperone proteins (Berns *et al*., 1986; Olea *et al*., 1990; de Launoit *et al*., 1991), or in the sarcolemma of myocytes linked to a G-protein coupled receptor (Dent *et al*., 2012). AR exerts its effects via 2 known mechanisms: non-genomic and genomic signalling. Non-genomic signalling refers to the process by which sarcolemma-bound ARs activate the protein kinase B/mammalian target of rapamycin (Akt/mTOR) or mitogen-activated protein kinase (MAPK) pathways to increase protein synthesis. This has been shown in rat L6 (Wu *et al*., 2010; White *et al*., 2013) and mouse C2C12 (Basualto-Alarcón *et al*., 2013) myocytes *in vitro*, but it is unknown whether testosterone signals through these pathways in humans. Genomic signalling is a process by which cytosolic AR become phosphorylated, dissociate from chaperone proteins and translocate to the nucleus of the cell (nAR). nAR act as a transcription factor that increases the expression of over 1000 target genes containing an androgen response element (ARE) in their promoter (Jin *et al*., 2013; Leung & Sadar, 2017).

Testosterone and its bioactive metabolite dihydrotestosterone (DHT) exhibit anabolic properties. The administration of exogenous testosterone promotes a positive muscle protein turnover and increased muscle mass and function in young (Bhasin *et al*., 2001) or old males (Storer *et al*., 2017) and in pre-(Hirschberg *et al*., 2020) or post-menopausal females (Huang *et al*., 2014). Conversely, when testosterone concentrations are pharmacologically suppressed, the protein balance switches in favour of protein degradation (Ferrando *et al*., 1998; Sheffield-Moore *et al*., 1999) and leads to reduced muscle mass and strength in males (Mauras *et al*., 1998; Overkamp *et al*., 2023). There is evidence of positive associations between endogenous total testosterone and muscle mass or strength in large male cohorts across the lifespan (*n*=252 (Mouser *et al*., 2016), *n*=3,875 (Ye *et al*., 2021)). Other, smaller studies refute the existence of such an association (*n*=49 (Morton *et al*., 2018), *n*=23 (Mitchell *et al*., 2013), *n*=67 (Mobley *et al*., 2018), *n*=49 (Morton *et al*., 2016)) and instead propose that increased skeletal muscle AR protein content (Morton *et al*., 2018), or nAR (Hatt *et al*., 2024), but not total or bioavailable circulating testosterone (Morton *et al*., 2016; Morton *et al*., 2018), is associated with increased muscle hypertrophy and function in young males.

Our knowledge of the association between endogenous testosterone and muscle mass is limited in females (Alexander *et al*., 2022) and the available evidence stems from cross-sectional cohorts. Evidence from murine (Yoshioka *et al*., 2007) and human (Pataky *et al*., 2023; Hatt *et al*., 2024) models however suggest that there are sex-specific differences in androgen action on the skeletal muscle transcriptome. We and others show that there is no association between endogenous total testosterone or nAR and muscle mass and strength in pre-(Alexander *et al*., 2021; Hatt *et al*., 2024) or post-menopausal females (Gower & Nyman, 2000; Carmina *et al*., 2009; van Geel *et al*., 2009; Pöllänen *et al*., 2011; Rariy *et al*., 2011; Kogure *et al*., 2015). Instead, the free androgen index (FAI), which is indicative of the amount of bioavailable testosterone, is weakly associated with muscle mass in pre-menopausal females (Carmina *et al*., 2009; Alexander *et al*., 2021).

The aim of the current study was to investigate the association between total testosterone and muscle mass, strength and power in pre-menopausal females at baseline or with the changes in muscle mass, strength or power following a tightly controlled 12-week resistance exercise training protocol. A secondary aim was to identify whether the FAI is more closely associated with muscle mass, strength, power or anabolic potential. We also investigated whether markers of AR expression, activity or localisation, or markers of muscle protein synthesis or degradation were associated with muscle mass, strength, power or anabolic potential. An *in vitro* model was used to further our mechanistic understanding of the role of testosterone in female human primary myocytes.

## 2. Methods

### 2.1 Ethical approval

This research was granted ethical approval by the Deakin University Human Research Ethics Committee (DUHREC 2018-388). All participants provided written, informed consent before taking part in the study, which was conducted in accordance with the Declaration of Helsinki (World Medical Organisation, 2018) and its later amendments.

### 2.2 Participants and exclusion criteria

Thirty-five healthy females aged 18-40 years were recruited from the general population. Four participants were not able to continue the training program due to COVID-19-related interruptions in 2020, 2 participants withdrew for health-related reasons and 2 participants withdrew for personal reasons. Therefore, 27 females completed the training program. Participants were not resistance-trained (defined as having performed structured resistance training at least twice per week in the previous 6 months), pregnant or breastfeeding, did not smoke and displayed no contraindications to exercise according to the Exercise and Sports Science Australia adult pre-exercise screening system (Exercise and Sports Science Australia, 2019). Participants were excluded if they had a history of anabolic hormone use, used medications or supplements that could affect the anabolic response to training, or if their daily protein intake was outside the Australian dietary guidelines of 15-25% total macronutrient intake, measured through a mobile phone application for 4 days including 1 weekend day (Easy Diet Diary) (Xyris Software, 2019). The health, fitness and anabolic status of young, healthy females are not expected to change over a 12-week period as a passage of time. Therefore, each participant acted as her own control in a pre-post study design.

### 2.3 Assessment of confounding factors

Participants completed a chronotype questionnaire (Horne & Östberg, 1976) to assess the time of day at which they are most alert. Participant chronotype was later tested as a potential covariate in statistical analysis in case it was significantly associated with both the independent and dependent variables of interest. To monitor sleep quantity and energy expenditure, participants wore an activity monitor (Actical Z MiniMitter, Phillips Respironics Inc, Bend, OR) on their non-dominant wrist for 7 days, accompanied by a sleep diary that incorporates several validated sleep rating systems (Samn & Perelli, 1982; Jay *et al*., 2006). Sleep quantity (total hours) and total daily energy expenditure (metabolic equivalent; METs) measurements were repeated for 24 hours on weeks 3, 6 and 9 of the trial to ensure participants’ sleep and energy expenditure remained consistent, as any changes to either of these variables could potentially affect the outcome of this study. Sleep quantity and total daily energy expenditure (METs) were tested as covariates in later statistical analysis and were added to the models if they were significantly associated with both the independent and dependent variables of interest.

Protein intake, daily physical activity and sleep quantity were measured at baseline and every 3 weeks throughout the training program. Sleep and protein intake did not change significantly during the program, suggesting the participants maintained their habitual diet, and sleep patterns throughout the entire 12 weeks (Supplementary Figure 1A-B). Participants decreased their total energy expenditure by 14% during week 6 (*p*<0.001) and by 10.5% during week 9 (*p=*0.041) when compared to baseline (Supplementary Figure 1C). Despite this change in total energy expenditure, Akaike Information Criterion (AIC) tests revealed that total energy expenditure was not a significant confounder of the linear models and was therefore not included in subsequent analyses.

### 2.4 Menstrual phase standardisation and hormonal contraception use

The pre-post design of this study allowed each participant to act as her own control, therefore we did not exclude participants based on hormonal contraceptive (HC) use. This study included both normally menstruating females and females using HC. Some research suggests that muscle strength may be greater in the late follicular phase (days 7-14) compared to other phases (Knowles *et al*., 2019). More recent research however suggests that there is no difference in muscle strength between menstrual cycle phases (Colenso-Semple *et al*., 2023). Despite this, we aimed to minimise any potential confounding effect of the menstrual cycle on muscle performance by avoiding the late follicular phase (days 7-14) of the menstrual cycle during pre- and post-training testing in normally menstruating participants. The data collection period lasted 12 weeks, 3 full cycles of a typical menstrual cycle lasting 28 days, allowing each normally menstruating participant to undergo pre- and post-testing during the same phase of their cycle. Menstrual phases were verified through menstrual diaries and hormonal analysis, in line with published guidelines for the inclusion of females in exercise physiology cohorts (Knowles *et al*., 2019; Elliott-Sale *et al*., 2021). Two separate researchers verified the menstrual phase of each participant and consensus was reached in each case. The menstrual phase of each testing timepoint is summarised in Supplementary Table 1. HC use and menstrual phase were tested as covariates in all subsequent statistical analysis.

### 2.5 Familiarisation to the training program

Prior to beginning the training program, participants attended 3 familiarisation sessions at Deakin University. During these sessions, the participants were coached through all training exercises with little-to-no weight (Rate of Perceived Exertion (RPE) <3/10-“moderate”) (Borg, 1982a; Borg, 1982b) to ensure all participants used the safe and correct technique for all movements. These sessions also aimed to minimise any potential learning effects that may have occurred due to the novelty of the exercises for some participants.

### 2.6 Strength and power testing

Peak muscle power was assessed using a portable force plate (AMTI, Watertown, MA). Participants performed a countermovement jump (CMJ), without an arm swing. Four attempts were made, separated by 3 minutes rest and the highest values were recorded as peak muscle power.

Participants’ repetition maximum (RM) was assessed for leg press, as well as all the exercises included in the training program. Leg press was included in the strength testing but not in the training program and represents the major measure of muscle strength in this study. Using an exercise that was not included in the training program minimised any learning effect, as participants did not train and therefore, learn the movement. Lower body 1RM was calculated from 5RM tests using the equation (Abadie & Wentworth, 2000): estimated 1RM = 4.67 + (1.14 × weight lifted. Upper body 1RM was calculated from 10RM tests using the equation (Abadie & Wentworth, 2000): estimated 1RM = 1.43 + (1.20 × weight lifted).

### 2.7 Plasma and urine collection

Plasma was collected from participants in the fasted state before and after the training program, as well as before exercise in weeks 2, 4, 6, 8 and 10. At 0700 h, 10 mL of venous blood was taken from the antecubital vein in vacutainer tubes containing 7.2 mg K2 EDTA (Becton Dickinson, Franklin Lakes, NJ). Blood was centrifuged immediately for 10 min at 1 500 *g*, 4°C and plasma was stored at −80°C until further use. A first-void urine sample was collected at the same time points and the urine was stored at −20°C until further use.

### 2.8 Body composition analysis

Participants’ body composition was assessed before and after the 12-week training program via bioelectrical impedance analysis (BIA; Tanita, Kewdale, WA) and dual-energy X-ray absorptiometry (DXA; Lunar Prodigy Advance, GE Healthcare, Madison, WI). At the time of measurement, participants had abstained from vigorous exercise, caffeine, and alcohol for the previous 48 hours, minimising the chances of water-retention or dehydration that may occur, as per standard recommendations (Walter-Kroker *et al*., 2011).

### 2.9 Assessment of thigh muscle cross sectional area

The cross-sectional area (CSA) of the thigh muscle groups (quadriceps and hamstrings) at 50% of femur length were assessed via peripheral quantitative computed tomography (pQCT) (XCT 3000, Stratec Medizintechnik GmBH, Pforzheim, Germany).

### 2.10 Collection of muscle tissue

Participants abstained from caffeine, alcohol and vigorous activity for 48 hours prior to the collection of muscle biopsies. The night before, participants consumed a low-protein, standardised meal of pasta and tomato-based sauce as previously described (Lamon *et al*., 2021). Portion size and water consumption were *ad libitum*. Participants recorded the portion size and water consumption from the pre-training trial and replicated this for the post-training trial.

Participants arrived at the testing facility at 0700 h after an overnight fast from 2100 h the previous evening. A muscle biopsy of the *vastus lateralis* was performed via a percutaneous needle biopsy technique modified to include suction (Bergstrom, 1962). Briefly, the skin over the *vastus lateralis* was sterilised and the area anaesthetised with 1% Lidocaine without epinephrine. An incision was made through the skin and muscle fascia. A muscle sample of 150-300 mg in size (Russell *et al*., 2013) was immediately snap frozen in liquid N_2_-cooled isopentane and stored in liquid N_2_ until required.

### 2.11 Training programs

After all the baseline measures were assessed, the 12-week resistance training program commenced. Every Monday, Wednesday and Friday, participants arrived at Deakin University between 0600-0800 h after an overnight fast from 2100 h the previous evening. Due to the various SARS-CoV-2-related lockdowns experienced throughout Victoria, Australia in 2020 and 2021 (Dunstan, 2021), there was a requirement for a sub-cohort of participants (*n=*11) to undertake a portion of their training sessions online. These training sessions were delivered via videoconferencing, replicating the time and days of the gym-based training sessions. Briefly, the gym-based training program consisted of squats, leg extensions, hamstring curls, shoulder press, biceps curls and seated row exercises. Participants were provided with weights and performed 3 sets of 8-10 repetitions at 60-80% 1RM. Supplementary Table 2 outlines the 2 different training programs undertaken by participants. Both programs were designed by an Exercise and Sports Science Australia-accredited exercise scientist and an Australian Strength and Conditioning Association-accredited strength and conditioning coach. Progressive overload (add 5% load) was applied to each exercise when an individual was able to complete 2 additional repetitions in the last set of an exercise in 2 consecutive sessions.

All participants were given a 25-g protein supplement (Ascent Protein, Denver, CO) either immediately before or after each training session to optimise the anabolic response to resistance training. The protein supplement was approved by Informed Choice (Informed Choice, 2021), thereby minimising the risk that the supplement contained any substances that are banned by the World Anti-Doping Agency (WADA).

### 2.12 Laboratory analysis

#### 2.12.1 Hormone analysis

Testosterone (sensitivity 0.18 ng·mL^-1^, intra-assay coefficient of variation (CV) 3.1-5.4%, inter-assay CV 4.2-7.4%), sex hormone binding globulin (SHBG; sensitivity 0.23 nmol·L^-1^, intra-assay CV 2.3-4.8%, inter-assay CV 5.2-6.3%), dehydroepiandrosterone (DHEA; sensitivity 0.03 ng·mL^-1^, intra-assay CV 3.9-7.6%, inter-assay CV 5.1-10.4%) and 5α-dihydrotestosterone (DHT; sensitivity 7.23 pg·mL^-1^, intra-assay CV 3.33-6.25%, inter-assay CV 6.49-7.47%) were measured via Enzyme-Linked Immunosorbant Assay (ELISA; #IBRE52151, #IB30176808, #IBRE52221, #IBDB5202, Abacus Dx, Parkville, Australia), according to manufacturer’s instructions.

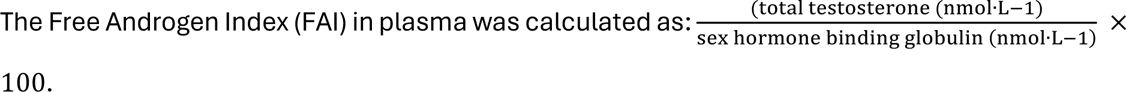

The full steroid profile in urine (including testosterone, its precursors and metabolites) was measured via gas chromatography mass spectrometry (GC/MS^n^) in a WADA-accredited laboratory as described previously (Salamin *et al*., 2022). Liquid chromatography mass spectrometry (LC/MS) was used to exclude confounding factors having a potential impact on endogenous testosterone production (e.g., alcohol, ketoconazole, aromatase inhibitors), while also ensuring that the participants were not using exogenous testosterone. All hormonal markers from urine were corrected for the specific gravity of urine, using the equation:

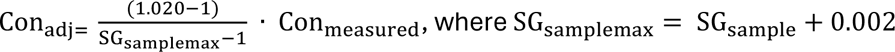

For validation of menstrual phase, oestradiol (E2; inter-assay CV 5.5-10.7%) and progesterone (P; inter-assay CV 6.4-19.2%) were measured via a competitive binding immune-enzymatic assay according to manufacturer’s instructions (Beckman Coulter, Lane Cove, Australia). Luteinising hormone (LH) was analysed via a sequential 2-step immune-enzymatic assay (inter-assay CV 5.2-7.8%) according to manufacturer’s instructions (Beckman Coulter) and follicle stimulating hormone (FSH) was analysed via a microparticle enzyme immunoassay (MEIA) (inter-assay CV 3.8-4.3%) according to manufacturer’s instructions (Beckman Coulter).

#### 2.12.2 Protein extraction

Protein was extracted from 20-25 mg of skeletal muscle tissue via manual homogenisation in 15 µL·mg^-1^ muscle 1× RIPA lysis buffer 1 (#J62524-AE, Thermo Fisher Scientific) containing 10 µL·mL^-1^ phosphatase inhibitor and 1 µL·mL^-1^ protease inhibitor cocktail (#78440, Thermo Fisher Scientific). The homogenised protein lysates were then gently spun at 4°C for 1 hour. After 1 hour, samples were centrifuged at 13,000 g for 15 min at 4°C. The protein concentration of each sample was determined via Pierce Bicinchoninic Acid (BCA) assay (#23225, Thermo Fisher Scientific) according to manufacturer’s instructions. Absorbance of samples was read at 562 nm using bovine serum albumin as a standard.

#### 2.12.3 Western blotting

The total protein and phospho-protein levels of the AR as well as markers of skeletal muscle protein synthesis were analysed via western blot. Following protein extraction, the samples were denatured for 5 min at 95°C with 1× Nupage sample reducing agent (#NP0004, Thermo Fisher Scientific) and 1× Nupage LDS sample buffer (#NP0007, Thermo Fisher Scientific). Thirty µg of total protein from each sample was loaded into a 4–15% gradient Criterion Tris-Glycine extended (TGX) Stain Free gel (#5678085, Bio-Rad, Gladesville, Australia), separated via electrophoresis at 200 V, 40 min and visualised on a Universal Hood II GelDoc (BioRad). The separated proteins were transferred to an Immobilon PVDF-FL membrane (#IPFL00005, Millipore, Billerica, MA) at 100 V, 60 min and blocked for 1 hour in 5% skim milk in Tris buffered saline plus 0.1% Tween-20 (TBST; #P1379, Sigma-Aldrich, North Ryde, Australia). Membranes were incubated at 4°C overnight in the primary antibody. The antibodies and conditions used were for Akt (#2920, 1:1000, mouse, Cell Signalling Technologies, Danvers, MA), p-Akt^ser473^ (#4060, 1:1000, rabbit, Cell Signalling Technologies), AR (#5153, 1:500, rabbit, Cell Signalling Technologies), p-AR^ser213^ (#PA537478, 1:500, rabbit, Thermo Fisher Scientific), p-AR^ser650^ (#537479, 1:500, rabbit, Thermo Fisher Scientific), MAPK (#4696, 1:1000, mouse, Cell Signalling Technologies), p-MAPK^thr202/tyr204^ (#9101, 1:500, rabbit, Cell Signalling Technologies), mTOR (#4517, 1:1000, mouse, Cell Signalling Technologies), p-mTOR^ser2448^ (#5536, 1:1000, rabbit, Cell Signalling Technologies), MuRF-1 (#MP3401, 1:1000, rabbit, ECM Biosciences), 4E-BP1 (#9452, 1:1000, rabbit, Cell Signalling Technologies), p-4E-BP1^thr37/46^ (#2855, 1:500, rabbit, Cell Signalling Technologies), rpS6 (#2217, 1:1000, rabbit, Cell Signalling Technologies) and p-rpS6^ser235/236^ (#4856, 1:1000, rabbit, Cell Signalling Technologies). The membranes were washed thrice for 10 min in TBST and incubated with the corresponding secondary antibody (#5151, 1:10,000, anti-rabbit IgG Dylight 800 or #5470, 1:10,000, anti-mouse IgG DyLight 680; Cell Signalling Technologies) for 1 hour at room temperature. Following 2× 10-min washes in TBST and 1× 10-min wash in 1× phosphate buffered saline (PBS), the proteins were exposed on an Odyssey® CLx Infrared Imaging System and individual protein band optical densities were determined using the Odyssey® Infrared Imaging System software (Image Studio V5.2, Licor Biosciences). All blots were normalized against total protein load using the Bio-Rad Image Lab software (v6.0).

#### 2.12.4 Immunohistochemical staining

Muscle fibre type composition and cross-sectional area were assessed via immunohistochemistry (IHC), staining for myosin heavy chain (MHC I and IIx) and laminin. Eight-micron cross sections of the muscle samples were cut on a microtome cryostat and loaded onto glass slides. The muscle sections were blocked in 10% goat serum (#16210072, Thermo Fisher Scientific) in 1× PBS for 1 hour at room temperature. The muscle sections were incubated for 1 hour at room temperature in a primary antibody cocktail containing antibodies specific to anti-MHCI (#BA-F8, 1:20, Developmental Studies Hybridoma Bank; DSHB), anti-MHCIIx (#6H1, 1:20, DSHB) and anti-laminin (#L9393, 1:100, Sigma Aldrich) in 10% goat serum/PBS. The muscle sections were washed thrice in PBS and then incubated in a secondary antibody cocktail containing goat anti-mouse IgG2b Alexa Fluor 647 (#A-21242, 1:500, Thermo Fisher Scientific), goat anti-mouse IgM Alexa Fluor 555 (#A-21426, 1:500, Thermo Fisher Scientific) and Goat anti-rabbit IgG Alexa Fluor 405 (#A-31556, 1:500, Thermo Fisher Scientific) in 10% goat serum/PBS for 1 hour at room temperature. The muscle sections were washed thrice in PBS and then mounted using Vectashield fluorescent mounting medium (#H-1900, Vector Laboratories, Abacus Dx). One image of the entire muscle section was visualised using a Fluoview fv0i confocal microscope (Olympus) at 10× magnification and analysed using Semi-automatic Muscle Analysis using Segmentation of Histology (SMASH) software (MATLAB application, Mathworks, USA) (Schneider *et al*., 2012). The average number of myofibres per section was 753.8 ± 370.6.

*In vivo* AR localisation was also assessed via IHC. Eight-micron muscle sections were thawed for 10 minutes before they were fixed in 4% paraformaldehyde (PFA) for 10 min. The muscle sections were washed thrice in PBS and permeabilised in 0.1% Triton-X 100 for 5 min. After permeabilization, the muscle sections were blocked for 1 hour at room temperature in 5% bovine serum albumin/PBS. Following blocking, the muscle sections were incubated in an antibody against the androgen receptor (#5153, 1:50, Cell Signalling Technologies) in blocking buffer at 4°C overnight. The following day the sections were washed and incubated in a secondary antibody cocktail containing goat anti-rabbit IgG Alexa Fluor 488 (#A-11008, 1:500, Thermo Fisher Scientific) and wheat germ agglutinin (#W32466, 1:1000, Thermo Fisher Scientific) in blocking buffer for 1 hour at room temperature. The cells were washed thrice in PBS and stained with 0.1 µg·mL^-1^ DAPI stain (#62248, 1:1000, Thermo Fisher Scientific) in PBS for 10 minutes and mounted using Vectashield fluorescent mounting medium (#H-1900, Abacus Dx). Ten images of each muscle section, with an average of 29.9 ± 8.6 fibres per image were obtained with dedicated software at 40× magnification (Eclipse Ti2, Nikon, Tokyo, Japan).

#### 2.12.5 RNA extraction and quantification

Frozen skeletal muscle (∼15 mg) was combined with lysis buffer (#1053393, Qiagen, Clayton, Australia) and homogenised with 650-800 mg silica beads for 2× 30 second homogenisation steps at 6500 rpm (MagNA lyser, Roche Diagnostics, North Ryde, Australia). RNA was extracted from the homogenised lysate using an Allprep DNA/RNA/miRNA Universal extraction kit (#80224, Qiagen) according to manufacturer’s instructions, including a proteinase K and DNase treatment.

The quality and quantity of the RNA extracted was assessed using the TapeStation System according to manufacturer’s instructions (Agilent Technologies, Mulgrave, Australia). An RNA integrity number (RIN) of >7 was considered acceptable for downstream analysis. The average sample yield was 73.1 ng·μL^-1^ ± 22.5 ng·μL^-1^ and the RIN average was 8.1 ± 0.8.

#### 2.12.6 RNAseq

The RNAseq libraries were prepared using the Illumina TruSeq Stranded Total RNA with Ribo-Zero Gold protocol and sequenced with 150-bp paired-end reads on the Illumina Novaseq6000 (Macrogen Oceania Platform). Reads underwent quality check with FastQC (v0.11.9); Kallisto (v0.46.1) was used to map reads to the human reference genome (*HomoSapien GRCh38)* and to generate transcript counts. Genes with an average across all samples of 10 reads per million (RPM) or less reads were removed (63%) from further analysis leaving a total of 14,979 for analysis. All RNA sequencing data generated or analysed during this study are included in this published article, its supplementary information files and publicly available repositories (GEO: link pending). The R code used for the analysis is available at https://github.com/DaniHiam/TESTO_RNAseq

### 2.13 In vitro experiments

#### 2.13.1 Isolation of primary myocytes

To test whether there is a causal association between testosterone treatment and the Akt/mTOR pathway, we isolated myocytes from a sub-cohort of participants (*n=*6) that underwent the 12-week resistance training program. To ensure a heterogenous sample, we selected participants from across a range of responses to the resistance training program. A third biopsy was collected at rest from this subset of participants from the *vastus lateralis* in exactly the same manner as described above approximately 6 months after the second muscle biopsy. Approximately 100-200 mg of muscle was placed in ice-cold serum free Hams F10 nutrient mixture (#11550043, Thermo Fisher Scientific). The muscle was washed thrice in ice-cold serum-free Hams F10 nutrient mixture. The tissue was minced manually and resuspended in serum-free Hams F10 media and centrifuged at room temperature for 5 min, 230 *g*. The supernatant was removed, and the resulting pellet was resuspended in warm (37°C) 0.05% Trypsin/EDTA (#25300062, Thermo Fisher Scientific) and dissociated thrice on an orbital shaker for 20 min at 37°C. After the final dissociation, 10% v/v horse serum (#16050122, Thermo Fisher Scientific) was added to the tissue slurry and cells were filtered through a 75 µm cell strainer. The resultant flow-through was centrifuged at room temperature for 5 min, 530 *g* and the resulting pellet resuspended in proliferation media containing Hams F10 nutrient mixture, 20% foetal bovine serum (FBS; #10099141, Thermo Fisher Scientific), 1% penicillin-streptomycin (#15140122, Thermo Fisher Scientific), 0.5% Amphotericin B (#15290018, Thermo Fisher Scientific) and 25 µg·mL^-1^ fibroblast growth factor (fGFb; #PHG0026, Thermo Fisher Scientific). The cells were plated in flasks pre-coated with an extracellular matrix (ECM) gel from Engelbreth-Holm-Swarm murine sarcoma (#E1270, Sigma-Aldrich). The cells were maintained in humidified air at 37°C, 5% CO_2_. Proliferation media was changed every 48 h and the cells passaged once they had reached 70-80% confluence.

#### 2.13.2 Purification of cultured human myoblasts

Once the cells had reached 70-80% confluence, myogenic satellite cells were purified using Magnetic Activated Cell Sorting (MACS) with anti-CD56+ microbeads (#130-050-401, Miltenyi Biotec, Bergisch Gladbach, Germany) as previously described (Agley *et al*., 2013; McIlvenna *et al*., 2021).

#### 2.13.3 Differentiation of cultured human myoblasts

Once the enriched myogenic cells reached 60-70% confluence, differentiation was induced by replacing the proliferation medium with Dulbecco’s Modified Eagle’s Medium (DMEM) without phenol red (#21063029, Thermo Fisher Scientific) supplemented with 2% horse serum (#16050122, Thermo Fisher Scientific) and 1% penicillin-streptomycin (#15140122, Thermo Fisher Scientific). DMEM without phenol red was used to eliminate the oestrogenic effects of phenol red (Estrada *et al*., 2003; Wannenes *et al*., 2008; Eriksen *et al*., 2014). Cells were differentiated either in the presence of 100 nM testosterone dissolved in ethanol (#DRE-C17322500, Novachem, Heidelberg West, Australia) (henceforth referred to as testosterone treated, or TT) or the equivalent volume (2 µL·well^-1^) of the vehicle control, ethanol (henceforth referred to as control, or CON).

The cells were harvested for protein extraction at baseline (Day 0) and after 1 (D1), 4 (D4) and 7 days (D7) of differentiation, using 150 µL·well^-1^ 1× RIPA lysis buffer (#J62524-AE, Thermo Fisher Scientific) containing 10 µL·mL^-1^ phosphatase inhibitor and 1 µL·mL^-1^ protease inhibitor cocktail (#78440, Thermo Fisher Scientific). The homogenised protein lysates were then gently spun at 4°C for 1 hour. Samples were centrifuged at 13 000 *g* for 15 min at 4°C and the protein concentration of each sample determined via Pierce Bicinchoninic Acid (BCA) assay (#23225, Thermo Fisher Scientific) according to manufacturer’s instructions. Western blots were completed under the same conditions as described above on the resultant cell protein lysate, loading 10 µg total protein·well^-1^.

#### 2.13.4 Immunohistochemical analysis of AR location

In addition to western blot analyses, we also investigated the effect of testosterone treatment on the localisation of the AR in myocytes. At baseline, and after 1, 4 and 7 days of differentiation, human primary myocytes were washed thrice for 5 min in PBS. The myocytes were fixed with 2% paraformaldehyde (PFA)/PBS for 10 min and washed thrice for 5 min in PBS. The fixed cells were permeabilised in 0.1% Triton X-100/PBS for 5 min and blocked with 3% BSA/PBS for 40 min at room temperature. Cells were incubated for 1 hour at room temperature with the primary antibody against total androgen receptor (#5153,1:50 in blocking buffer, rabbit, Cell Signalling Technologies). Following this, the cells were incubated with AlexaFluor 488 goat anti-rabbit IgG (#A-11008, 1:5000, Thermo Fisher Scientific) and AlexaFluor 647 phalloidin (#A22287, 1:200, Thermo Fisher Scientific) in 1% BSA/PBS for 1 hour at room temperature. The cells were washed thrice in PBS and stained with 0.1 µg·mL^-1^ DAPI (#62248, 1:1000, Thermo Fisher Scientific) stain in PBS for 10 minutes. Cell images were obtained with dedicated software at 100× magnification *(Eclipse Ti2, Nikon, Tokyo, Japan)*.

#### 2.13.5 Quantification of androgen receptor intensity

Quantification of androgen receptor content was performed using the open-source image analysis software *CellProfiler* (version 4.2.5) and an analysis pipeline developed within this study (Stirling *et al*., 2021). For *in vivo* analysis, 5-10 images per participant for each time point were captured at 40× magnification, imported into CellProfiler and split into individual grayscale images of sarcolemma, nuclei and AR staining based on RGB channels. DAPI-stained myonuclei were identified as objects with a diameter range between 8 and 50 pixels, as previously described (Sanz *et al*., 2019). To quantify the nAR/AR ratio, a binary mask of the DAPI-stained myonuclei was applied to the AR-stained images and the total intensity of AR expression within the nuclei was expressed as a ratio of total AR intensity per field. The percentage of nuclei positive for AR expression was quantified by counting the number of DAPI-stained myonuclei encompassed within a binary mask of the AR-stained images. The percentage of nuclei highly expressing AR was determined by creating a binary mask of regions expressing 3x the mean intensity of AR staining per field from the AR image and identifying the percentage of nuclei within this mask. The total number of muscle fibres per field was quantified using the previously described Muscle2View CellProfiler pipeline (Sanz *et al*., 2019). Each binary mask was manually reviewed by 2 independent analysts, and any masks containing apparent visual artifacts were excluded from analysis (an average of 3.8 images per participant were excluded).

For *in vitro* analysis, images of 1 field per well from 6 wells were imported into CellProfiler and split into individual grayscale images of actin, nuclei and AR staining based on RGB channels. A binary mask of the myocytes was created using the actin images, and the total area within each field occupied by this mask quantified. The total intensity of the androgen receptor stain was measured and expressed relative to the area occupied by myocytes per each field, as the measure “AR intensity”. This CellProfiler pipeline was validated for the accurate and complete identification of both myocytes and androgen receptor protein through a manual review of 20% of the total images, performed by 2 independent analysts.

### 2.14 Statistical analysis

The statistical analyses for this study were performed using GraphPad Prism version 8 *(GraphPad Software, La Jolla, CA)* and R software version 4.0.2 using the packages *lmerTest* (Kuznetsova *et al*., 2017), *tidyverse* (Hadley Wickham *et al*., 2019), *car* (Fox and Weisberg, 2019), *AICcmodavg* (Mazerolle, 2020).

Two-tailed, paired t-tests in GraphPad were used to assess the effect of a 12-week resistance training program on the changes to participant anthropometric data, the protein expression of all measured proteins and thigh muscle size, strength and power. Leg press strength and muscle power were expressed as values relative to the individual’s total body lean mass (kg) and the changes to muscle size, strength and power were expressed as delta percent change (Δ%). The phosphorylation status of all proteins is expressed as the amount of phosphorylation relative to the total protein content of that protein (e.g., p-AR^ser213^= p-AR^ser213^/total AR content) and the changes in protein content from pre- to post-training is expressed as fold-change from the pre-training levels. If the protein content of a given protein did not change, the average of the pre- and post-training values was used for all post-training (delta change) linear models.

One-way, repeated measures analyses of variance (ANOVA) were used in GraphPad to assess the effect of 12 weeks of resistance training on the concentrations of testosterone, SHBG, DHT, DHEA, E2 and P and changes in dietary protein intake and total energy expenditure. If hormone concentrations did not significantly change, the area under the curve (AUC) using the trapezoidal method was used for all post-training (delta change) linear models. The AUC provides a surrogate measure for the total amount of hormone that participants were exposed to across the entire 12 weeks.

Linear models were used in Rstudio to examine whether the outcome (muscle size, strength and power at baseline, or the delta change in these variables) was influenced by the independent variables of serum testosterone concentrations, the FAI, DHT, DHEA, the protein expression of AR or p-AR, the nAR/AR ratio, or the proportion of AR+ nuclei. The model was of the form: *outcome* = *independent variable* + *covariate* (*if applicable*). Before further analyses, the normality of all variables was assessed, and variables were log-transformed if necessary.

Possible covariates included age, BMI, E2, P, luteinising hormone, follicle stimulating hormone, chronotype, hormonal contraceptive use, average protein intake, average daily physical activity and menstrual phase. Before fitting the linear models, Akaike Information Criterion (AIC) tests were run on linear models containing all possible combinations of the covariates to establish which covariates were required in the final model. The model with the lowest AIC that explained the largest proportion of variance in the association was chosen as the final model. The collinearity of linear models with appropriate covariates was assessed through variance inflation factors (VIF), with a threshold of 3 set. The homoscedasticity of each model was assessed through residual and QQ plots.

Transcriptomic data were analysed using Rstudio 4.1.3 (R Core Team, 2021). Differential gene analysis was conducted using the R package DeSeq2 (Love *et al*., 2014) using the model: Genes ∼ ID + timepoint. ID was used to account for repeated measurements. ChIP-X enrichment analysis 3 (ChEA3) (https://maayanlab.cloud/chea3/) was used to perform transcription factor (TF) enrichment analysis on the differentially expressed genes (Keenan *et al*., 2019). The mean rank integration method was used to calculate the ranking of the most enriched TFs.

We considered significant genes and transcription factors significant with an FDR adjusted p value <0.05. The following packages were also used in our analysis; *tidyverse* (Wickham *et al*., 2019), *superheat* (Barter & Yu, 2018), *biomaRt* (Durinck *et al*., 2009).

Graphpad software was used to performed two-way ANOVAs with multiple comparisons to assess the effect of 7 days of testosterone treatment on myocyte diameter and the effect of acute testosterone treatment on the protein content of the androgen receptor and markers of protein synthesis.

All values are presented and mean ± SD, unless otherwise stated. Significance for all statistical tests was set at *p*<0.05.

## 3. Results

Of the 35 females enrolled in the study, 27 females completed the 12-week resistance training program. Due to the SARS-CoV-2-related lockdowns experienced throughout Victoria, Australia in 2020 and 2021 (Dunstan, 2021), a sub-cohort of participants (*n*=11 of the 27 having completed the study) undertook a portion of their 36 training sessions at home (range: 2 to 6 sessions, average 4 sessions) delivered via videoconferencing. There were no differences in age, height, weight, BMI, ratio of hormonal contraceptive users or calculated baseline 1RM for any exercise between the participants who performed all of their training sessions in the gym and those that performed some sessions via video conferencing (Table 1). There was no between-group difference in the trajectory of working weight progression for any exercise (Supplementary Figure 2). There was also no between-group difference in training-induced changes in working weight for any exercise, thigh muscle cross sectional area, power (Supplementary Figure 3), or pre- or post-training hormone levels (Supplementary Figure 4). The 2 training regimes were therefore considered equivalent and the results from both cohorts were pooled for all further analyses.

**Table 1.** Baseline anthropometric and strength data separated for all participants that completed the gym-based training program (n=16) or participants who completed the blended gym- and home-based training program (n=11) and for all participants combined (n=27).

|  | Participants undergoing normal training (n=16) | Participants spending 2-6 sessions at home (n=11) | All participants (n=27) |
| --- | --- | --- | --- |
| **Age (years)** | 24.1 ± 5.0 | 21.9 ± 3.9<br>(p = 0.24) | 23.5 ± 4.8 |
| **Height (m)** | 1.7 ± 0.1 | 1.6 ± 0.1<br>(p = 0.62) | 1.7 ± 0.1 |
| **Weight (kg)** | 65.8 ± 13.1 | 66.3 ± 11.1<br>(p = 0.92) | 65.9 ± 12.4 |
| **BMI (kg·m<sup>-2</sup>)** | 23.3 ± 3.7 | 24.5 ± 4.1<br>(p = 0.42) | 23.6 ± 3.8 |
| **Hormonal contraceptive use** | Users: 46%<br>Non-users: 54% | Users: 67%<br>Non-users: 33%<br>(p = 0.30) | Users: 51%<br>Non-users: 49% |
| **Baseline testosterone concentrations (nmol·L<sup>-1</sup>)** | 2.1 ± 0.5 | 2.0 ± 0.6<br>(p = 0.23) | 2.0 ± 0.6 |
| **Calculated squat 1RM (AU)** | 104.7 ± 31.4 | 105.4 ± 24.3<br>(p = 0.97) | 104.8 ± 30.0 |
| **Calculated leg press 1RM (AU)** | 179.8 ± 42.3 | 180.1 ± 27.5<br>(p = 0.89) | 179.3 ± 39.1 |
| **Calculated leg extension 1RM (AU)** | 52.8 ± 12.2 | 45.9 ± 13.0<br>(p = 0.15) | 51.0 ± 12.9 |
| **Calculated hamstring curl 1RM (AU)** | 35.9 ± 6.4 | 36.9 ± 6.9<br>(p = 0.74) | 36.1 ± 6.7 |
| **Calculated seated row 1RM (kg)** | 40.4 ± 7.5 | 38.4 ± 7.3<br>(*p* = 0.42) | 39.7 ± 7.5 |
| **Calculated shoulder press 1RM (kg)** | 9.4 ± 2.9 | 8.7 ± 2.4<br>(*p* = 0.75) | 9.3 ± 2.8 |
| **Calculated biceps curl 1RM (kg)** | 8.2 ± 1.4 | 8.4 ± 1.4<br>(*p* = 0.97) | 8.2 ± 1.4 |

Values are mean ± standard deviation. *p*-values shown in brackets are for pairwise comparisons to participants having completed normal training using unpaired *t*-tests.

### 3.1 Effect of 12 weeks of resistance training on body composition and muscle mass and function

Twelve weeks of resistance training increased body mass by 1.4% (pre-training: 65.4 ± 10.9 kg, post-training: 66.2 ± 10.5 kg, *p*<0.05). Total body lean mass increased by 1.9% (pre-training: 42.1 ± 5.5 kg, post-training: 42.9 ± 5.7 kg, *p*<0.01). Total body fat mass remained unchanged (pre-training 21.0 ± 7.8 kg, post-training: 21.5 ± 7.7 kg, *p*=0.125).

Muscle strength (measured via leg press 1RM) increased by 27.3% (*p<*0.001), thigh muscle cross-sectional area (CSA; measured via pQCT) by 5.9% (*p<*0.001) and muscle power (measured via vertical jump) by 13.0% (*p*<0.05) (Figure 1A-C).

**Figure 1.**
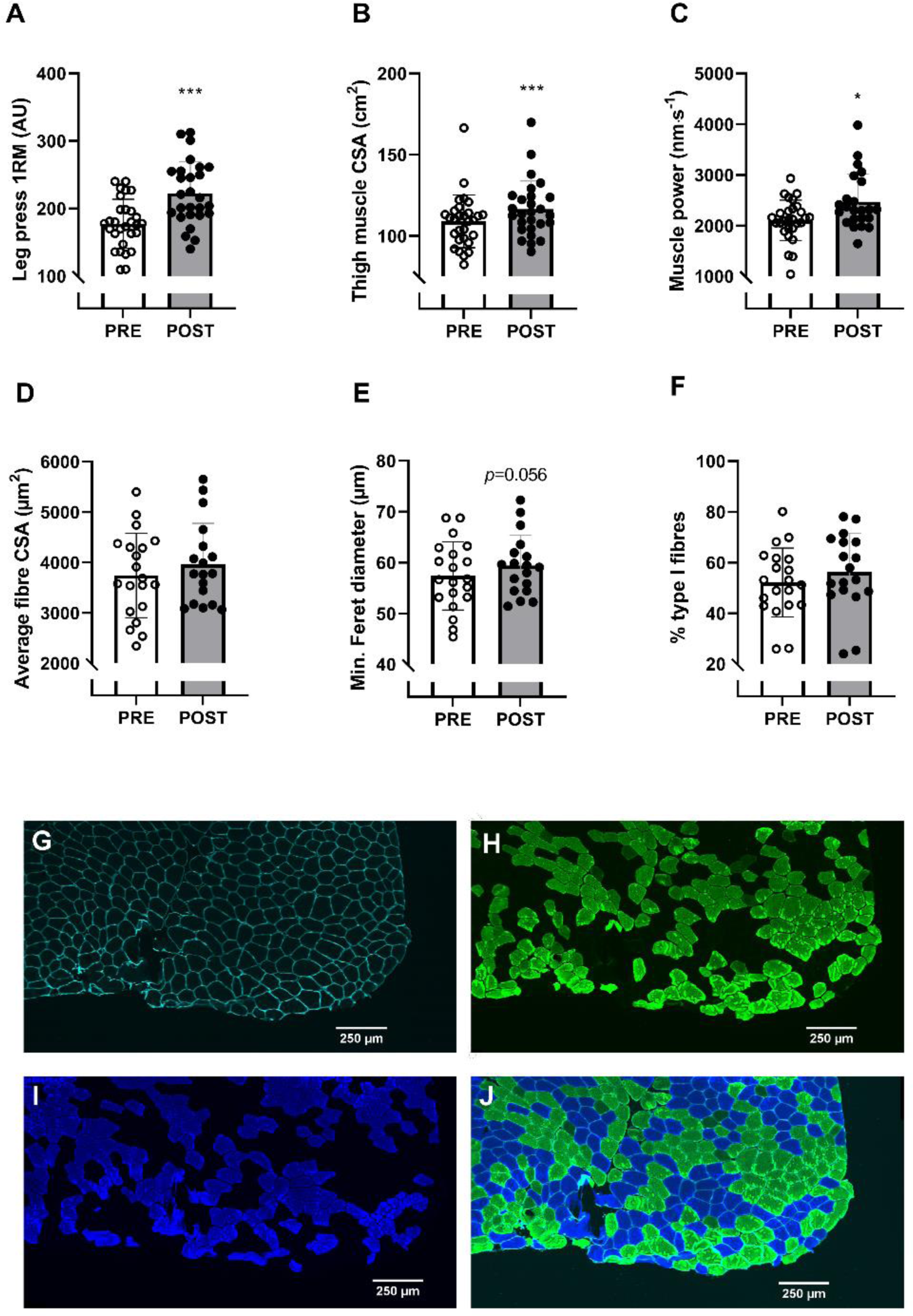
A) Leg press 1RM, B) (B) thigh muscle cross sectional area (CSA) (C) muscle power D) mixed muscle myofibre CSA, E) minimum Feret diameter and F) percentage of type I fibres in untrained, pre-menopausal females before and ager 12 weeks of resistance training (n=27). G) Representative images showing the laminin-stained sarcolemma of a muscle section, H) type I fibres, I) type II fibres and J) a composite image of both. Scale for representative images is 1.24 µm·pixel^-1^ for all images shown. n=753.8 ± 370.6 fibres per slice. White scale bar represents 250 µm. Pre-training (PRE) values are indicated by clear bars and post-training (POST) values are indicated by dark bars. *indicates p<0.05, ***indicates p<0.001. Data were analysed using two-tailed, paired t-tests. Values are represented as mean ± SD.

### 3.2 Effect of 12 weeks of resistance training on the cellular markers of muscle hypertrophy

The average myofibre CSA (µm^2^) did not change with resistance training (mixed fibre increase 7.7%, *p*=0.100; type I fibre CSA increase: 8.7%, *p*=0.131; type II fibre CSA increase: 5.7%, *p*=0.269). Similarly, the minimum Feret diameter, which is indicative of the circularity of the fibre and robust against experimental errors such as the orientation of the fibre upon cutting and therefore a more reliable indication of fibre CSA, increased by 4% but did not reach statistical significance (*p*=0.056). The percentage of type I and type II fibres did not change with training (Figure 1D-F) (*p*=0.725). Representative images are shown in Figure 1G-J.

### 3.3 Effect of 12 weeks of resistance training on androgen receptor protein content, phosphorylation status and nuclear localisation

Twelve weeks of resistance training did not induce any change in the protein (*p*=0.672) content of the androgen receptor (Figure 2A), or its phosphorylation status at either serine residue 213 (p-AR^ser213^, *p*=0.730; Figure 2B) or serine residue 650 (p-AR^ser650^, *p*=0.750; Figure 2C). *AR* mRNA expression also did not change with 12 weeks of resistance training (FDR=0.956; Figure 2D). Similarly, 12 weeks of resistance training did not induce any change in nAR *in vivo*. This was shown in both the ratio of nAR to total AR stain (nAR/AR ratio; Figure 2E) and the percentage of nuclei that were AR positive (%AR+; Figure 2F).

**Figure 2.**
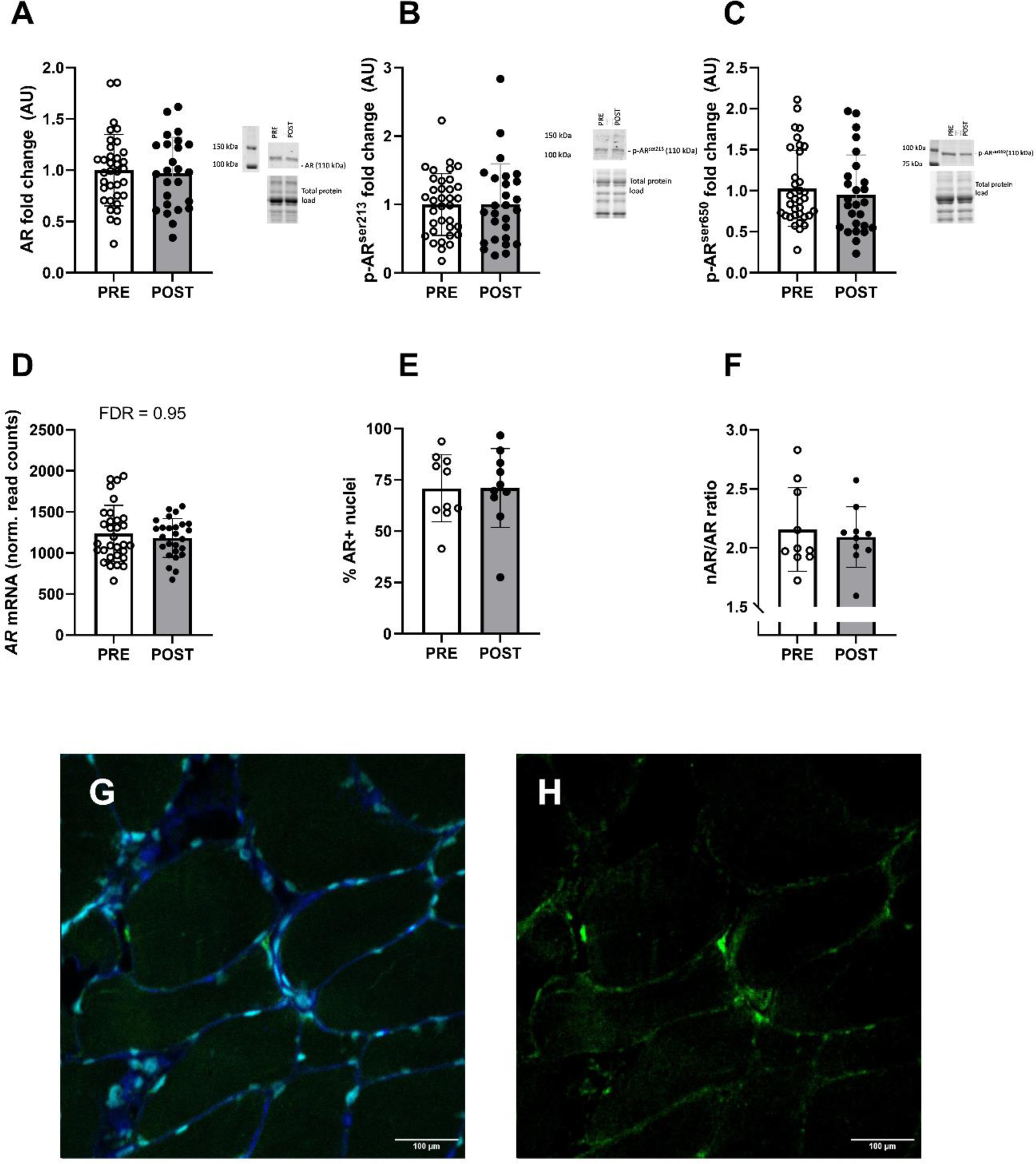
The protein content of A) total AR protein, B) p-AR^ser213^, C) p-AR^ser650^, D) AR mRNA expression (n=27), E) the percentage of AR+ nuclei or F) the nAR/AR ratio (n=10 participants) before and after 12 weeks of resistance training in previously untrained, pre-menopausal females. Western blots for each protein are resented beside the corresponding graph. Pre-training (PRE) values are indicated by clear bars and post-training (POST) values are indicated by dark bars. G) Representative composite image of a muscle section stained with DAPI (cyan; stains the nucleus), wheat germ agglutinin (blue; stains the sarcolemma) and α-AR (green) at 40x magnification. H) Representative image of α-AR (green) stain at 40x magnification. Scale for representative images is 0.62 μm·pixel^-1^. White scale bar represents 100 μm. n=5-10 images per section, average 29.9 ± 8.6 fibres per image. Data were analysed via two-way, paired t-tests. Values are represented as mean ± SD.

### 3.4 Effect of 12 weeks of resistance training on molecular markers of protein synthesis and degradation

Twelve weeks of resistance training increased the levels of muscle protein synthesis signalling molecules total Akt protein by 13% (*p*<0.05) (Figure 3A). There was no change in the basal total or phospho-protein content of other markers of protein synthesis or degradation p-Akt^ser473^ (*p*=0.972), mTORC1 (*p*=0.365), p-mTORC1^ser2448^ (*p*=0.976), MAPK (*p*=0.330), p-MAPK^thr202/tyr204^ (*p*=0.191), rpS6 (*p*=0.890), p-rpS6^ser235/236^ (*p*=0.533), 4E-PB1 (*p*=0.830), p-4E-PB1^thr37/46^ (*p*=0.977), and MuRF1 (TRIM63) (*p*=0.209) (Supplementary Figure 5).

**Figure 3.**
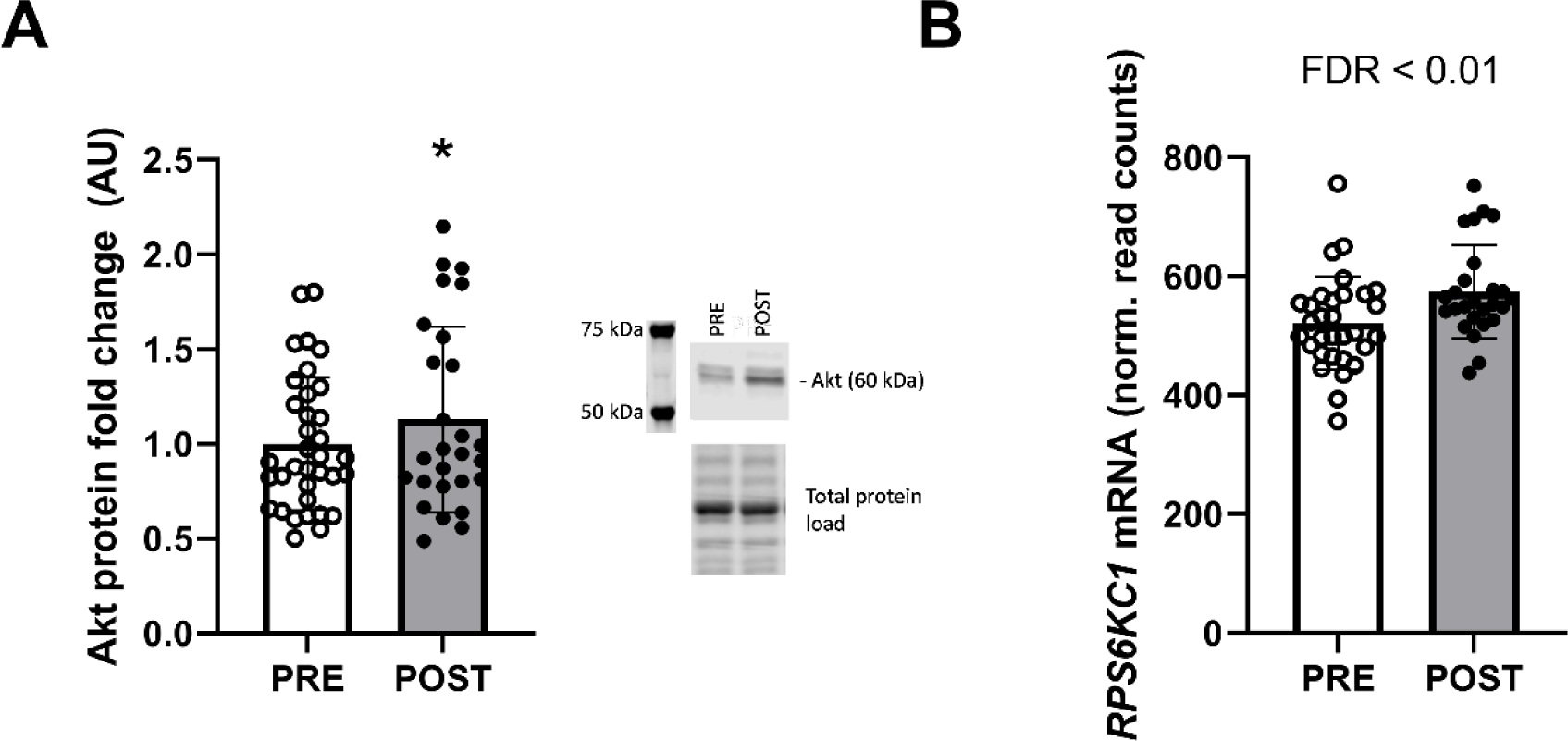
(A) total Akt protein and (B) RPS6Kc1 mRNA before and after 12 weeks of resistance training in pre-menopausal, previously untrained females (n=27). Representative Western blots for total Akt are presented beside the corresponding graph. Pre-training (PRE) values are indicated by clear bars and post-training (POST) values are indicated by dark bars. Values are represented as mean ± SD.

mRNA expression of *RPS6KC1* mRNA increased by 11% (FDR<0.01; Figure 3B). There was no change in the mRNA levels of protein degradation markers *TRIM63* (FDR*=*0.688), *FBXO32* (FDR=0.757), *TRAF6* (FDR=0.443), *FOXO1* (FDR=0.430) or *FOXO3* (FDR=0.268) (Supplementary Figure 5).

### 3.5 Effect of 12 weeks of resistance training on plasma and urine sex hormone concentrations

The average plasma testosterone concentration at baseline was 2.0 ± 0.6 nmol·L^-1^, ranging between 1.1 and 3.1 nmol·L^-1^. Twelve weeks of resistance training did not change the plasma levels of testosterone, DHT, DHEA or the FAI (Figure 4A-D). The androgen profile from urine measured via LC-MS, which included testosterone, epitestosterone, androsterone, etiocholanolone, 5α-adiol, 5β-adiol, DHEA and DHT, did not change across 12 weeks of resistance training and confirmed what was observed in plasma (Supplementary Figure 6). As circulating hormones directly reflect the form that is utilised by the muscle, plasma hormone concentrations were used over urine concentrations in all further analyses. Since there were no training-induced changes at any time point, the AUC of these hormones, which is indicative of the total exposure to this hormone during the 12-week training period, was used in subsequent analyses, where stated.

**Figure 4.**
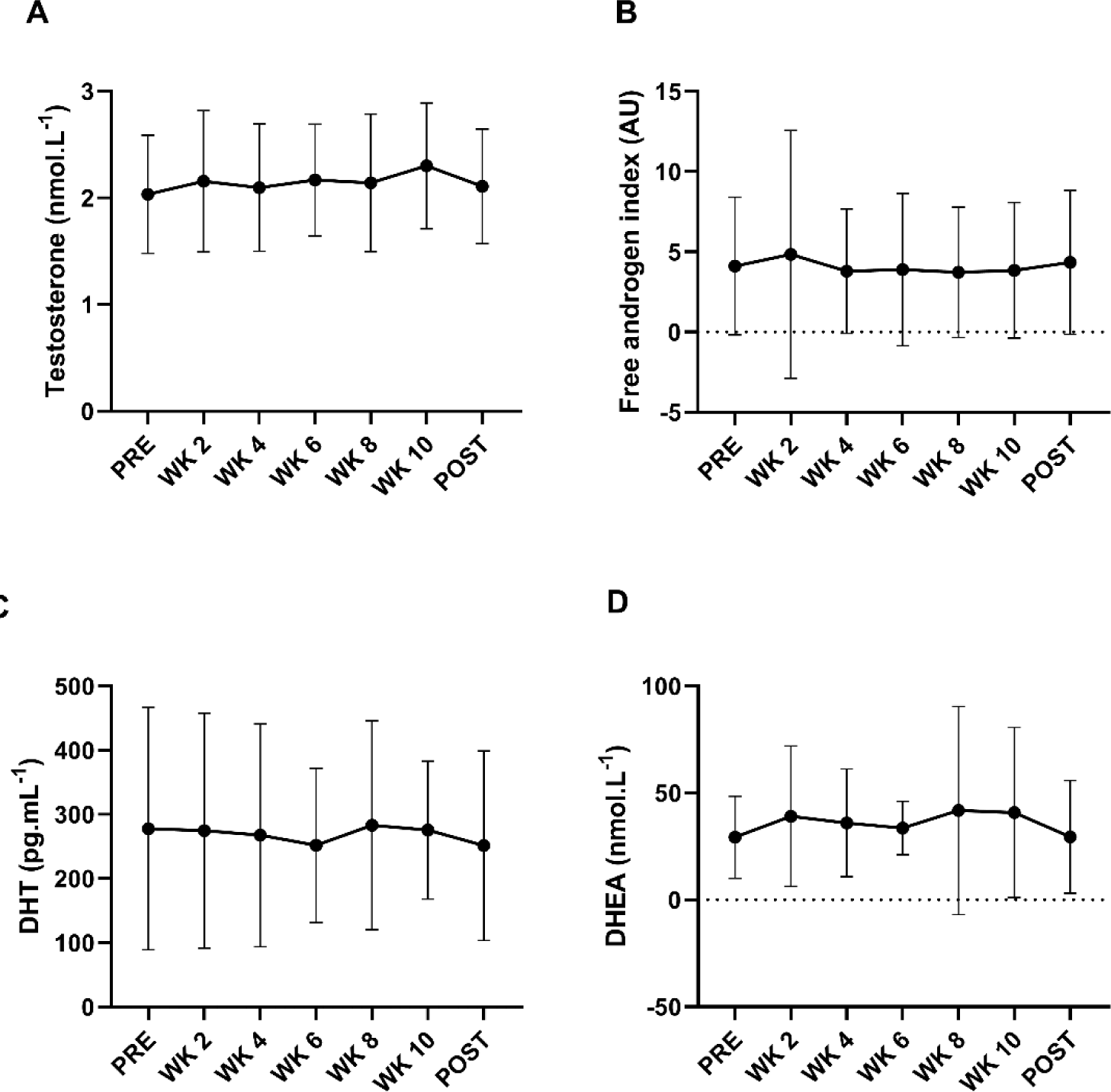
Twelve weeks of resistance training did not affect concentrations of (A) total testosterone (nmol·L^-1^), (B) the free androgen index (FAI; AU), (C) dihydrotestosterone (DHT; pg·mL^-1^) or (D) dehydroepiandrosterone (DHEA; nmol·L^-1^) in plasma of previously untrained, pre-menopausal females (n=27). Data were analysed using a one-way ANOVA. Values are represented as mean ± SD.

### 3.6 Effect of 12 weeks of resistance training on the muscle transcriptomic profile

Two hundred and fourteen transcripts were differentially expressed between pre- and post-training (122 up-regulated, 92 down-regulated, *FDR*<0.05) (Figure 5A). We then investigated the putative role of androgens in the regulation of the muscle transcriptome, by conducting transcription factor enrichment (TF) analysis on the differentially expressed genes using the mean rank integration method to rank the most enriched TFs. The top-15 TFs included muscle-specific transcription factors MYOG, MYOD1 and MEOX2 (Figure 5B). The AR was ranked 284 of 1632 TFs ranked, indicating that the androgen receptor and its binding to androgen response elements (ARE) may only play a minor role, if any, in the female global muscle transcriptomic response to anabolic stimulation. In line with this finding, none of the 14,979 individual detected transcripts were significantly associated with total testosterone concentrations.

**Figure 5.**
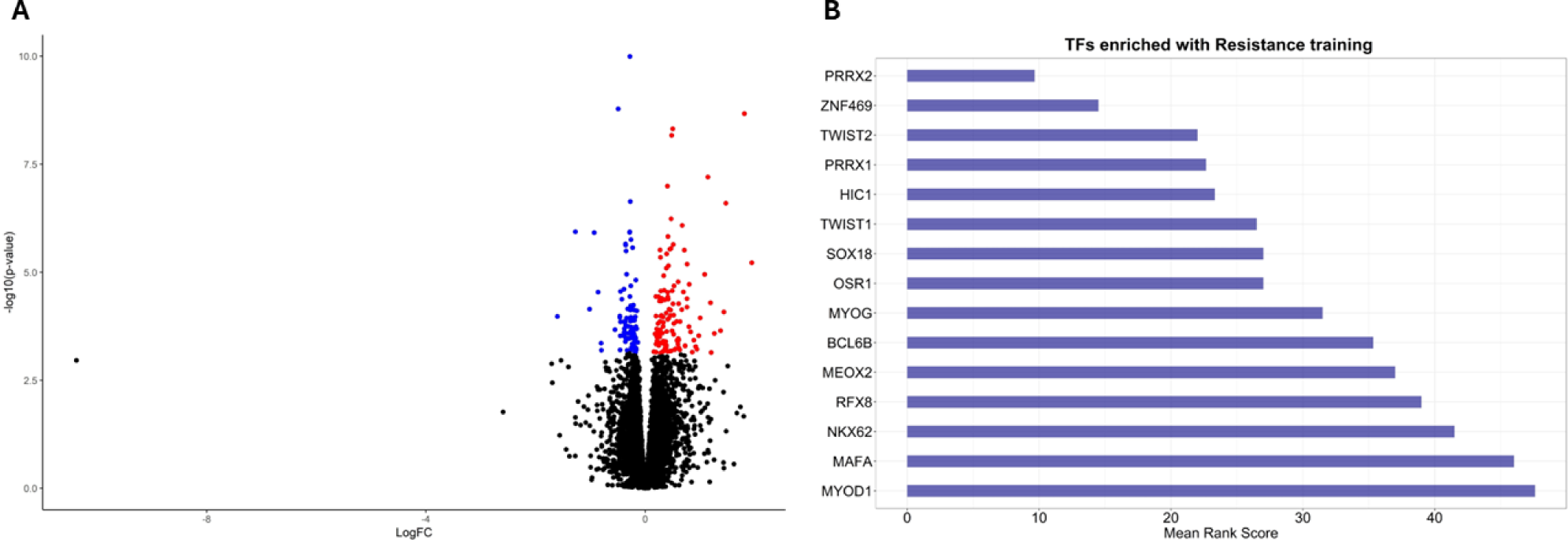
(A) Volcano Plot displaying 122 up-regulated and 92 down-regulated transcripts in response to 12 weeks of resistance training in pre-menopausal females (n=35 pre-training, 27 post training). Each point represents a transcript. Red points indicate an increase in mRNA expression following exercise. Blue points indicate a decrease in mRNA expression following exercise. Black dots represent genes which were not significantly differentially expressed. Significance was set at a false discovery rate (FDR) adjusted p value <0.05. (B) Top-15 transcription factors regulating the differentially expressed genes were ranked according to ChIP-X enrichment analysis 3 (ChEA3) using the mean rank integration method. Significance was set at FDR adjusted p value <0.05.

### 3.7 Associations between androgen hormone concentrations and muscle strength, size and power pre- and post-training

We next used linear models to test the association between baseline androgen concentrations, or total exposure to androgens during the 12-week training period, and baseline or training-induced changes in muscle size and function, respectively. AIC tests were used to identify the moderators to be included in each model. There was no evidence of an association between baseline total testosterone and pre-training muscle strength (*p*=0.445), CSA (*p*=0.417), power (*p*=0.929) and fibre CSA (*p*=0.147). Similarly, there was no evidence of an association between the AUC of testosterone (indicative of the total exposure to testosterone across 12 weeks) and the training-induced changes in muscle CSA (*p*=0.969), strength (*p*=0.744), power (*p*=0.279) and muscle fibre CSA (*p*=0.534) (Table 2). These results were replicated with testosterone precursor and metabolite DHEA and DHT, respectively (all *p*-values >0.05) (Supplementary Table 3).

**Table 2.**
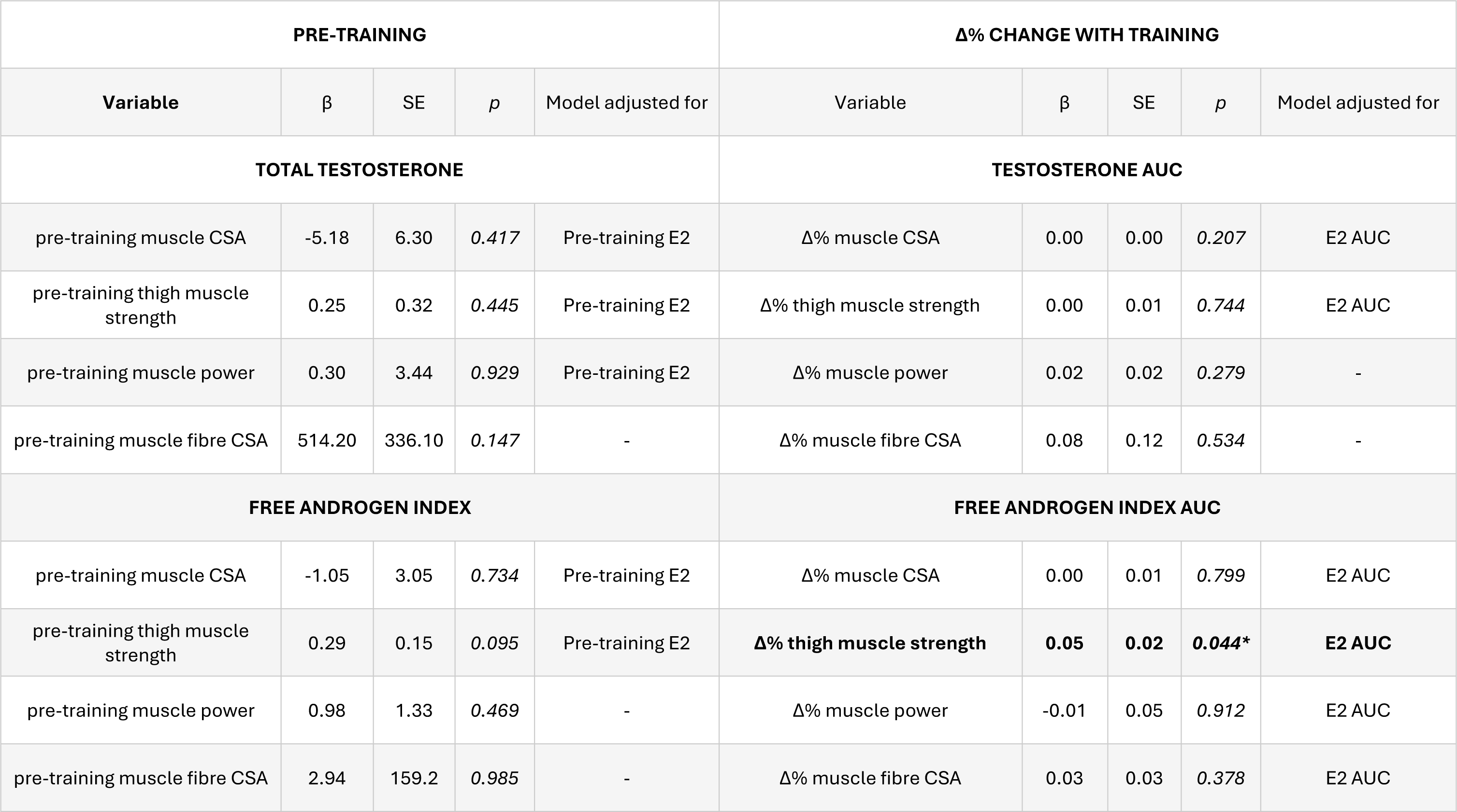
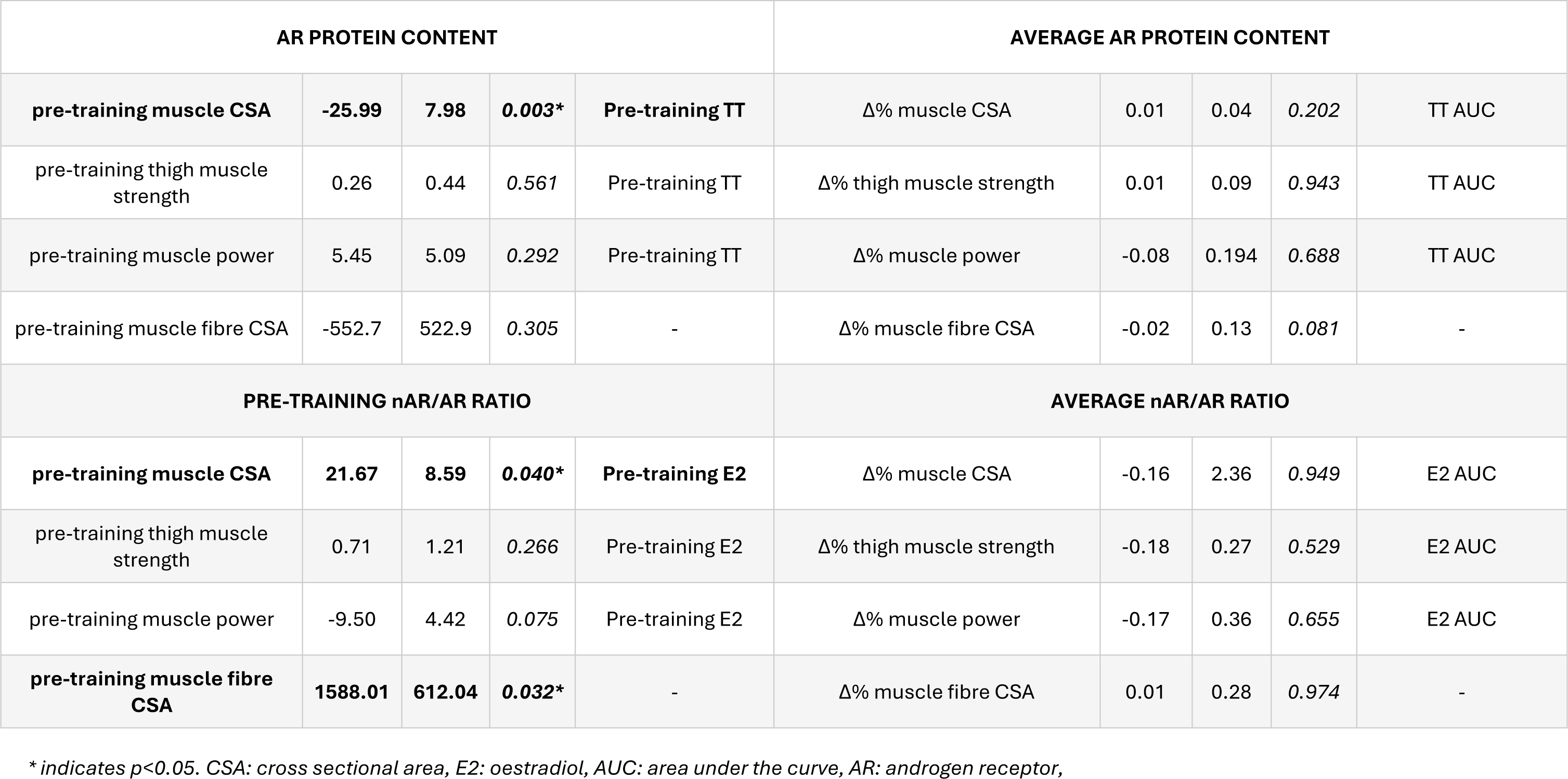
Linear mixed models of the association between total testosterone, the free androgen index, and markers of AR signalling and muscle size, strength and power before and after 12 weeks of resistance training in pre-menopausal females. n=35 pre-training, 27 change with training.

| PRE-TRAINING |  |  |  |  | Δ% CHANGE WITH TRAINING |  |  |  |  |
| --- | --- | --- | --- | --- | --- | --- | --- | --- | --- |
| Variable | β | SE | *p* | Model adjusted for | Variable | β | SE | *p* | Model adjusted for |
| TOTAL TESTOSTERONE |  |  |  |  | TESTOSTERONE AUC |  |  |  |  |
| pre-training muscle CSA | -5.18 | 6.30 | 0.417 | Pre-training E2 | Δ% muscle CSA | 0.00 | 0.00 | 0.207 | E2 AUC |
| pre-training thigh muscle strength | 0.25 | 0.32 | 0.445 | Pre-training E2 | Δ% thigh muscle strength | 0.00 | 0.01 | 0.744 | E2 AUC |
| pre-training muscle power | 0.30 | 3.44 | 0.929 | Pre-training E2 | Δ% muscle power | 0.02 | 0.02 | 0.279 | - |
| pre-training muscle fibre CSA | 514.20 | 336.10 | 0.147 | - | Δ% muscle fibre CSA | 0.08 | 0.12 | 0.534 | - |
| FREE ANDROGEN INDEX |  |  |  |  | FREE ANDROGEN INDEX AUC |  |  |  |  |
| pre-training muscle CSA | -1.05 | 3.05 | 0.734 | Pre-training E2 | Δ% muscle CSA | 0.00 | 0.01 | 0.799 | E2 AUC |
| pre-training thigh muscle strength | 0.29 | 0.15 | 0.095 | Pre-training E2 | **Δ% thigh muscle strength** | **0.05** | **0.02** | **0.044\*** | **E2 AUC** |
| pre-training muscle power | 0.98 | 1.33 | 0.469 | - | Δ% muscle power | -0.01 | 0.05 | 0.912 | E2 AUC |
| pre-training muscle fibre CSA | 2.94 | 159.2 | 0.985 | - | Δ% muscle fibre CSA | 0.03 | 0.03 | 0.378 | E2 AUC |

| AR PROTEIN CONTENT |  |  |  |  | AVERAGE AR PROTEIN CONTENT |  |  |  |  |
| --- | --- | --- | --- | --- | --- | --- | --- | --- | --- |
| **pre-training muscle CSA** | **-25.99** | **7.98** | **0.003\*** | **Pre-training TT** | $\Delta\%$ muscle CSA | 0.01 | 0.04 | 0.202 | TT AUC |
| pre-training thigh muscle strength | 0.26 | 0.44 | 0.561 | Pre-training TT | $\Delta\%$ thigh muscle strength | 0.01 | 0.09 | 0.943 | TT AUC |
| pre-training muscle power | 5.45 | 5.09 | 0.292 | Pre-training TT | $\Delta\%$ muscle power | -0.08 | 0.194 | 0.688 | TT AUC |
| pre-training muscle fibre CSA | -552.7 | 522.9 | 0.305 | - | $\Delta\%$ muscle fibre CSA | -0.02 | 0.13 | 0.081 | - |
| PRE-TRAINING nAR/AR RATIO |  |  |  |  | AVERAGE nAR/AR RATIO |  |  |  |  |
| **pre-training muscle CSA** | **21.67** | **8.59** | **0.040\*** | **Pre-training E2** | $\Delta\%$ muscle CSA | -0.16 | 2.36 | 0.949 | E2 AUC |
| pre-training thigh muscle strength | 0.71 | 1.21 | 0.266 | Pre-training E2 | $\Delta\%$ thigh muscle strength | -0.18 | 0.27 | 0.529 | E2 AUC |
| pre-training muscle power | -9.50 | 4.42 | 0.075 | Pre-training E2 | $\Delta\%$ muscle power | -0.17 | 0.36 | 0.655 | E2 AUC |
| **pre-training muscle fibre CSA** | **1588.01** | **612.04** | **0.032\*** | - | $\Delta\%$ muscle fibre CSA | 0.01 | 0.28 | 0.974 | - |

\* indicates $p < 0.05$ . CSA: cross sectional area, E2: oestradiol, AUC: area under the curve, AR: androgen receptor,

The same model was then used to investigate the association between the FAI, indicative of the amount of bioavailable testosterone, and whole thigh muscle CSA, strength, power and fibre CSA. There was no evidence of an association between pre-training FAI and muscle CSA, strength, fibre CSA and power at baseline (Table 2). The AUC of the FAI across 12-weeks of resistance training was positively associated to the changes in muscle strength (β=0.05, SE=0.02, *p*=0.044) that occurred with 12 weeks of resistance training (Figure 6A).

**Figure 6.**
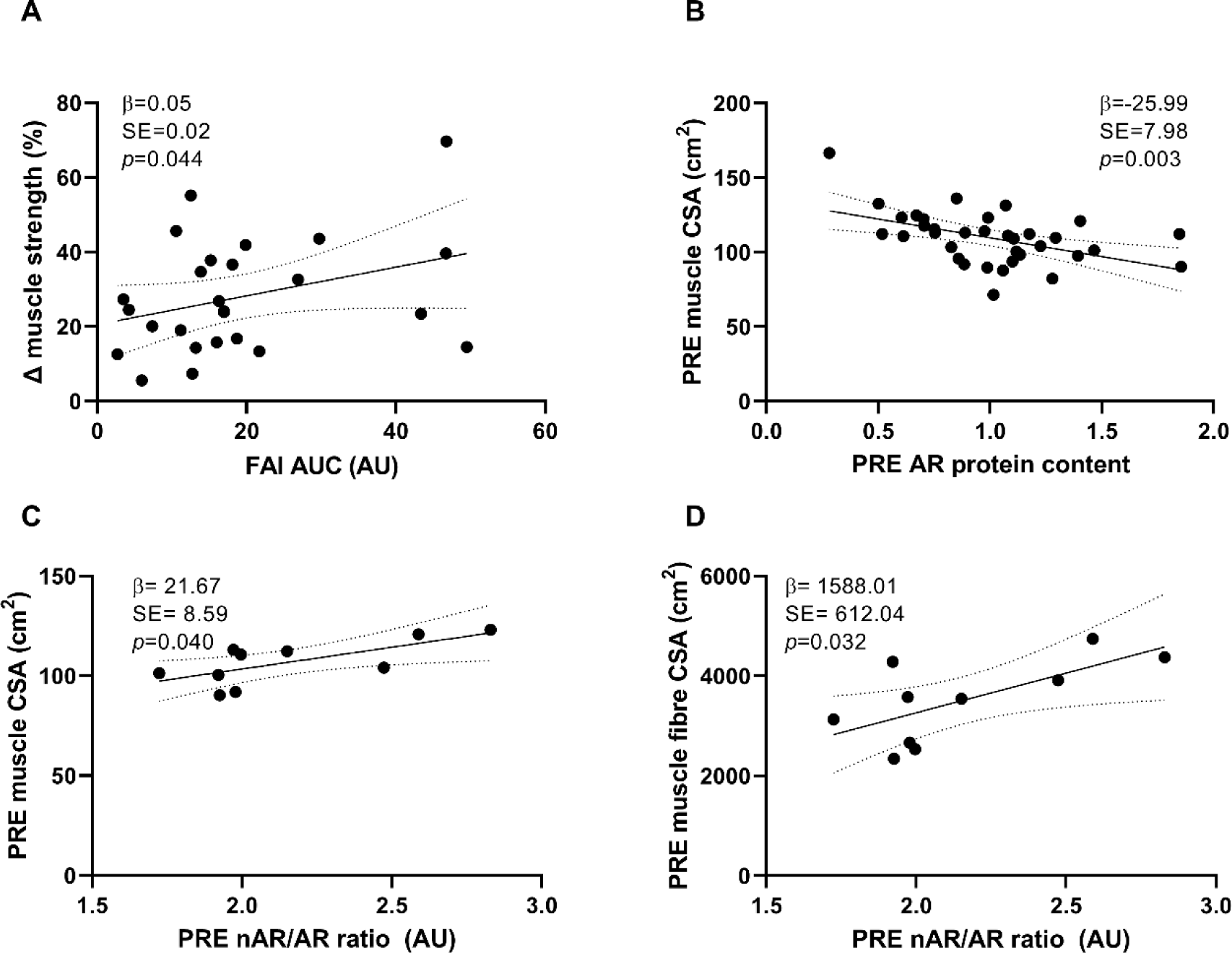
Linear associations between A) the free androgen index area under the curve (FAI AUC; AU) and the changes in muscle strength, (B) baseline AR protein content (AU) and baseline thigh muscle CSA (cm^2^) (C) pre-training nAR/AR ratio (AU) and pre-training muscle CSA (cm^2^) and (D) pre-training nAR/AR ratio (AU) and pre-training mixed muscle fibre CSA (cm^2^) in pre-menopausal females. Dashed lines represent 95% confidence intervals.

There were no correlations between total testosterone or the FAI and total or phospho-protein content of markers of protein synthesis Akt, mTOR, MAPK, rpS6 or 4E-BP1 before or after 12 weeks of resistance training (data not shown).

### 3.8 Associations between androgen receptor content and phosphorylation and muscle strength, size and power

In young males, the protein content of the androgen receptor (AR) is positively associated with resistance training-induced hypertrophy (Morton *et al*., 2018). We therefore tested the association between the total or phospho-protein content of the AR and muscle CSA, strength, power and fibre CSA prior to, or in response to a 12-week resistance training program. Total androgen receptor content was negatively associated with thigh muscle CSA pre-training (β=−25.99, SE=7.98, *p*=0.003; Table 2, Figure 6B), but this association was not maintained after resistance training. There was no significant association between the phospho-AR content and any outcomes as any time point (Supplementary Table 3).

### 3.9 Associations between androgen receptor nuclear localisation and muscle strength, size and power

Linear models assessed the association between AR cellular localisation and muscle mass, strength and power prior to, or in response to a 12-week resistance training program (Table 2). The ratio of nAR to total AR intensity (nAR/AR ratio) was positively associated with whole muscle CSA (β=21.67, SE=8.59, *p=*0.04; Figure 6C) and muscle fibre CSA (β=1588.01, SE=612.04, p=0.032; Figure 6D), but not strength or power pre-training. There were no significant associations between the nAR/AR ratio and resistance-training induced changes in muscle mass, strength or power (Table 2). There were trends (*p=0.05-0.08*) for the percentage of nuclei that were AR+ to be positively associated with pre-training muscle CSA, strength and myofibre CSA, but these did not reach statistical significance (Supplementary Table 3).

### 3.10 Testosterone treatment increases female human primary myotube diameter through the nuclear translocation of the AR but does not activate the Akt/mTOR pathway

To further examine whether testosterone plays a direct, regulatory role in female skeletal muscle, we cultured human primary myocytes from 6 donor participants from the human study. Seven days of testosterone treatment increased myotube diameter by 37% compared to a vehicle control (*p<*0.01) and increased total AR protein content by over 4-fold, 3-fold and 2-fold after 1, 4 and 7 days of testosterone treatment, respectively (*p<*0.001; Figure 7A-B). This result was replicated using immunohistochemical staining, where the intensity of the AR (indicative of AR content) was significantly greater after 1 and 7 days of testosterone treatment, compared to a vehicle control (*p*<*0.*01; Figure 7C.) However, the protein content of Akt, p-Akt, mTOR, p-mTOR and p-MAPK did not change with testosterone treatment at any time point, despite their expression levels fluctuating across the differentiation time course (*p*<0.05; Supplementary Figure 7). Instead, immunohistochemical staining showed that, in myocytes treated with a vehicle control, the AR is distributed throughout the cytosol of the cell with no or little nuclear localisation (Figure 7D, upper panel). After 24 hours of 100 nM testosterone treatment, there was a striking translocation of the AR to the nucleus of myoblasts (Figure 7D, lower panel) paralleled by an increase in AR protein content in the cell. The AR protein then remained in the nucleus across the 7 days of differentiation of the testosterone treated myotubes (Figure 7E).

**Figure 7.**
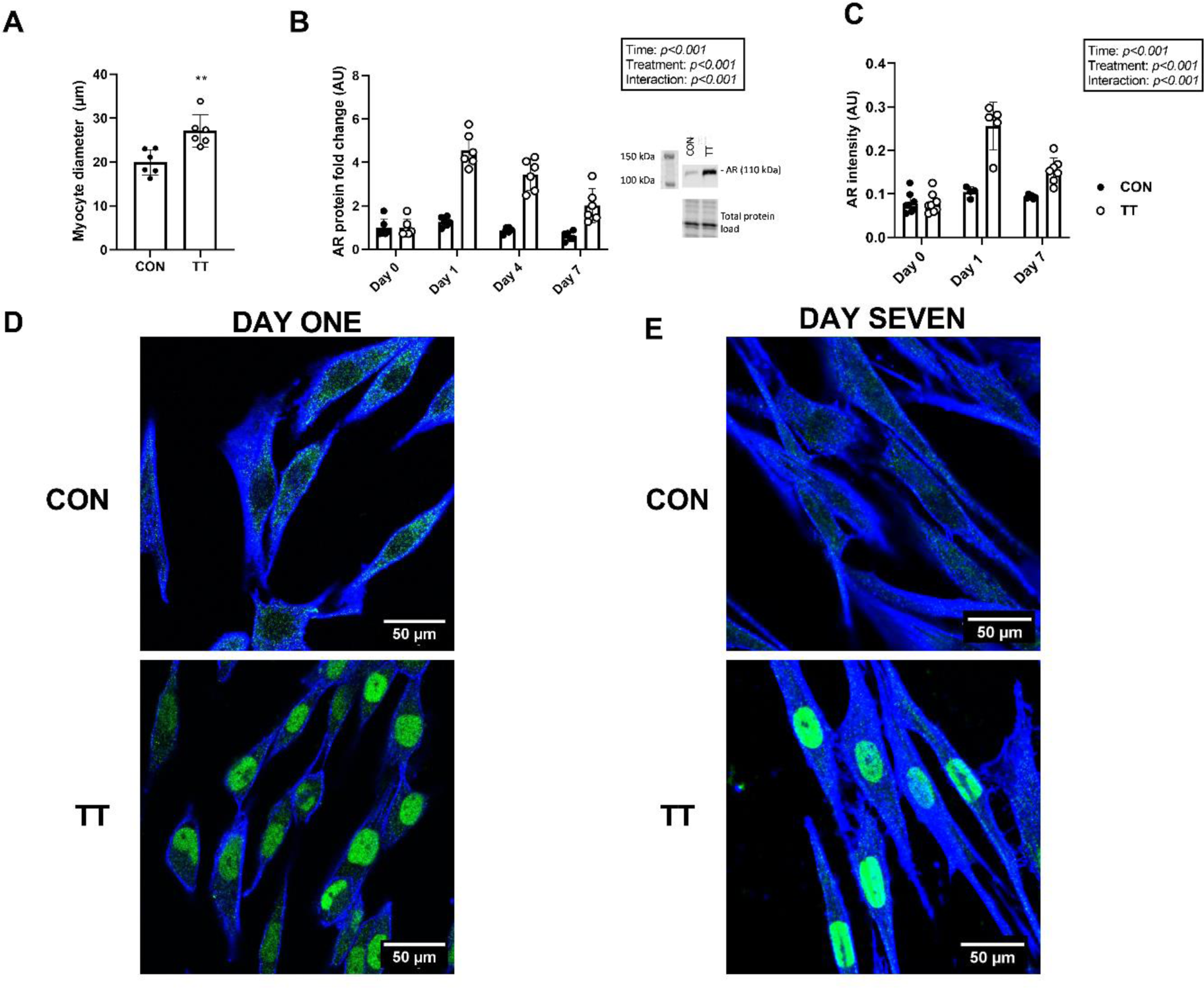
Testosterone treatment over 7 days of differentiation increased (A) myotube diameter, (B) AR protein content and (C) AR intensity relative to the proportion of the visual field occupied by myocytes. AR cellular localisation in primary muscle cell lines treated with vehicle (CON) or 100 nM testosterone (TT) after D) 1 day or E) 7 days of treatment (n=6 female donors). Phalloidin (stains actin) appears in blue. AR appears in green. Scale is 0.25 µm·pixel^-1^ for all images. White scale bar represents 50 µm. Data were analysed using two-tailed, paired t-tests and two-way ANOVA. Values are represented mean ± SD. **indicates p<0.01.

## 4. Discussion

We showed that total testosterone was not associated with muscle CSA, strength, power or the muscle anabolic response to 12 weeks of resistance training in pre-menopausal females. Transcriptomic data support the hypothesis that androgen genomic signalling, via the androgen response element, does not play a significant role in determining the muscle transcriptomic profile pre- or post-resistance training. In contrast, the bioavailable fraction of testosterone and the nAR/AR ratio were positively associated with muscle mass and strength at baseline, but not with resistance-training induced changes in muscle mass or strength. These findings shed light on the limited body of knowledge regarding the role of testosterone in the regulation of female skeletal muscle.

Our results demonstrate that total testosterone is not a direct determinant of muscle mass, strength, or the muscle response to anabolic stimulation in pre-menopausal females. These results are in accordance with previous cross-sectional data showing that total testosterone is not associated with muscle mass or strength in pre- and post-menopausal females (Gower & Nyman, 2000; Carmina *et al*., 2009; van Geel *et al*., 2009; Pöllänen *et al*., 2011; Rariy *et al*., 2011; Kogure *et al*., 2015). While there is some cross-sectional evidence of a positive relationship between testosterone and muscle mass in males (Mouser et al., 2016), there is also no relationship between testosterone and resistance-training induced adaptations in training in males (Morton et al., 2016; Morton et al., 2018). These results suggest that the association between testosterone and skeletal muscle differs between endogenous and exogenous testosterone. We posit that there are only positive associations between total testosterone and muscle mass and function when homeostasis is perturbed and concentrations are manipulated pharmacologically to above (Ferrando *et al*., 1998; Sheffield-Moore *et al*., 1999; Bhasin *et al*., 2001; Sinha-Hikim *et al*., 2004; Hirschberg *et al*., 2020) or below (Mauras *et al*., 1998; Overkamp *et al*., 2023) physiological concentrations in both males and females.

The bioavailable fraction of testosterone measured via the FAI was positively associated with resistance training-induced muscle strength, suggesting that the bioavailable rather than the total fraction of testosterone may play a regulatory role in the anabolic response of female skeletal muscle. In line with these findings, the FAI was positively associated with resistance training-induced thigh muscle hypertrophy in a small study of pre-menopausal females (*n*=5) (Häkkinen *et al*., 1992), and to muscle mass in a large cross-sectional cohort of pre-menopausal females conducted by our group (*n*=706) (Alexander *et al*., 2021). However, it is important to bear in mind that the effect sizes in this study and our previous work were small (β=0.05 and 0.01, respectively). While the association was significant, the bioavailable fraction of testosterone only explains a small proportion of the variance in muscle mass in pre-menopausal females.

We show, for the first time, a negative association between total AR protein content and whole muscle CSA in pre-menopausal females. This is in contrast to males, where the AR protein content was positively associated with resistance training-induced hypertrophy and strength in healthy young (Ahtiainen *et al*., 2011; Mitchell *et al*., 2013; Morton *et al*., 2018) and older (Ahtiainen *et al*., 2011) males, further suggesting sex-specific differences in the role of the AR in skeletal muscle regulation. In support of this finding, male AR knockout (ARKO) mice displayed significant reductions in muscle mass and strength compared to their wildtype littermates (MacLean *et al*., 2008). Conversely, female ARKO mice did not display any differences in muscle mass or strength compared to their wildtype controls (MacLean *et al*., 2008), further demonstrating sex-specific differences in the role of the AR in the maintenance of muscle mass and function. We also confirm that resistance training does not affect AR protein content, phosphorylation status or nuclear localisation in female skeletal muscle, in agreement with previous work showing no change in AR protein content or nuclear localisation following 10 weeks of resistance training (n=13 females) (Hatt *et al*., 2024). Males, in contrast, display significant increases in both AR protein content and nuclear localisation after the same resistance training program (Hatt *et al*., 2024), suggesting further sex-specific AR regulation with chronic resistance exercise.

The proportion of AR localised to the nucleus (nAR/AR ratio) was positively associated with muscle size at both the fibre and whole-muscle level. Therefore, the ability to recruit the AR and translocate it to the nucleus rather than total AR content or testosterone concentrations may be more physiologically relevant to the maintenance of muscle mass in females. Taken together, these results point towards a negative feedback loop between total AR and muscle mass regulation. The positive association between nAR/AR and muscle size suggests that individuals with increased AR sensitivity, and therefore a greater ability to recruit and translocate AR to the nucleus, may require less total AR protein content to maintain their muscle mass. This is supported by our *in vitro* findings showing that, after an initial increase in total AR protein content with testosterone treatment, AR protein content begins to return to baseline levels after 4 and 7 days of treatment. Despite this decrease in total AR protein content, the amount of AR in the nucleus was sustained across 7 days of treatment, suggesting a negative regulation of the AR by testosterone or with increased AR sensitivity.

Our data showing a rapid increase in AR protein content and nAR within 24 hours of testosterone treatment *in vitro* are in line with previous findings showing that 6 days of testosterone treatment increased AR protein content in primary muscle cell cultures from male donors *in vitro* (Sinha-Hikim *et al*., 2004) and in muscle biopsies obtained from healthy, young males *in vivo* (*n=*6) after 20 weeks of treatment with 600 mg·week^-1^ testosterone (Sinha-Hikim *et al*., 2004). Taken together, our data and others indicate that testosterone treatment primarily increases myotube diameter through genomic AR signalling and induces a marked and sustained translocation of the AR into myonuclei *in vitro* and *in vivo* (Bhasin *et al*., 2001), rather than non-genomic signalling pathways such as the Akt/mTOR or MAPK as previously suggested (Wu *et al*., 2010; Basualto-Alarcón *et al*., 2013; White *et al*., 2013).

No markers of protein synthesis or degradation were associated with total or bioavailable testosterone in our human cohort. This is mirrored by our findings that testosterone treatment did not promote Akt/mTOR or MAPK signalling in myocytes taken from female donors, despite significant increases in myotube diameter with testosterone treatment. In support of this, exogenous testosterone administration did not change molecular regulators of muscle mass and mitochondrial biosynthesis, including markers of mTOR signalling from resting biopsies in males (*n=*50) (Howard *et al*., 2020) and females (*n*=48) (Horwath *et al*., 2022) *in vivo*, or female primary myocytes *in vitro* (Pataky *et al*., 2023). Instead, testosterone administration to primary myotubes from female donors induced changes of proteins within the sarcoplasmic compartment, including myosin and titin, both of which play important roles in the contractile apparatus and muscle hypertrophy (Pataky *et al*., 2023). This suggests that, in contrast to *in vitro* data from rat (Wu *et al*., 2010; White *et al*., 2013) and murine (Basualto-Alarcón *et al*., 2013) myocytes, the primary mechanism of action of testosterone in humans may not be through the upregulation of the mTOR pathway. Instead, the changes in lean mass seen with exogenous testosterone administration in males (Bhasin *et al*., 2001; Howard *et al*., 2020) and females (Horwath *et al*., 2022) may stem from a net positive protein turnover in favour of protein accretion driven by an increase in the genomic signalling of the AR rather than through activation of the Akt/mTOR non-genomic signalling pathways. This genomic signalling may lead to an increase in transcription of target genes and eventually in translational capacity, as evidenced by increases in total ribosome number (Mobley *et al*., 2018) and muscle RNA content (Howard *et al*., 2020).

### 4.1 Limitations

While this study included 4 participants (∼15%) who had been diagnosed with polycystic ovary syndrome (PCOS), there was only 1 participant who consistently had total testosterone concentrations above the typical female reference range (testosterone levels >2.5 nmol·L^-1^ (Burger, 2002)). This limited range of testosterone concentrations restrict the generalisability of these results. Our previous research (Alexander *et al*., 2021) suggests that the association between the FAI and lean mass is not linear but quadratic in nature, where the association plateaus and eventually becomes negative with increasing testosterone concentrations. Including females with a wider range of testosterone levels, including hyperandrogenic females and individuals with DSD, would increase the generalisability of our findings and allow to validate the association between testosterone and muscle across a larger spectrum of androgen concentrations.

### 4.2 Conclusions

Our results suggest that, rather than total circulating testosterone concentrations, an individual’s sensitivity to bioavailable androgens and their ability to recruit the AR to the nucleus plays a significant role in the maintenance of muscle mass and strength in pre-menopausal females. This suggests that findings from studies demonstrating a large anabolic effect of testosterone administration on muscle are not generalisable to the association between endogenous testosterone concentrations and muscle mass and function and should therefore be interpreted with caution.

## Availability of data and materials

All RNA sequencing data generated or analysed during this study are included in this published article, its supplementary information files and publicly available repositories (GEO: link pending). The R code used for the analysis is available at https://github.com/DaniHiam/TESTO_RNAseq

## Acknowledgments

SeL received the protein supplement used in this study as in-kind payment for consulting work performed for Ascent Protein, Denver. The authors would like to acknowledge and thank Dr Raul Nicoli, Dr Carine Schweizer and Dr Tia Kuuranne of the Swiss Laboratory for Doping Analysis for their contribution to the hormone analysis and feedback on the document. The authors also wish to thank Dr Jessica Silver and Ms Ashwinder Kaur Goshel for their help delivering the training program. Finally, we would like to thank all our participants.

## Authors’ contributions

SEA was involved in the design and conception of the study, and performed data collection, laboratory and scientific analyses and preparation the manuscript. OEK was involved in the design of the training program and the delivery of the training program. RMW was involved with laboratory and statistical analyses. BG and KF were involved with data collection. PJ performed body composition scans. PDG was involved with laboratory analyses and the design of the *in vitro* experiments. AG performed all muscle biopsies. SeL was involved in the design and conception of the studies and preparation of the manuscript. DH performed statistical and transcriptomic analyses. GDW and BA were involved in the design and conception of the study. All authors contributed to the editing and reviewing of the manuscript. All authors approved the manuscript in its final form.

---

**Supplementary Figure 1.**
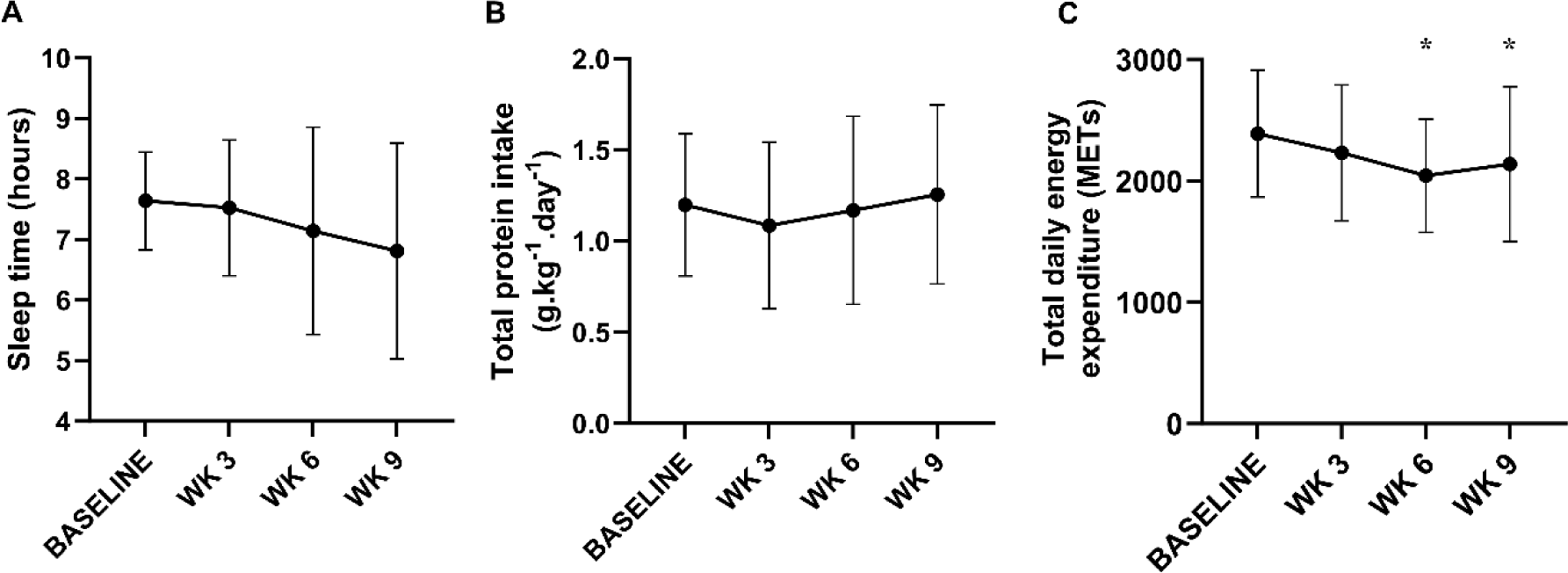
A) total sleep time did not differ across 12 weeks of resistance training in pre-menopausal females. B) Total protein intake (g·kg^-1^·day^-1^) did not fluctuate with 12 weeks of resistance training in pre-menopausal females. C) The total daily energy expenditure (METs) decreased at 6 weeks compared to baseline, but at no other timepoints across 12 weeks of resistance training in pre-menopausal females. Data were analysed via one-way ANOVA *denotes significant post-hoc test (*p*<0.05 compared to baseline).

**Supplementary Figure 2.**
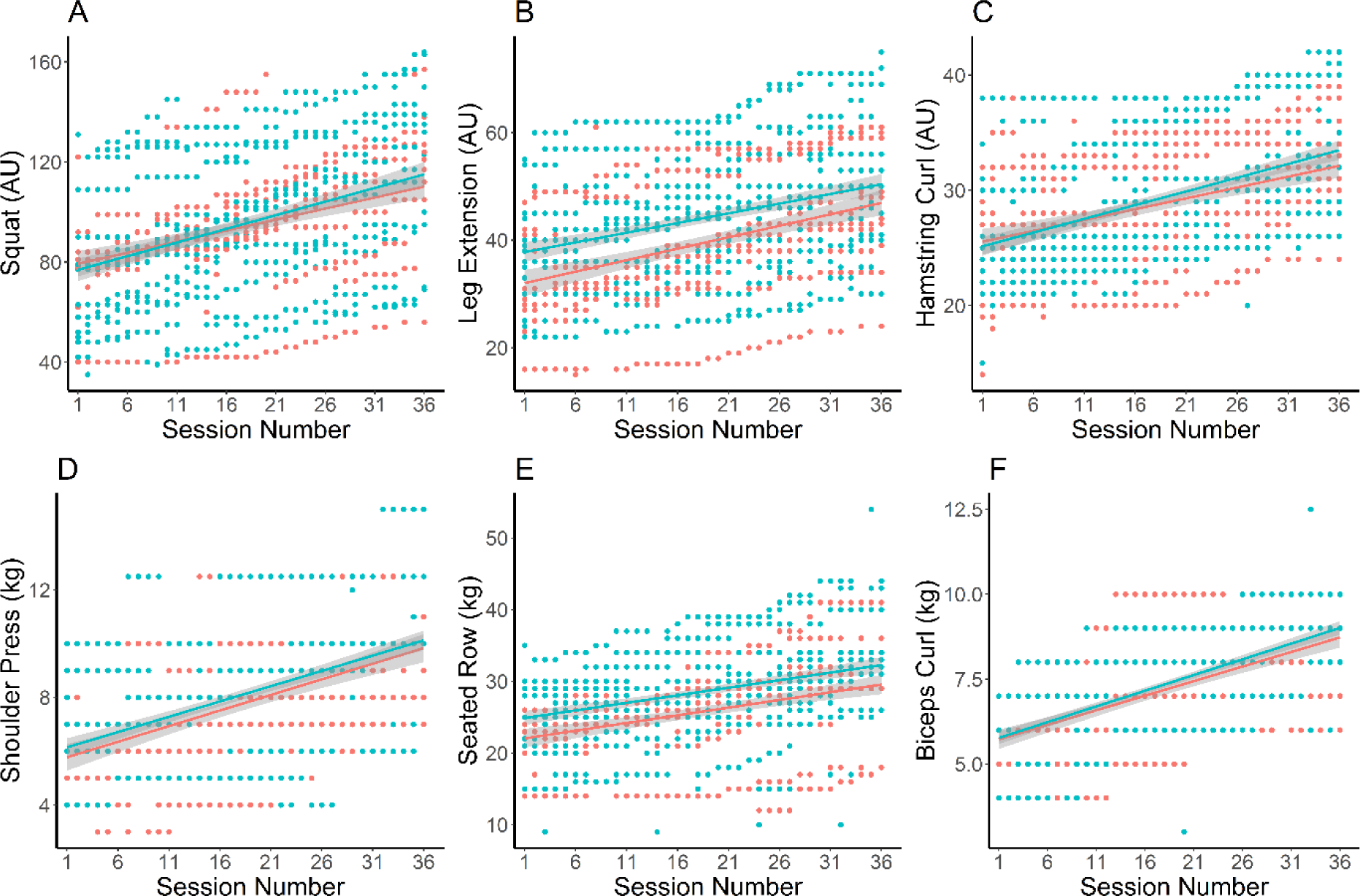
The trajectory of working weight progression for (A) squat, (B) leg extension, (C) hamstring curl, (D) shoulder press, (E) seated row and (F) biceps curl. Participants undertaking a 12-week gym-based training program are denoted in blue (n=16), and participants undertaking the blended resistance training program are denoted in red (n=11).

**Supplementary Figure 3.**
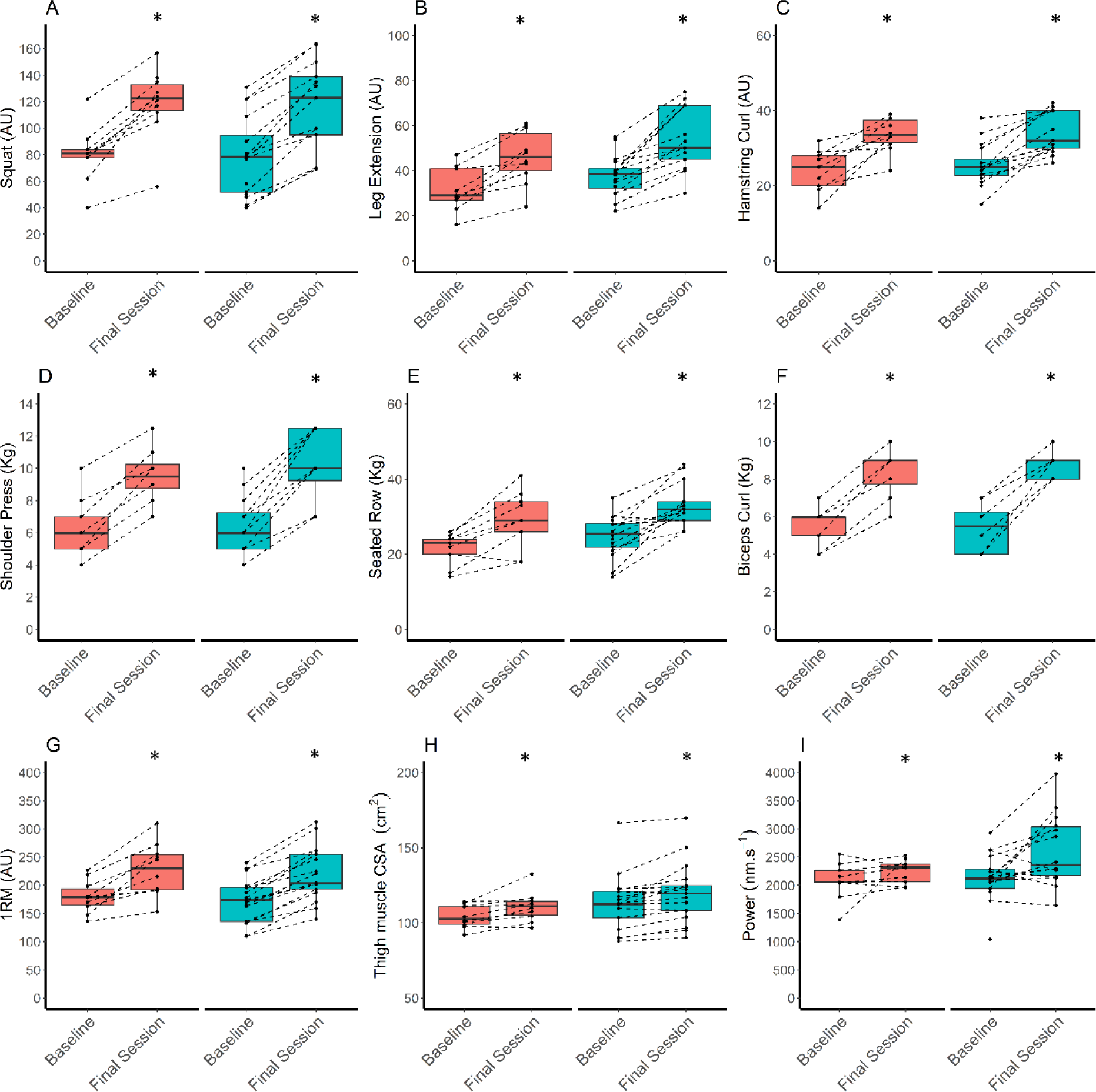
Working weight used in the first and final session for each participant. (A) squat, (B) leg extension, (C) hamstring curl, (D) shoulder press, (E) seated row and (F) biceps curl. (G) Lower limb strength (measured via leg press 1RM), (H) thigh muscle cross sectional area (CSA; cm^2^) and (I) muscle power (measured via counter movement jump) increased with 12 weeks of resistance training but this was not different between participants who undertook either a gym-based or blended training program. Participants undertaking a 12-week gym-based training program are denoted in blue (n=16), and participants undertaking the blended resistance training program are denoted in red (n=11). *indicates *p<0.05* compared to baseline.

**Supplementary Figure 4.**
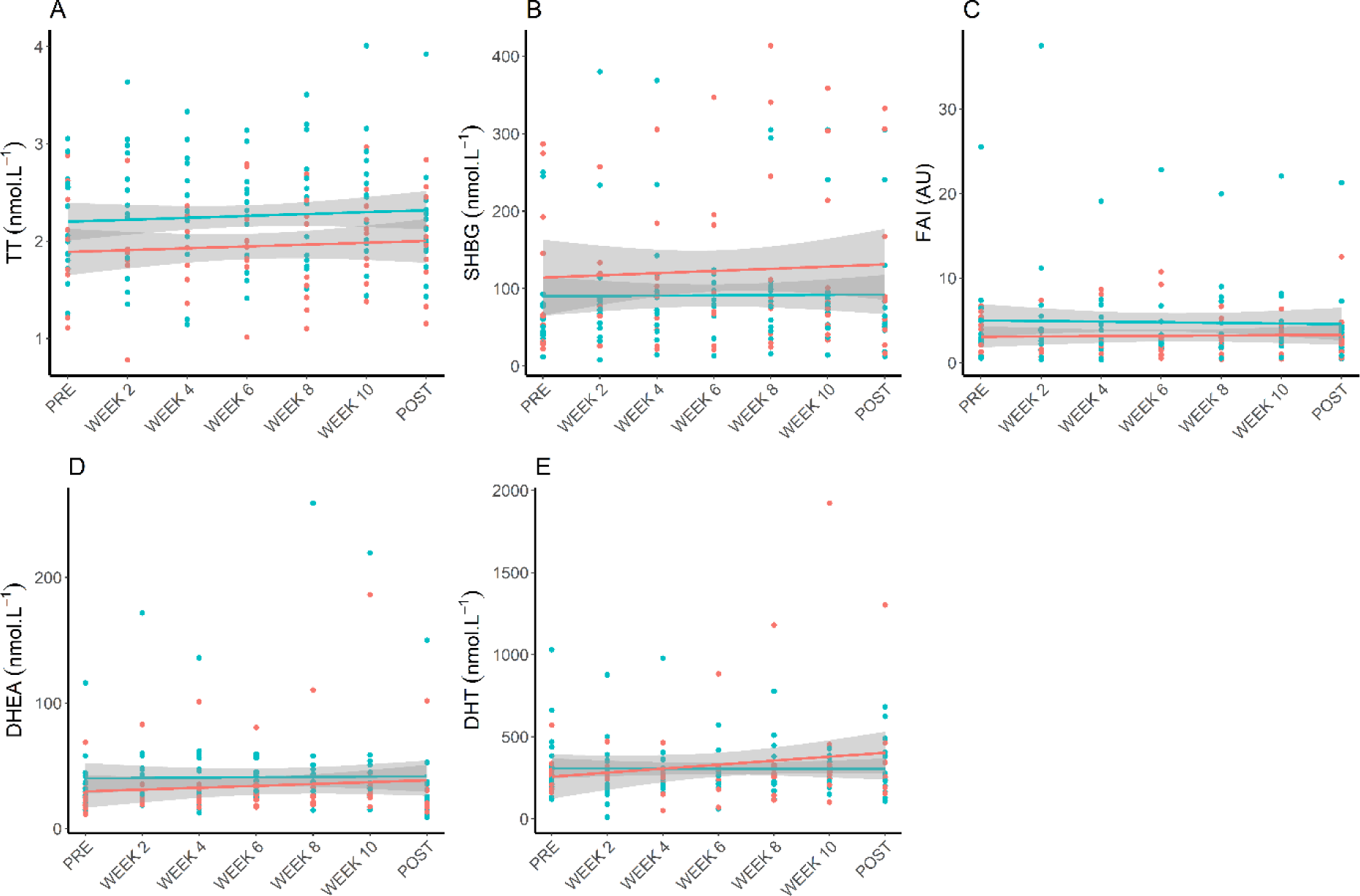
Hormone levels across 12 weeks of resistance training. (A) Total testosterone (TT), (B) sex hormone binding globulin (SHBG) and (C) free androgen index (FAI), (D) DHEA, (E) DHT concentrations. Participants undertaking a 12-week gym-based training program are denoted in blue (n=16), and participants undertaking the blended resistance training program are denoted in red (n=10).

**Supplementary Figure 5.**
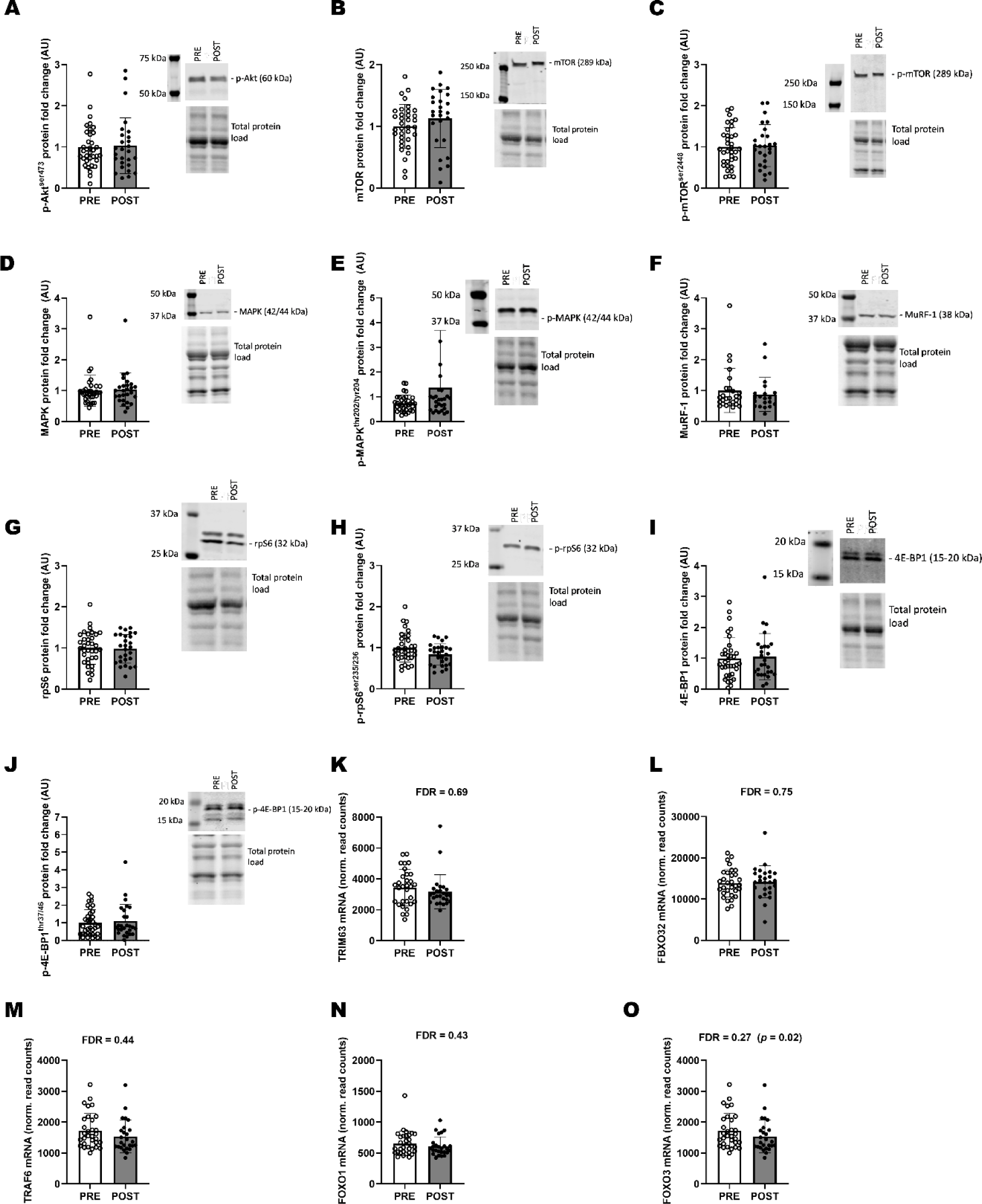
Twelve weeks of resistance training did not influence the protein content of markers of protein synthesis and degradation in (A) p-Akt^ser473^, (B) total mTOR, (C) p-mTOR^ser2448^, (D) total MAPK, (E) p-MAPK^thr202/tyr204^, (F) MuRF-1, (G) S6 ribosomal protein, (H) p-S6 ribosomal protein^ser235/236^, (I) total 4E-BP, (J) p-4E-BP1^thr37/46^ or the mRNA content of markers of protein degradation in (K) TRIM63, (L) FBXO32, (M) TRAF6, (N) FOXO1 and (O) FOXO3 in biopsies taken at rest from previously untrained, pre-menopausal females (n=35 baseline, n=27 post training). Data were analysed via two-tailed, paired t-tests. Representative western blots for each protein are presented next to the corresponding graph. All proteins were normalised against total protein content, mRNA was normalised to read counts per million. Pre-training (PRE) values are indicated by clear bars and post-training (POST) values are indicated by dark bars. Bars indicate mean ± SD.

**Supplementary Figure 6.**
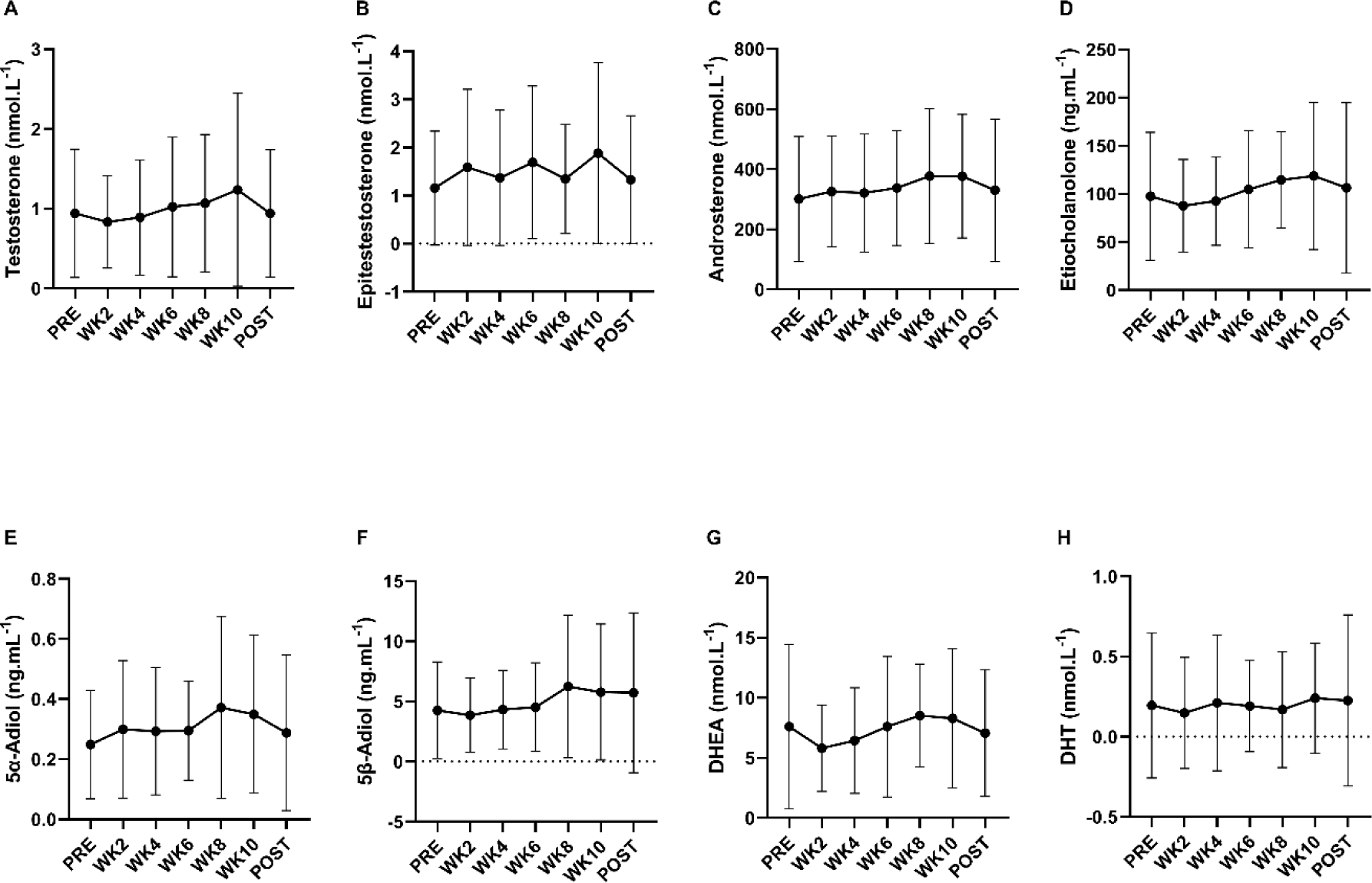
The androgen profile measured by LC/MS from urine in pre-menopausal females is not influenced by resistance training. The concentrations of a) testosterone (nmol·L^-1^), b) epitestosterone (nmol·L^-1^), c) androsterone (nmol·L^-1^), d) etiocholanolone (ng·mL^-1^), e) 5α-adiol (ng·mL^-1^), f) 5β-adiol (ng·mL^-1^), g) DHEA (nmol·L^-1^) and h) DHT (nmol·L^-1^) were stable across twelve weeks of resistance training in our cohort (n=27). Data were analysed with one-way ANOVA. Bars indicate mean ± SD.

**Supplementary Figure 7.**
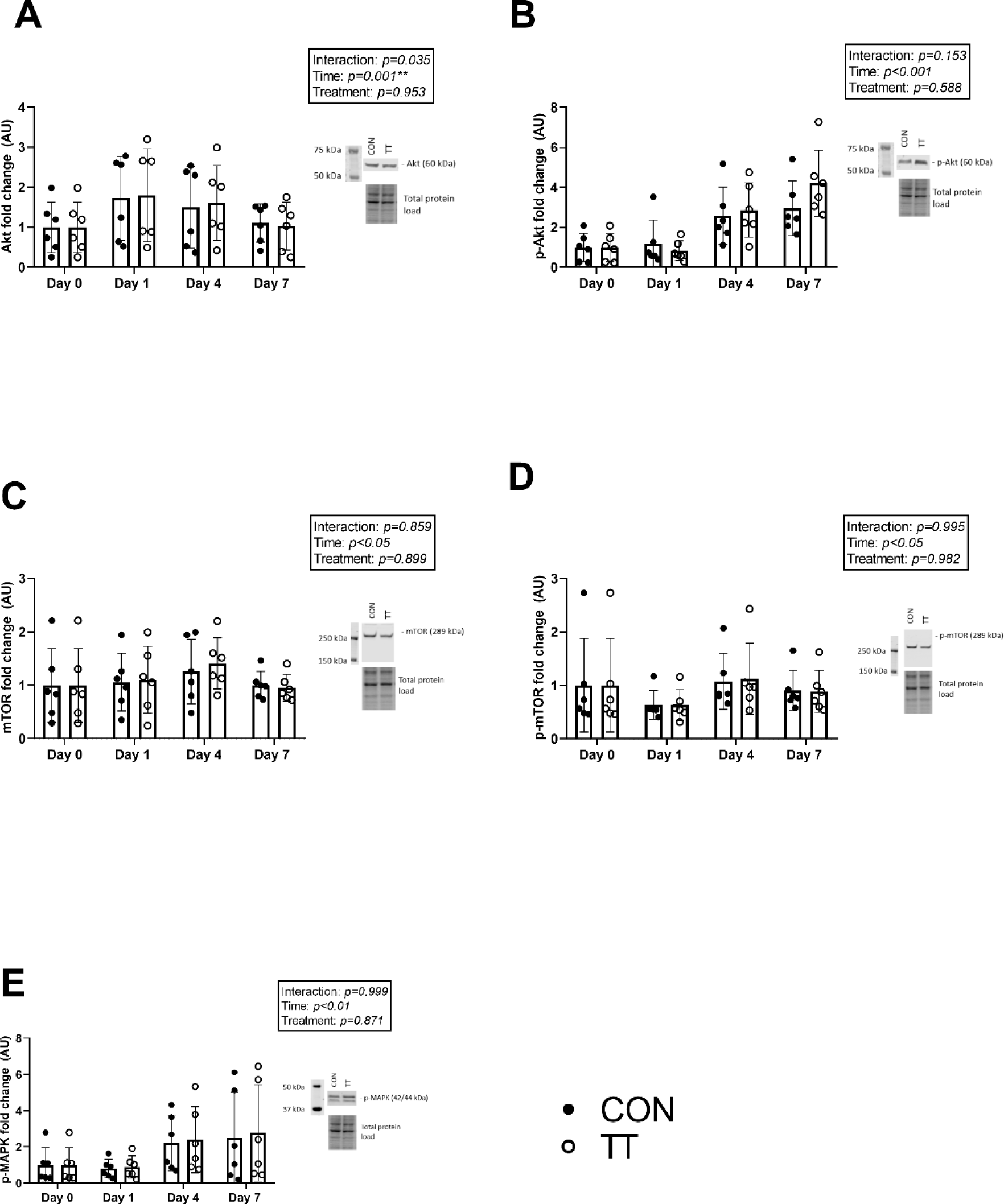
Treatment with 100 nM testosterone did not significantly change the expression of A) Akt, B) p-Akt, C) mTOR, D) p-mTOR or E) p-MAPK compared to a vehicle control across 1, 4 or 7 days of differentiation. Black circles represent vehicle control condition (CON), white circles represent testosterone treated (TT) condition. Data were analysed via two-way, repeated measures ANOVA. Bars represent mean ± SD.

**Supplementary Table 1.**
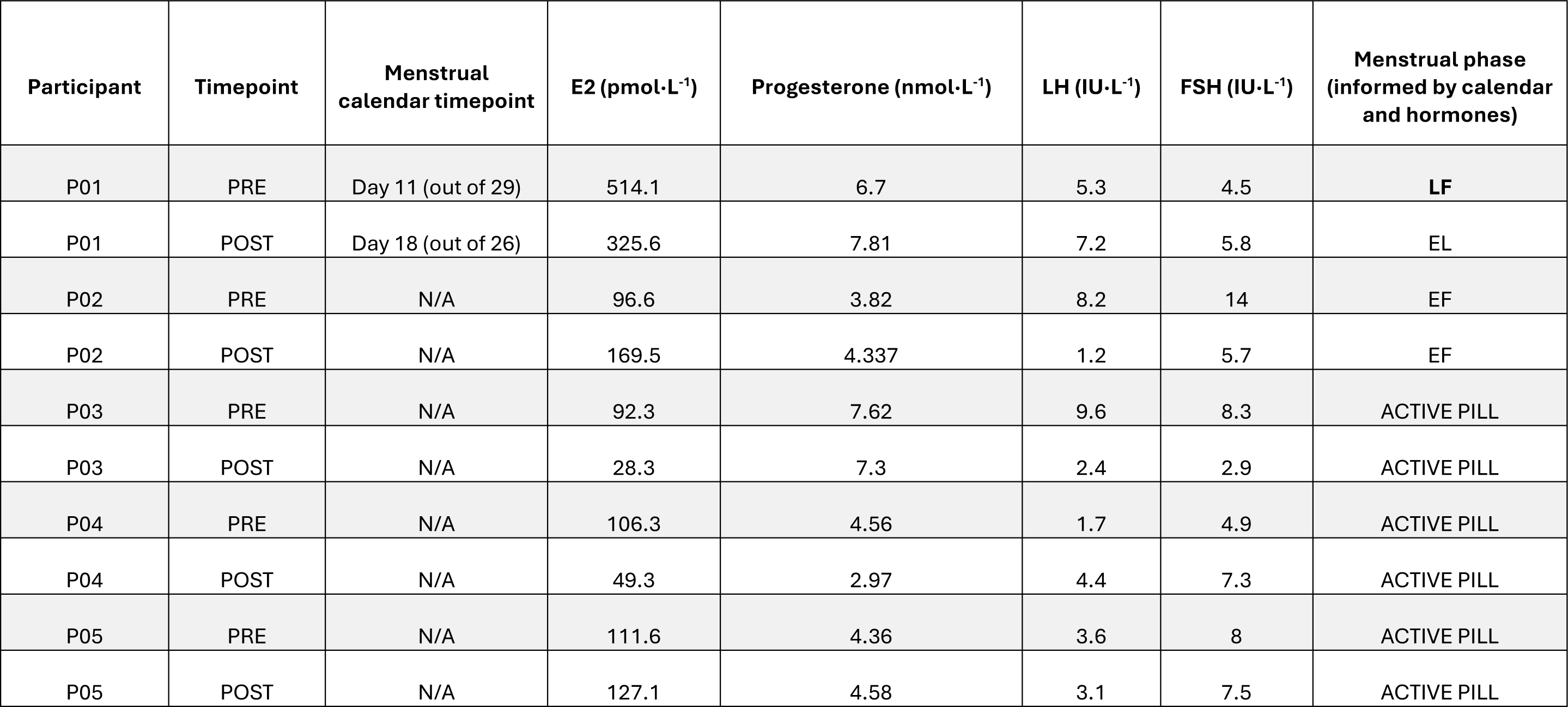
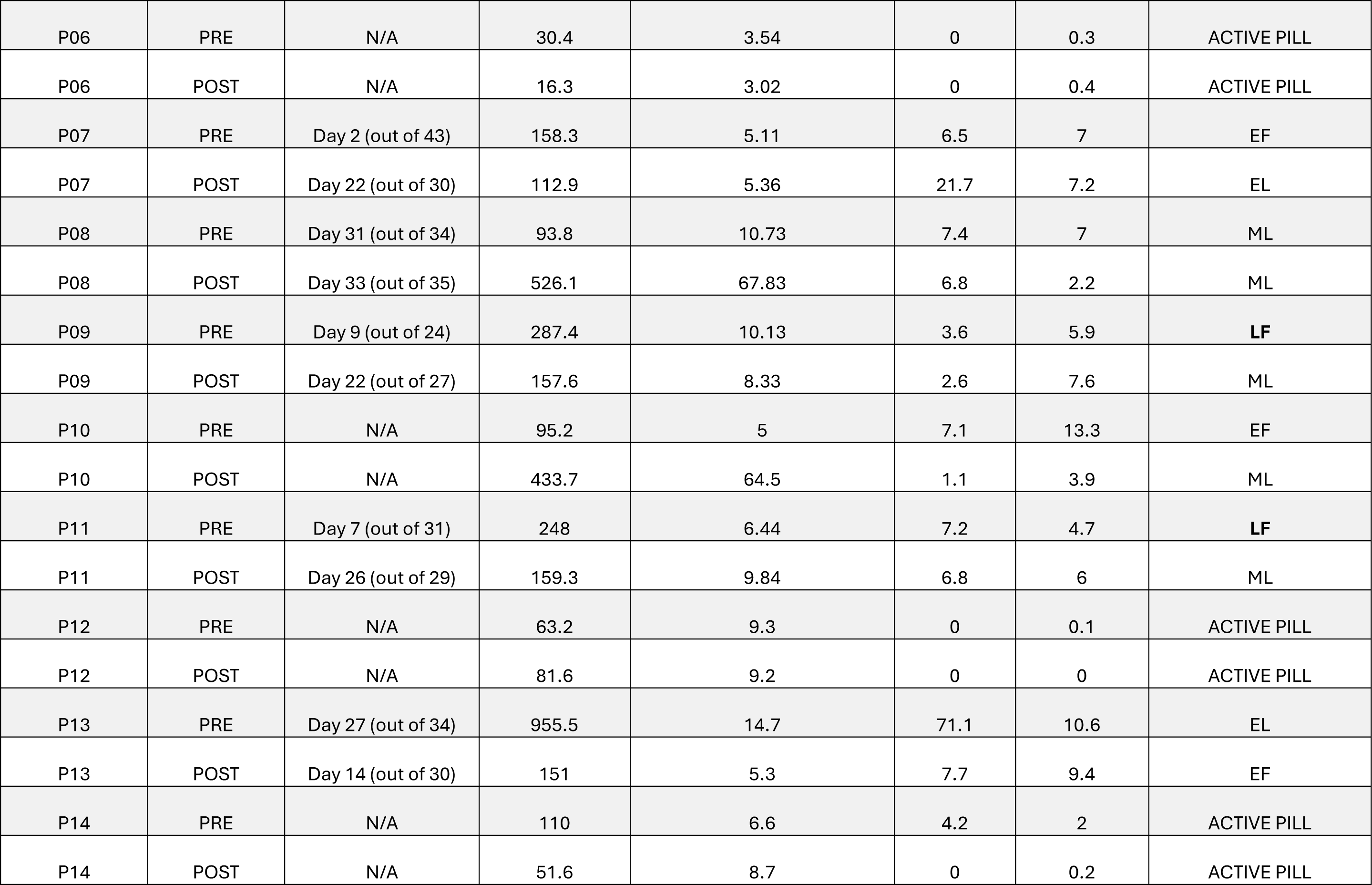
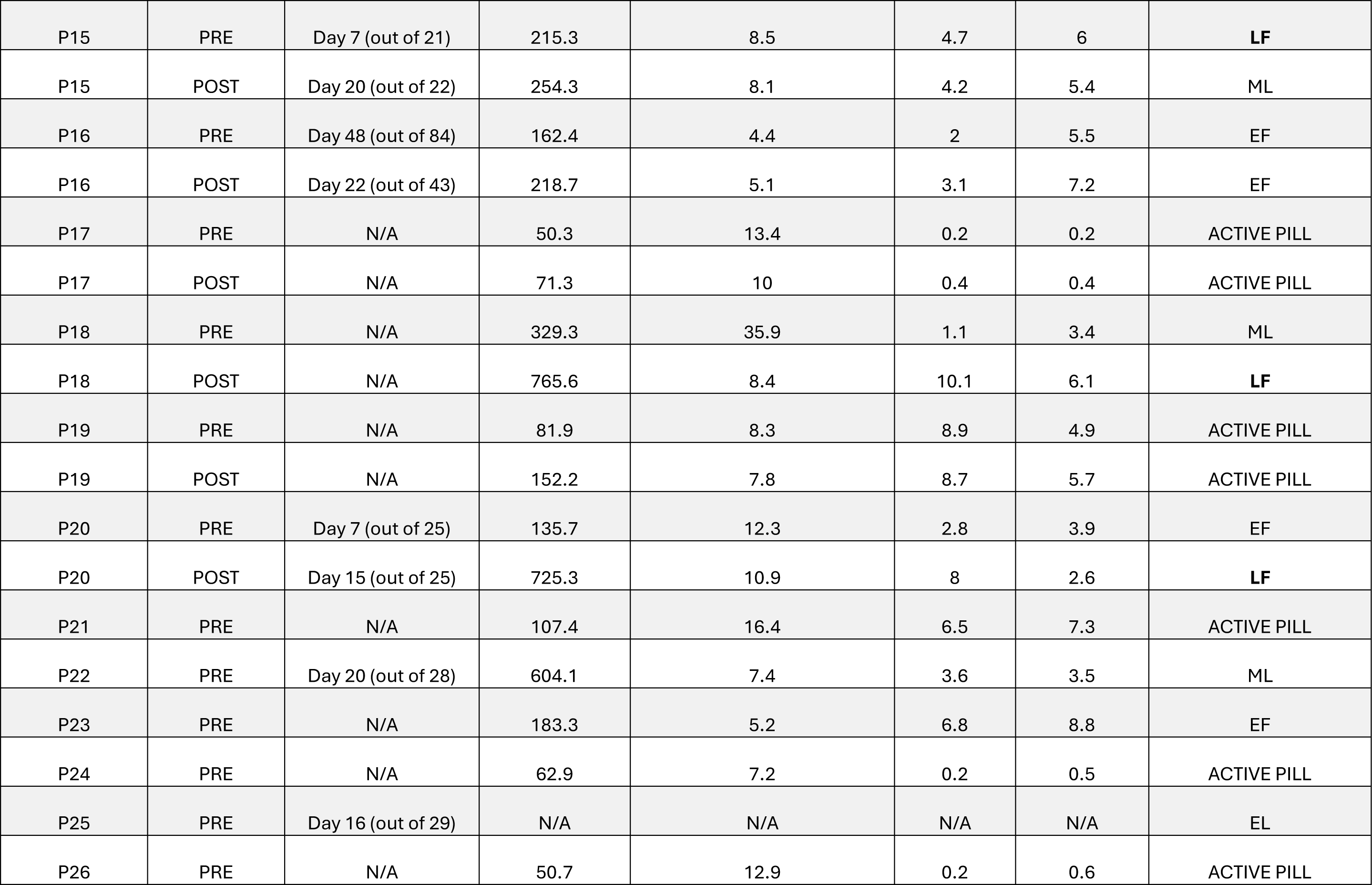
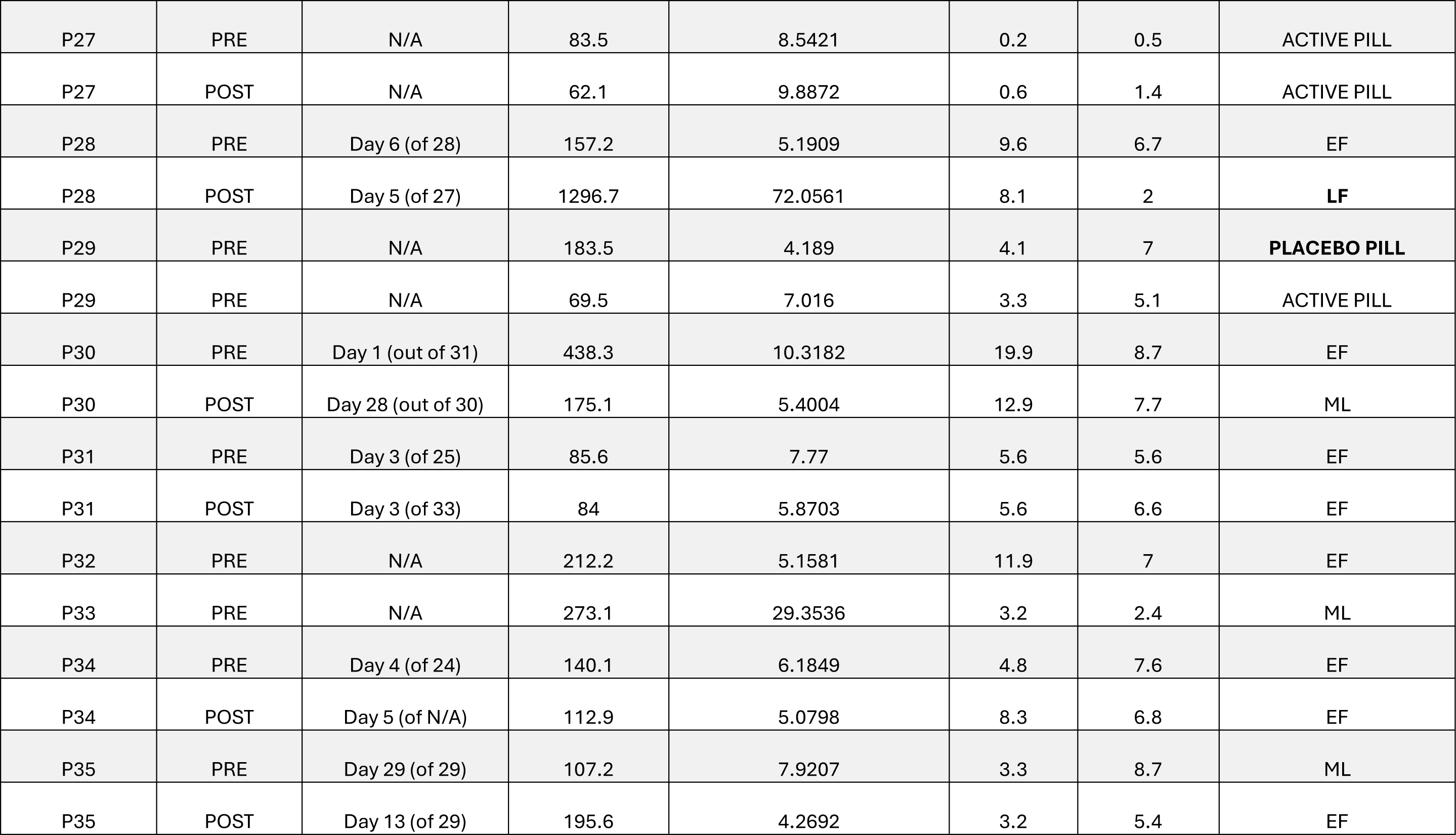
Summary of the menstrual phases of each participant at pre- and post-training testing timepoints. The menstrual phase was informed by a menstrual calendar collected by the participant and circulating concentrations of oestradiol (E2), progesterone, luteinising hormone (LH) and follicle stimulating hormone (FSH). Two independent researchers determined the menstrual phase for each entry and agreement was met at each case. Note: This study aimed to avoid testing and biopsy times during the late follicular (LF) phase of the menstrual cycle. Seven out of 62 (11%) testing times were taken during the late follicular phase. Statistical analyses will include the menstrual phase as a covariate to account for these times.

**Supplementary Table 2.**
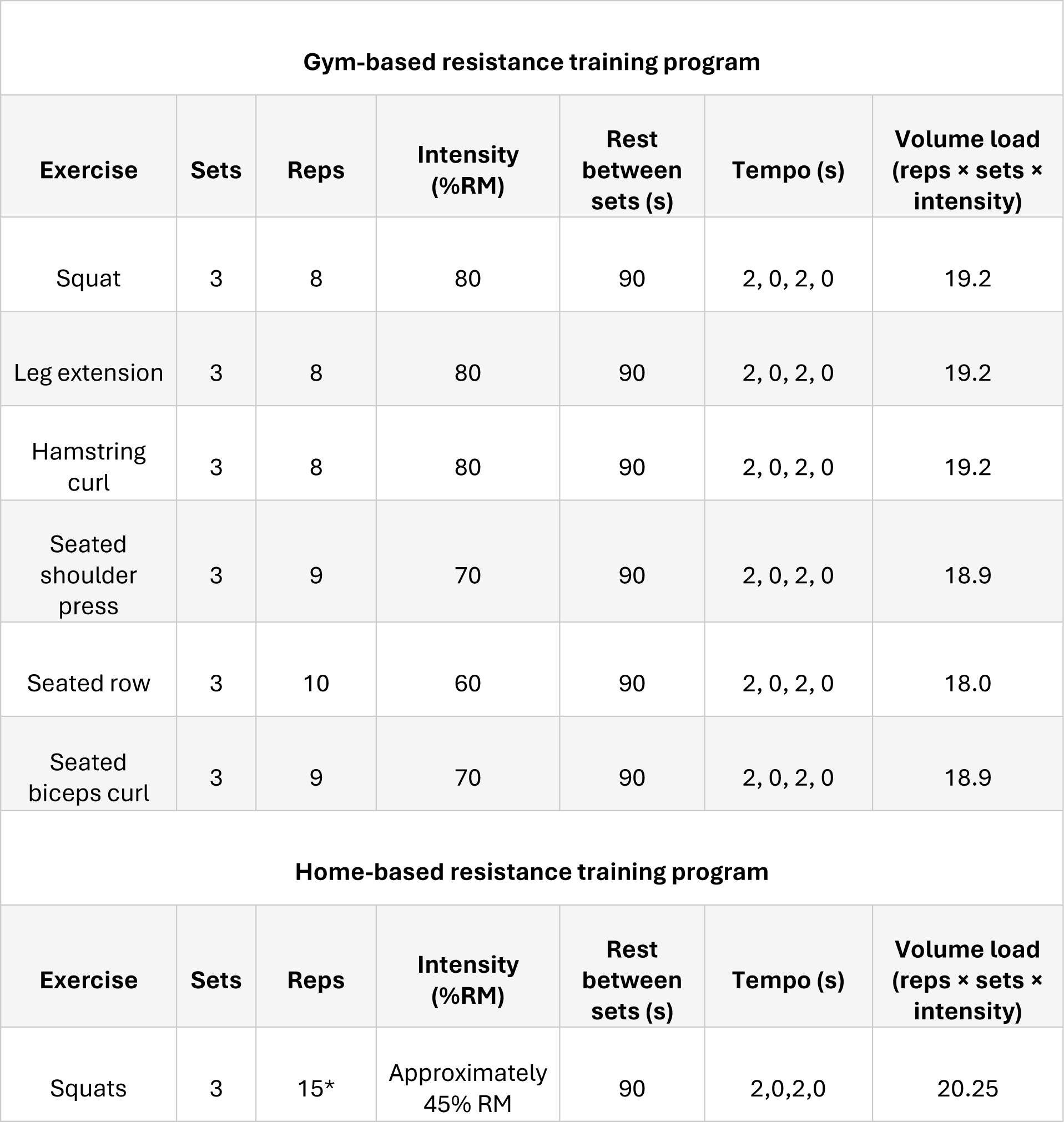
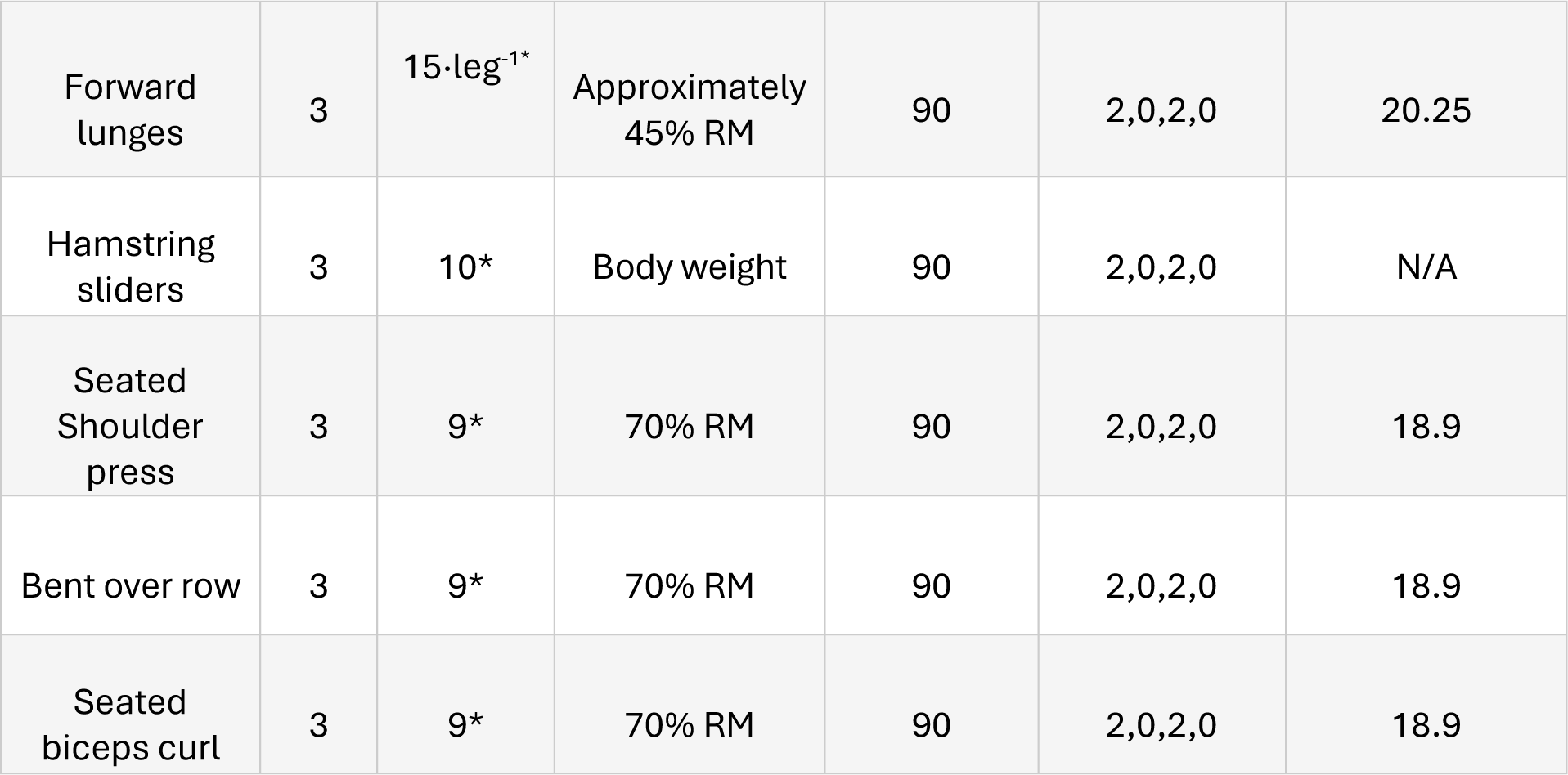
The resistance training program prescribed to participants (n=27). The gym-based program was followed by all participants when access to the gym was possible. A sub-cohort of participants (n=10) performed a portion of their training programs using the home-based resistance training program when access to the training facility was not possible. *last set was prescribed as many repetitions as possible (AMRAP).

**Supplementary Table 3.**
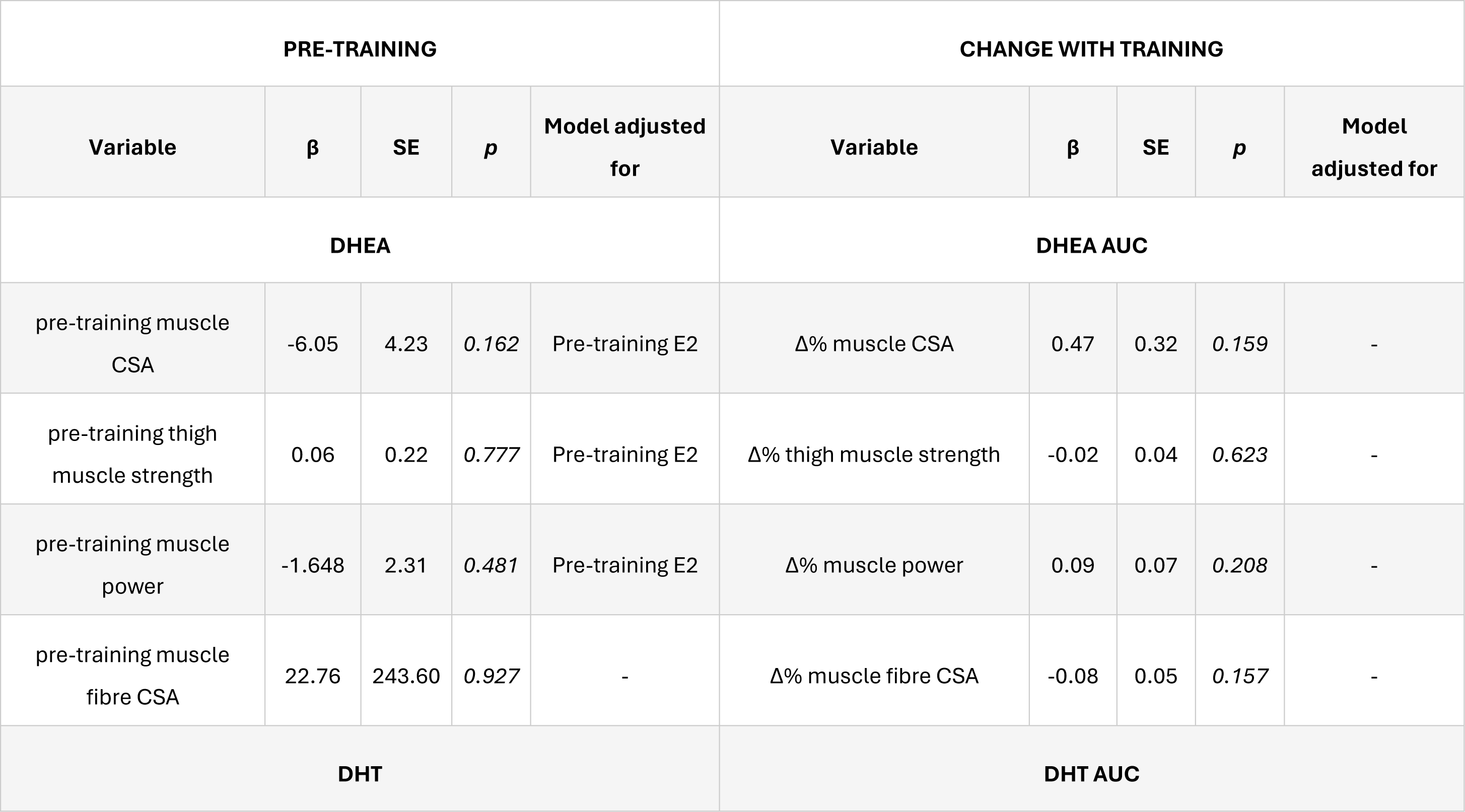
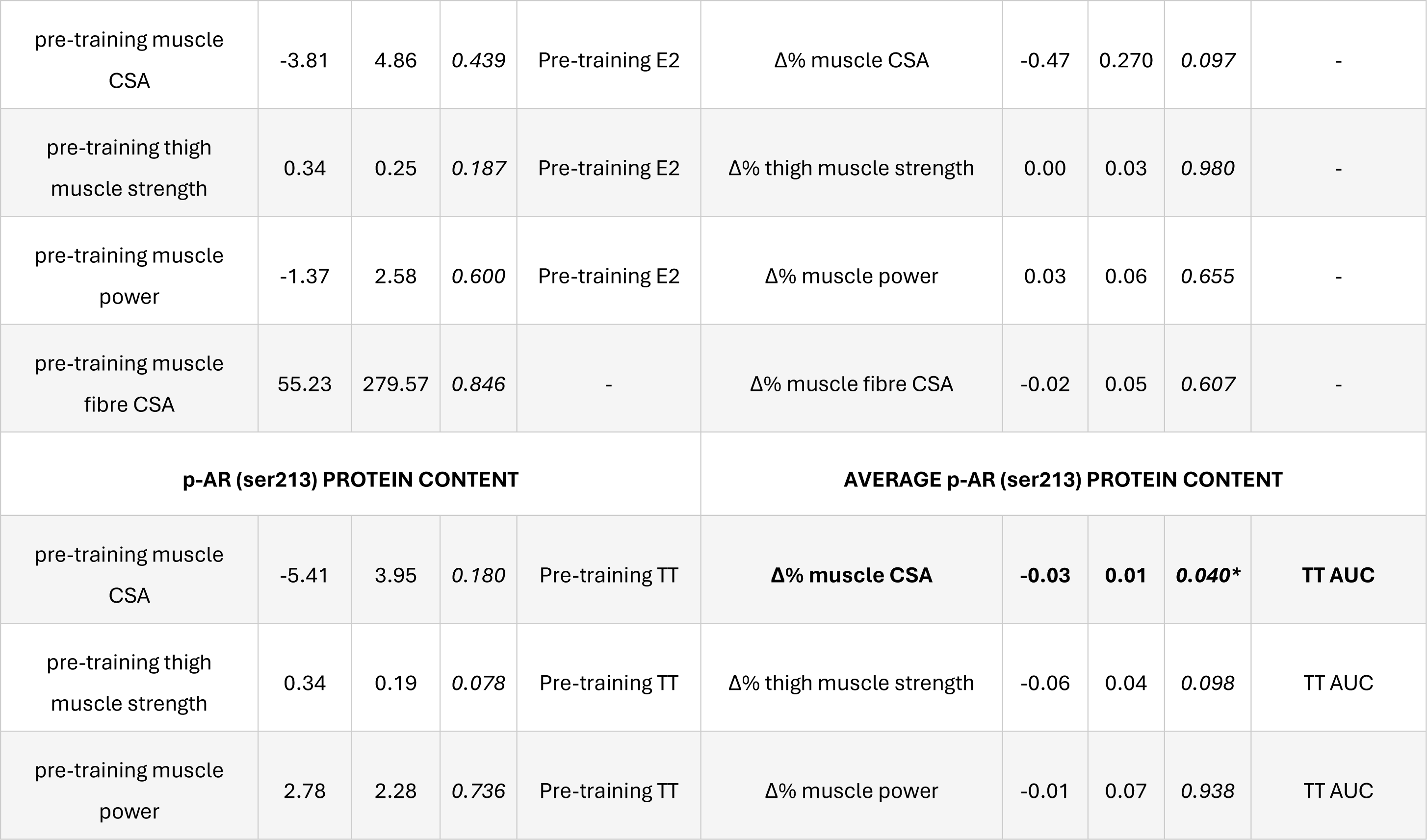
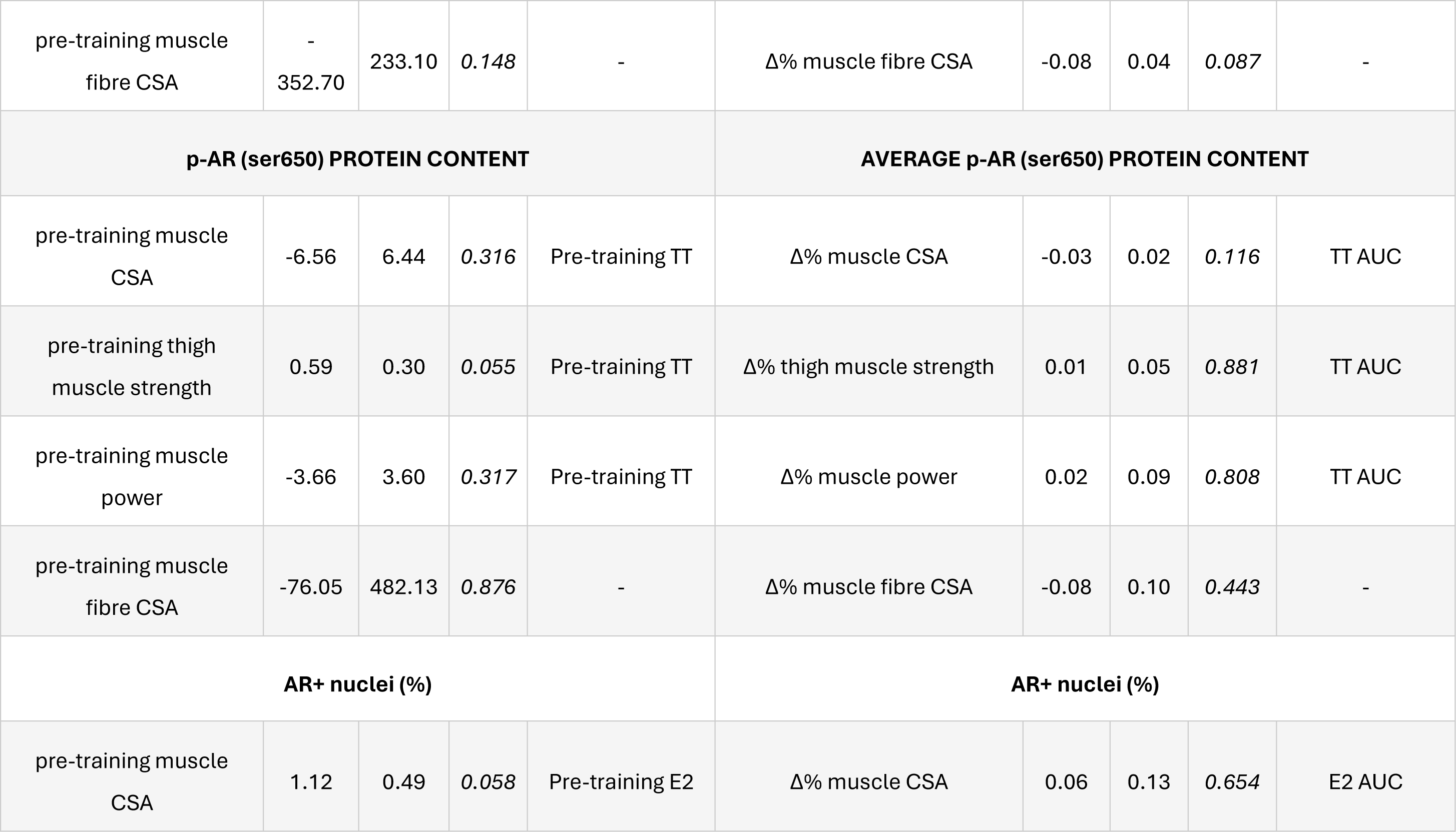
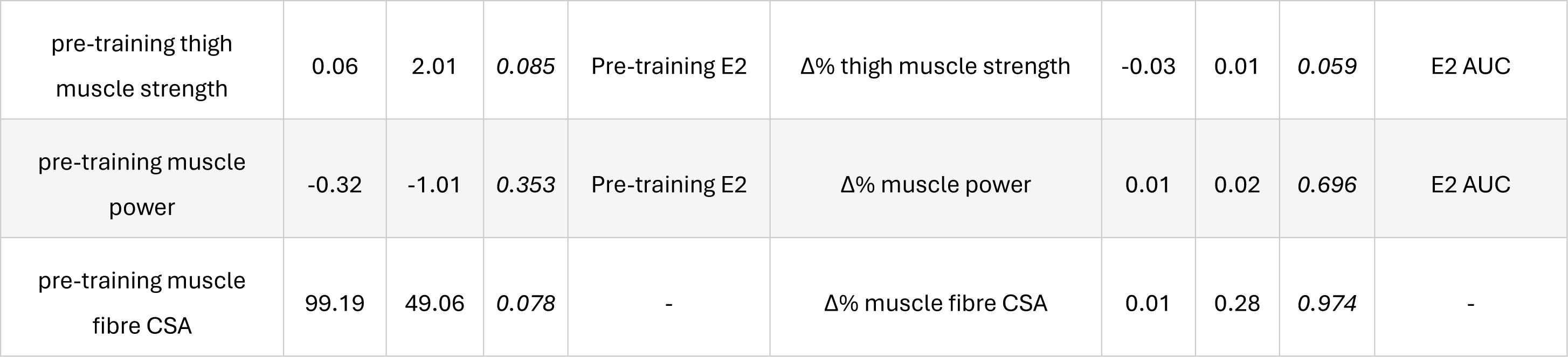
Linear mixed models assessing the association between markers of the androgen profile and AR activity and muscle size, strength and power before and after 12 weeks of resistance training in pre-menopausal females (n=35 pre-training, 27 change with training).

| PRE-TRAINING |  |  |  |  | CHANGE WITH TRAINING |  |  |  |  |
| --- | --- | --- | --- | --- | --- | --- | --- | --- | --- |
| Variable | $\beta$ | SE | *p* | Model adjusted for | Variable | $\beta$ | SE | *p* | Model adjusted for |
| DHEA |  |  |  |  | DHEA AUC |  |  |  |  |
| pre-training muscle CSA | -6.05 | 4.23 | 0.162 | Pre-training E2 | $\Delta\%$ muscle CSA | 0.47 | 0.32 | 0.159 | - |
| pre-training thigh muscle strength | 0.06 | 0.22 | 0.777 | Pre-training E2 | $\Delta\%$ thigh muscle strength | -0.02 | 0.04 | 0.623 | - |
| pre-training muscle power | -1.648 | 2.31 | 0.481 | Pre-training E2 | $\Delta\%$ muscle power | 0.09 | 0.07 | 0.208 | - |
| pre-training muscle fibre CSA | 22.76 | 243.60 | 0.927 | - | $\Delta\%$ muscle fibre CSA | -0.08 | 0.05 | 0.157 | - |
| DHT |  |  |  |  | DHT AUC |  |  |  |  |
| pre-training muscle CSA | -3.81 | 4.86 | 0.439 | Pre-training E2 | $\Delta\%$ muscle CSA | -0.47 | 0.270 | 0.097 | - |
| pre-training thigh muscle strength | 0.34 | 0.25 | 0.187 | Pre-training E2 | $\Delta\%$ thigh muscle strength | 0.00 | 0.03 | 0.980 | - |
| pre-training muscle power | -1.37 | 2.58 | 0.600 | Pre-training E2 | $\Delta\%$ muscle power | 0.03 | 0.06 | 0.655 | - |
| pre-training muscle fibre CSA | 55.23 | 279.57 | 0.846 | - | $\Delta\%$ muscle fibre CSA | -0.02 | 0.05 | 0.607 | - |
| **p-AR (ser213) PROTEIN CONTENT** |  |  |  |  | **AVERAGE p-AR (ser213) PROTEIN CONTENT** |  |  |  |  |
| pre-training muscle CSA | -5.41 | 3.95 | 0.180 | Pre-training TT | **$\Delta\%$ muscle CSA** | **-0.03** | **0.01** | **0.040\*** | **TT AUC** |
| pre-training thigh muscle strength | 0.34 | 0.19 | 0.078 | Pre-training TT | $\Delta\%$ thigh muscle strength | -0.06 | 0.04 | 0.098 | TT AUC |
| pre-training muscle power | 2.78 | 2.28 | 0.736 | Pre-training TT | $\Delta\%$ muscle power | -0.01 | 0.07 | 0.938 | TT AUC |
| pre-training muscle<br>fibre CSA | -<br>352.70 | 233.10 | 0.148 | - | $\Delta\%$ muscle fibre CSA | -0.08 | 0.04 | 0.087 | - |
| **p-AR (ser650) PROTEIN CONTENT** |  |  |  |  | **AVERAGE p-AR (ser650) PROTEIN CONTENT** |  |  |  |  |
| pre-training muscle<br>CSA | -6.56 | 6.44 | 0.316 | Pre-training TT | $\Delta\%$ muscle CSA | -0.03 | 0.02 | 0.116 | TT AUC |
| pre-training thigh<br>muscle strength | 0.59 | 0.30 | 0.055 | Pre-training TT | $\Delta\%$ thigh muscle strength | 0.01 | 0.05 | 0.881 | TT AUC |
| pre-training muscle<br>power | -3.66 | 3.60 | 0.317 | Pre-training TT | $\Delta\%$ muscle power | 0.02 | 0.09 | 0.808 | TT AUC |
| pre-training muscle<br>fibre CSA | -76.05 | 482.13 | 0.876 | - | $\Delta\%$ muscle fibre CSA | -0.08 | 0.10 | 0.443 | - |
| **AR+ nuclei (%)** |  |  |  |  | **AR+ nuclei (%)** |  |  |  |  |
| pre-training muscle<br>CSA | 1.12 | 0.49 | 0.058 | Pre-training E2 | $\Delta\%$ muscle CSA | 0.06 | 0.13 | 0.654 | E2 AUC |
| pre-training thigh muscle strength | 0.06 | 2.01 | 0.085 | Pre-training E2 | $\Delta\%$ thigh muscle strength | -0.03 | 0.01 | 0.059 | E2 AUC |
| pre-training muscle power | -0.32 | -1.01 | 0.353 | Pre-training E2 | $\Delta\%$ muscle power | 0.01 | 0.02 | 0.696 | E2 AUC |
| pre-training muscle fibre CSA | 99.19 | 49.06 | 0.078 | - | $\Delta\%$ muscle fibre CSA | 0.01 | 0.28 | 0.974 | - |

