## Supplementary table 3 for "Total testosterone is not associated with muscle mass, function or exercise adaptations in pre-menopausal females"

**Supplementary Table 3**. Linear mixed models assessing the relationship between markers of the androgen profile and AR activity and muscle size, strength and power before and after 12 weeks of resistance training in pre-menopausal females (n=35 pre-training, 27 change with training).

| PRE-TRAINING | | | | | CHANGE WITH TRAINING | | | | |
| --- | --- | --- | --- | --- | --- | --- | --- | --- | --- |
| Variable | **β** | **SE** | ***p*** | **Model adjusted for** | **Variable** | **β** | **SE** | ***p*** | **Model adjusted for** |
| DHEA | | | | | **DHEA AUC** | | | | |
| pre-training muscle CSA | -6.05 | 4.23 | *0.162* | Pre-training E2 | ∆ muscle CSA | 0.47 | 0.32 | *0.159* | - |
| pre-training thigh muscle strength | 0.06 | 0.22 | *0.777* | Pre-training E2 | ∆ thigh muscle strength | -0.02 | 0.04 | *0.623* | - |
| pre-training muscle power | -1.648 | 2.31 | *0.481* | Pre-training E2 | ∆ muscle power | 0.09 | 0.07 | *0.208* | - |
| pre-training muscle fibre CSA | 22.76 | 243.60 | *0.927* | - | ∆ muscle fibre CSA | -0.08 | 0.05 | *0.157* | - |
| DHT | | | | | **DHT AUC** | | | | |
| pre-training muscle CSA | -3.81 | 4.86 | *0.439* | Pre-training E2 | ∆ muscle CSA | -0.47 | 0.270 | *0.097* | - |
| pre-training thigh muscle strength | 0.34 | 0.25 | *0.187* | Pre-training E2 | ∆ thigh muscle strength | 0.00 | 0.03 | *0.980* | - |
| pre-training muscle power | -1.37 | 2.58 | *0.600* | Pre-training E2 | ∆ muscle power | 0.03 | 0.06 | *0.655* | - |
| pre-training muscle fibre CSA | 55.23 | 279.57 | *0.846* | - | ∆ muscle fibre CSA | -0.02 | 0.05 | *0.607* | - |
| AR PROTEIN CONTENT | | | | | **AVERAGE AR PROTEIN CONTENT** | | | | |
| pre-training muscle CSA | **-25.99** | **7.98** | ***0.003**** | **Pre-training TT** | ∆ muscle CSA | 0.01 | 0.04 | *0.202* | TT AUC |
| pre-training thigh muscle strength | 0.26 | 0.44 | *0.561* | Pre-training TT | ∆ thigh muscle strength | 0.01 | 0.09 | *0.943* | TT AUC |
| pre-training muscle power | 5.45 | 5.09 | *0.292* | Pre-training TT | ∆ muscle power | -0.08 | 0.194 | *0.688* | TT AUC |
| pre-training muscle fibre CSA | -552.7 | 522.9 | *0.305* | - | ∆ muscle fibre CSA | -0.02 | 0.13 | *0.081* | - |
| p-AR (ser213) PROTEIN CONTENT | | | | | **AVERAGE p-AR (ser213) PROTEIN CONTENT** | | | | |
| pre-training muscle CSA | -5.41 | 3.95 | *0.180* | Pre-training TT | **∆ muscle CSA** | **-0.03** | **0.01** | ***0.040**** | **TT AUC** |
| pre-training thigh muscle strength | 0.34 | 0.19 | *0.078* | Pre-training TT | ∆ thigh muscle strength | -0.06 | 0.04 | *0.098* | TT AUC |
| pre-training muscle power | 2.78 | 2.28 | *0.736* | Pre-training TT | ∆ muscle power | -0.01 | 0.07 | *0.938* | TT AUC |
| pre-training muscle fibre CSA | -352.70 | 233.10 | *0.148* | - | ∆ muscle fibre CSA | -0.08 | 0.04 | *0.087* | - |
| p-AR (ser650) PROTEIN CONTENT | | | | | **AVERAGE p-AR (ser650) PROTEIN CONTENT** | | | | |
| pre-training muscle CSA | -6.56 | 6.44 | *0.316* | Pre-training TT | ∆ muscle CSA | -0.03 | 0.02 | *0.116* | TT AUC |
| pre-training thigh muscle strength | 0.59 | 0.30 | *0.055* | Pre-training TT | ∆ thigh muscle strength | 0.01 | 0.05 | *0.881* | TT AUC |
| pre-training muscle power | -3.66 | 3.60 | *0.317* | Pre-training TT | ∆ muscle power | 0.02 | 0.09 | *0.808* | TT AUC |
| pre-training muscle fibre CSA | -76.05 | 482.13 | *0.876* | - | ∆ muscle fibre CSA | -0.08 | 0.10 | *0.443* | - |
| AR+ nuclei (%) | | | | | **AR+ nuclei (%)** | | | | |
| pre-training muscle CSA | 1.12 | 0.49 | *0.058* | Pre-training E2 | ∆ muscle CSA | 0.06 | 0.13 | *0.654* | E2 AUC |
| pre-training thigh muscle strength | 0.06 | 2.01 | *0.085* | Pre-training E2 | ∆ thigh muscle strength | -0.03 | 0.01 | *0.059* | E2 AUC |
| pre-training muscle power | -0.32 | -1.01 | *0.353* | Pre-training E2 | ∆ muscle power | 0.01 | 0.02 | *0.696* | E2 AUC |
| pre-training muscle fibre CSA | 99.19 | 49.06 | *0.078* | - | ∆ muscle fibre CSA | 0.01 | 0.28 | *0.974* | - |
