## Supplementary table 4 for "Total testosterone is not associated with muscle mass, function or exercise adaptations in pre-menopausal females"

**Supplementary Table 4**. Linear mixed models.

| PRE-TRAINING | | | | | CHANGE WITH TRAINING | | | | |
| --- | --- | --- | --- | --- | --- | --- | --- | --- | --- |
| Variable | **β** | **SE** | ***p*** | **Model adjusted for** | **Variable** | **β** | **SE** | ***p*** | **Model adjusted for** |
| TOTAL TESTOSTERONE (TT) | | | | | **TESTOSTERONE AUC** | | | | |
| pre-training muscle CSA | -5.18 | 6.30 | *0.417* | Pre-training E2 | ∆ muscle CSA | 0.00 | 0.00 | *0.207* | E2 AUC |
| pre-training thigh muscle strength | 0.25 | 0.32 | *0.445* | Pre-training E2 | ∆ thigh muscle strength | 0.00 | 0.01 | *0.744* | E2 AUC |
| pre-training muscle power | 0.30 | 3.44 | *0.929* | Pre-training E2 | ∆ muscle power | 0.02 | 0.02 | *0.279* | - |
| pre-training muscle fibre CSA | 514.20 | 336.10 | *0.147* | - | ∆ muscle fibre CSA | 0.08 | 0.12 | *0.534* | - |
| pre-training total AR | 0.13 | 0.12 | *0.246* | *­-* | Average AR | 0.01 | 0.02 | *0.672* | *-* |
| pre-training p-AR^ser213^ | 0.05 | 0.24 | *0.846* | *-* | Average p-AR^ser213^ | 0.00 | 0.05 | *0.991* | *-* |
| pre-training p-AR^ser650^ | 0.18 | 0.15 | *0.249* | *-* | Average p-AR^ser650^ | 0.01 | 0.05 | *0.881* | *-* |
| DHEA | | | | | **DHEA AUC** | | | | |
| pre-training muscle CSA | -6.05 | 4.23 | *0.162* | Pre-training E2 | ∆ muscle CSA | 0.47 | 0.32 | *0.159* | - |
| pre-training thigh muscle strength | 0.06 | 0.22 | *0.777* | Pre-training E2 | ∆ thigh muscle strength | -0.02 | 0.04 | *0.623* | - |
| pre-training muscle power | -1.648 | 2.31 | *0.481* | Pre-training E2 | ∆ muscle power | 0.09 | 0.07 | *0.208* | - |
| pre-training muscle fibre CSA | 22.76 | 243.60 | *0.927* | - | ∆ muscle fibre CSA | -0.08 | 0.05 | *0.157* | - |
| DHT | | | | | **DHT AUC** | | | | |
| pre-training muscle CSA | -3.81 | 4.86 | *0.439* | Pre-training E2 | ∆ muscle CSA | -0.47 | 0.270 | *0.097* | - |
| pre-training thigh muscle strength | 0.34 | 0.25 | *0.187* | Pre-training E2 | ∆ thigh muscle strength | 0.00 | 0.03 | *0.980* | - |
| pre-training muscle power | -1.37 | 2.58 | *0.600* | Pre-training E2 | ∆ muscle power | 0.03 | 0.06 | *0.655* | - |
| pre-training muscle fibre CSA | 55.23 | 279.57 | *0.846* | - | ∆ muscle fibre CSA | -0.02 | 0.05 | *0.607* | - |
| FREE ANDROGEN INDEX (FAI) | | | | | **FREE ANDROGEN INDEX AUC** | | | | |
| pre-training muscle CSA | -1.05 | 3.05 | *0.734* | Pre-training E2 | ∆ muscle CSA | 0.00 | 0.01 | *0.799* | E2 AUC |
| pre-training thigh muscle strength | 0.29 | 0.15 | *0.095* | Pre-training E2 | **∆ thigh muscle strength** | **0.05** | **0.02** | ***0.044**** | E2 AUC |
| pre-training muscle power | 0.98 | 1.33 | *0.469* | - | ∆ muscle power | -0.01 | 0.05 | *0.912* | E2 AUC |
| pre-training muscle fibre CSA | 2.94 | 159.2 | *0.985* | - | ∆ muscle fibre CSA | 0.03 | 0.03 | *0.378* | E2 AUC |
| pre-training total AR | -0.05 | 0.05 | *0.336* | *­-* | Average AR | -0.03 | 0.05 | *0.595* | *­-* |
| pre-training p-AR^ser213^ | -0.04 | 0.12 | *0.708* | *-* | Average p-AR^ser213^ | 0.00 | 0.12 | *0.989* | *-* |
| pre-training p-AR^ser650^ | 0.13 | 0.07 | *0.072* | *-* | **Average p-AR^ser650^** | **0.22** | **0.09** | ***0.020**** | *-* |
| OESTRADIOL (E2) | | | | | **OESTRADIOL AUC** | | | | |
| pre-training muscle CSA | -0.38 | 2.87 | *0.897* | Menstrual phase at PRE testing | ∆ muscle CSA | 0.40 | 0.33 | *0.252* | - |
| pre-training thigh muscle strength | 0.3. | 0.27 | *0.280* | Menstrual phase at PRE testing | ∆ thigh muscle strength | 0.02 | 0.02 | *0.327* | **-** |
| pre-training muscle power | 1.309 | 3.00 | *0.668* | Menstrual phase at PRE testing | ∆ muscle power | -0.02 | 0.05 | *0.672* | - |
| pre-training muscle fibre CSA | -61.25 | 206.26 | *0.771* | - | **∆ muscle fibre CSA** | **-0.09** | **0.03** | ***0.013**** | - |
| PROGESTERONE (P) | | | | | **PROGESTERONE AUC** | | | | |
| pre-training muscle CSA | -6.69 | 3.77 | *0.095* | Menstrual phase at PRE testing | ∆ muscle CSA | 0.11 | 0.34 | *0.744* | E2 AUC |
| pre-training thigh muscle strength | 0.70 | 0.36 | *0.072* | Menstrual phase at PRE testing | ∆ thigh muscle strength | -0.02 | 0.03 | *0.550* | E2 AUC |
| pre-training muscle power | 0.79 | 4.34 | *0.857* | Menstrual phase at PRE testing | ∆ muscle power | -0.06 | 0.06 | *0.349* | - |
| pre-training muscle fibre CSA | -152.8 | 382.8 | *0.696* | - | ∆ muscle fibre CSA | -0.04 | 0.06 | *0.570* | - |
| AR PROTEIN CONTENT | | | | | **AVERAGE AR PROTEIN CONTENT** | | | | |
| pre-training muscle CSA | **-25.99** | **7.98** | ***0.003**** | **Pre-training TT** | ∆ muscle CSA | 0.01 | 0.04 | *0.202* | TT AUC |
| pre-training thigh muscle strength | 0.26 | 0.44 | *0.561* | Pre-training TT | ∆ thigh muscle strength | 0.01 | 0.09 | *0.943* | TT AUC |
| pre-training muscle power | 5.45 | 5.09 | *0.292* | Pre-training TT | ∆ muscle power | -0.08 | 0.194 | *0.688* | TT AUC |
| pre-training muscle fibre CSA | -552.7 | 522.9 | *0.305* | - | ∆ muscle fibre CSA | -0.02 | 0.13 | *0.081* | - |
| p-AR (ser213) PROTEIN CONTENT | | | | | **AVERAGE p-AR (ser213) PROTEIN CONTENT** | | | | |
| pre-training muscle CSA | -5.41 | 3.95 | *0.180* | Pre-training TT | **∆ muscle CSA** | **-0.03** | **0.01** | ***0.040**** | **TT AUC** |
| pre-training thigh muscle strength | 0.34 | 0.19 | *0.078* | Pre-training TT | ∆ thigh muscle strength | -0.06 | 0.04 | *0.098* | TT AUC |
| pre-training muscle power | 2.78 | 2.28 | *0.736* | Pre-training TT | ∆ muscle power | -0.01 | 0.07 | *0.938* | TT AUC |
| pre-training muscle fibre CSA | -352.70 | 233.10 | *0.148* | - | ∆ muscle fibre CSA | -0.08 | 0.04 | *0.087* | - |
| p-AR (ser650) PROTEIN CONTENT | | | | | **AVERAGE p-AR (ser650) PROTEIN CONTENT** | | | | |
| pre-training muscle CSA | -6.56 | 6.44 | *0.316* | Pre-training TT | ∆ muscle CSA | -0.03 | 0.02 | *0.116* | TT AUC |
| pre-training thigh muscle strength | 0.59 | 0.30 | *0.055* | Pre-training TT | ∆ thigh muscle strength | 0.01 | 0.05 | *0.881* | TT AUC |
| pre-training muscle power | -3.66 | 3.60 | *0.317* | Pre-training TT | ∆ muscle power | 0.02 | 0.09 | *0.808* | TT AUC |
| pre-training muscle fibre CSA | -76.05 | 482.13 | *0.876* | - | ∆ muscle fibre CSA | -0.08 | 0.10 | *0.443* | - |
