## Supplementary figures and images for "Total testosterone is not associated with muscle mass, function or exercise adaptations in pre-menopausal females"

### Supplementary figure 1

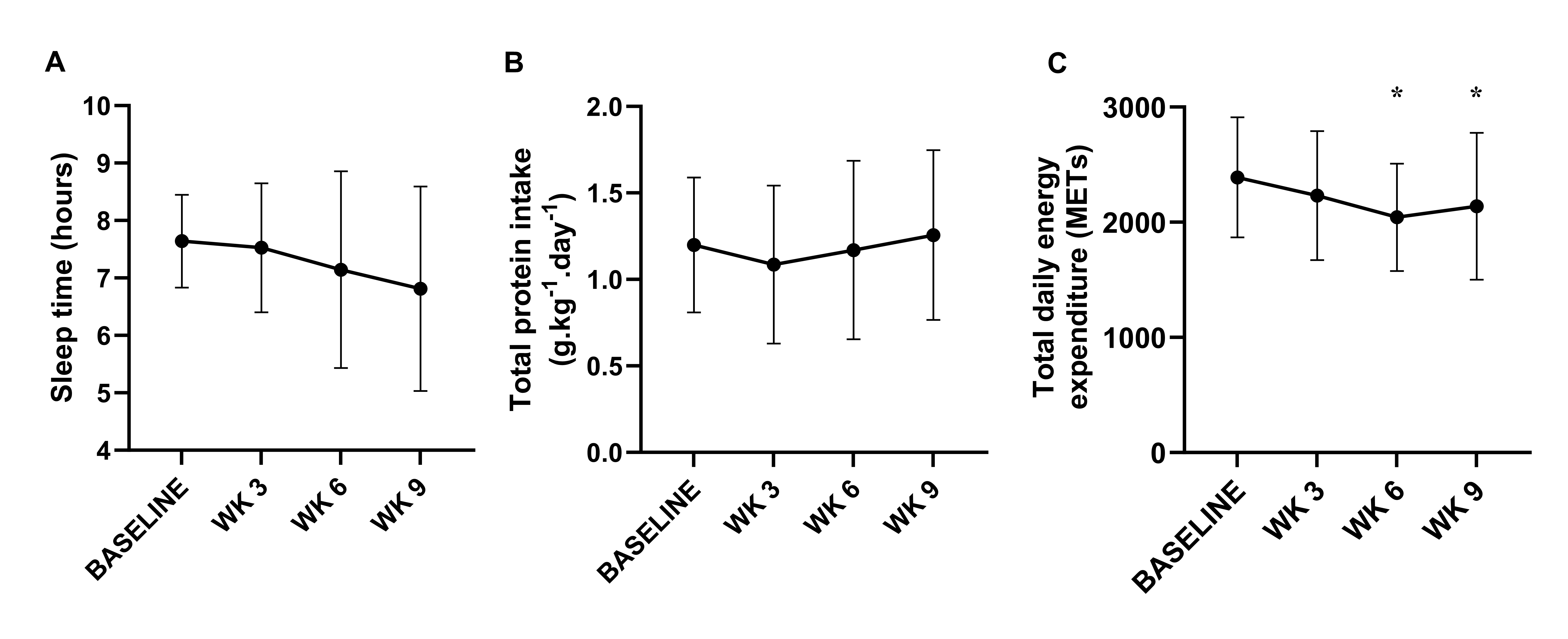

### supplementary figure 2

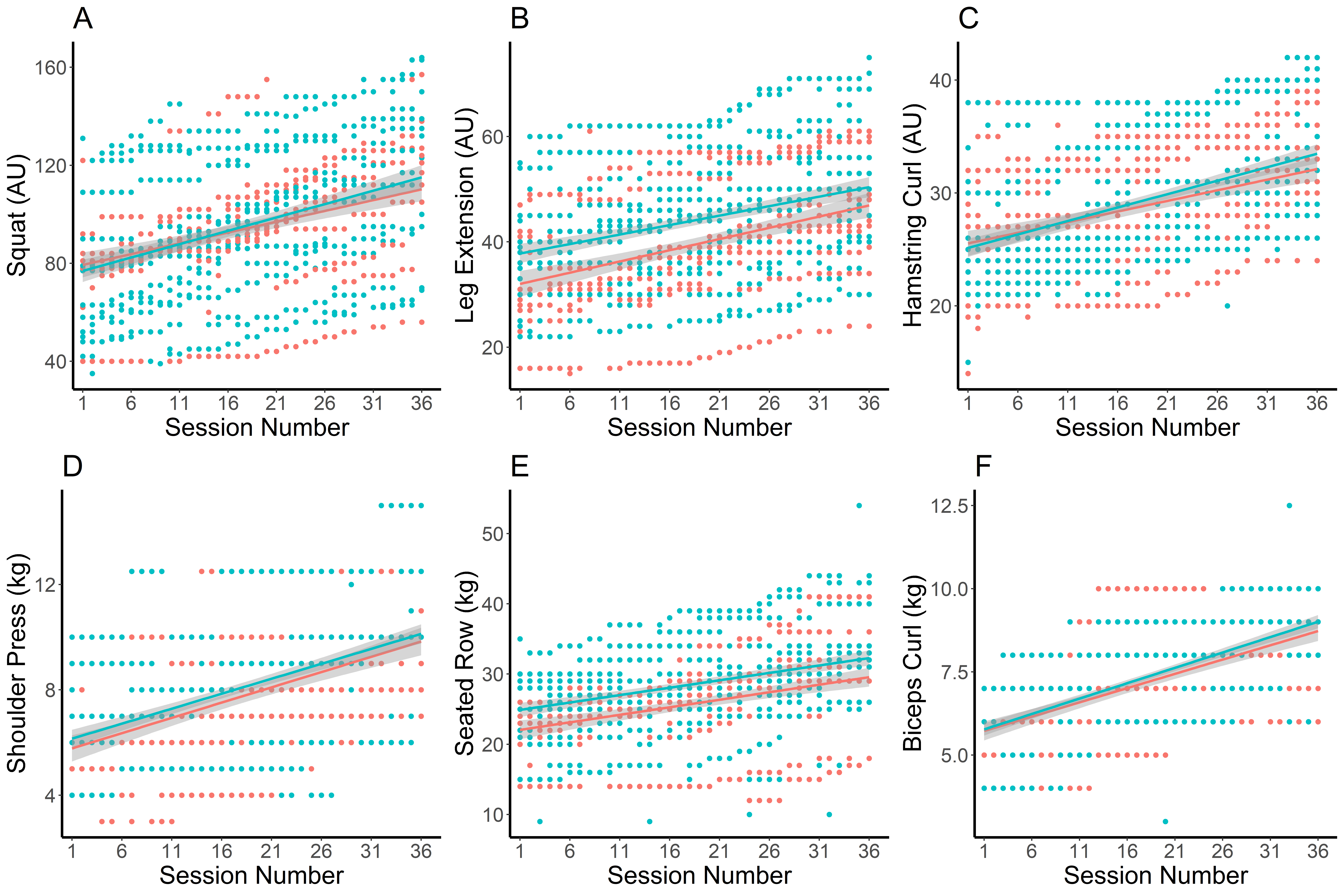

### Supplementary figure 3

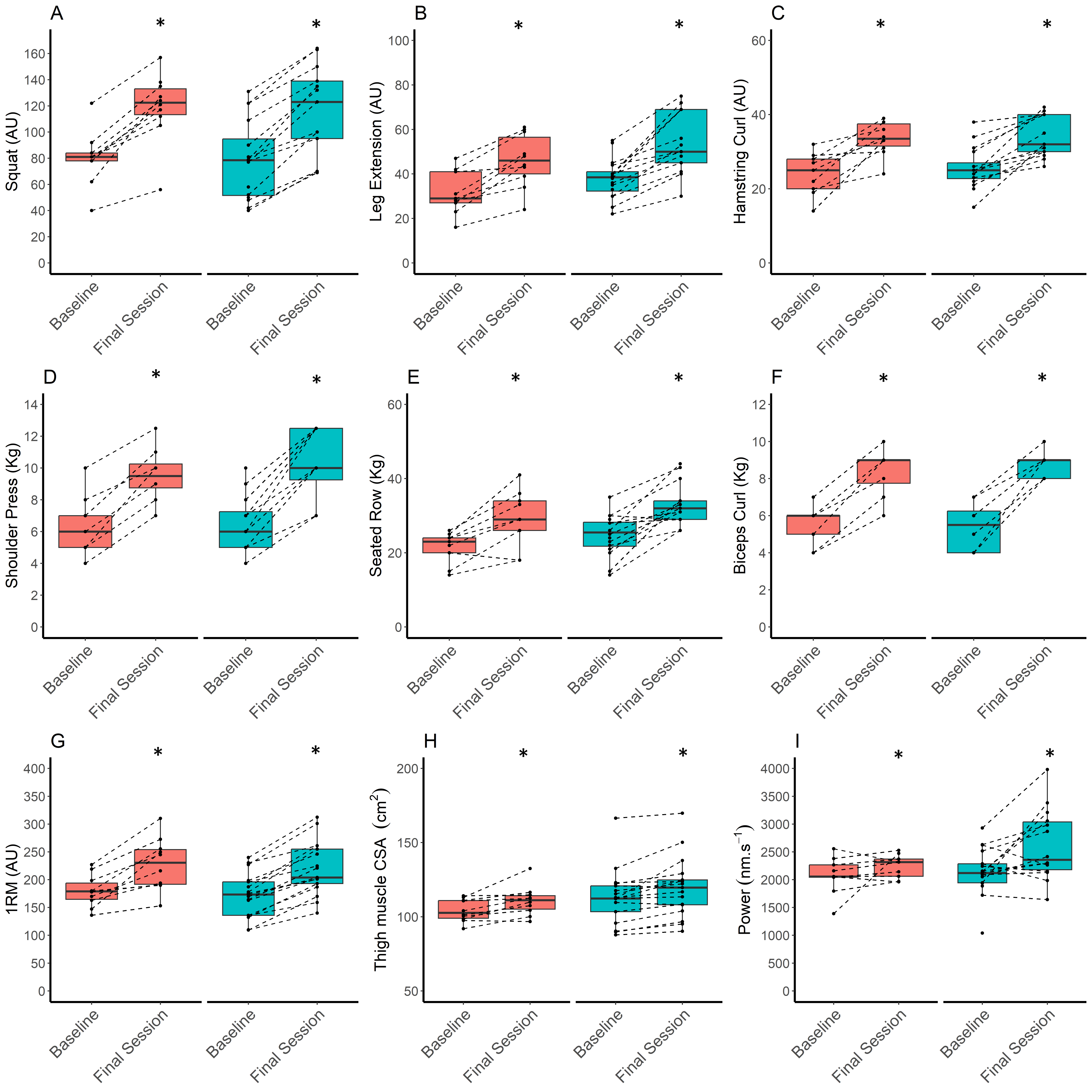

### supplementary figure 4

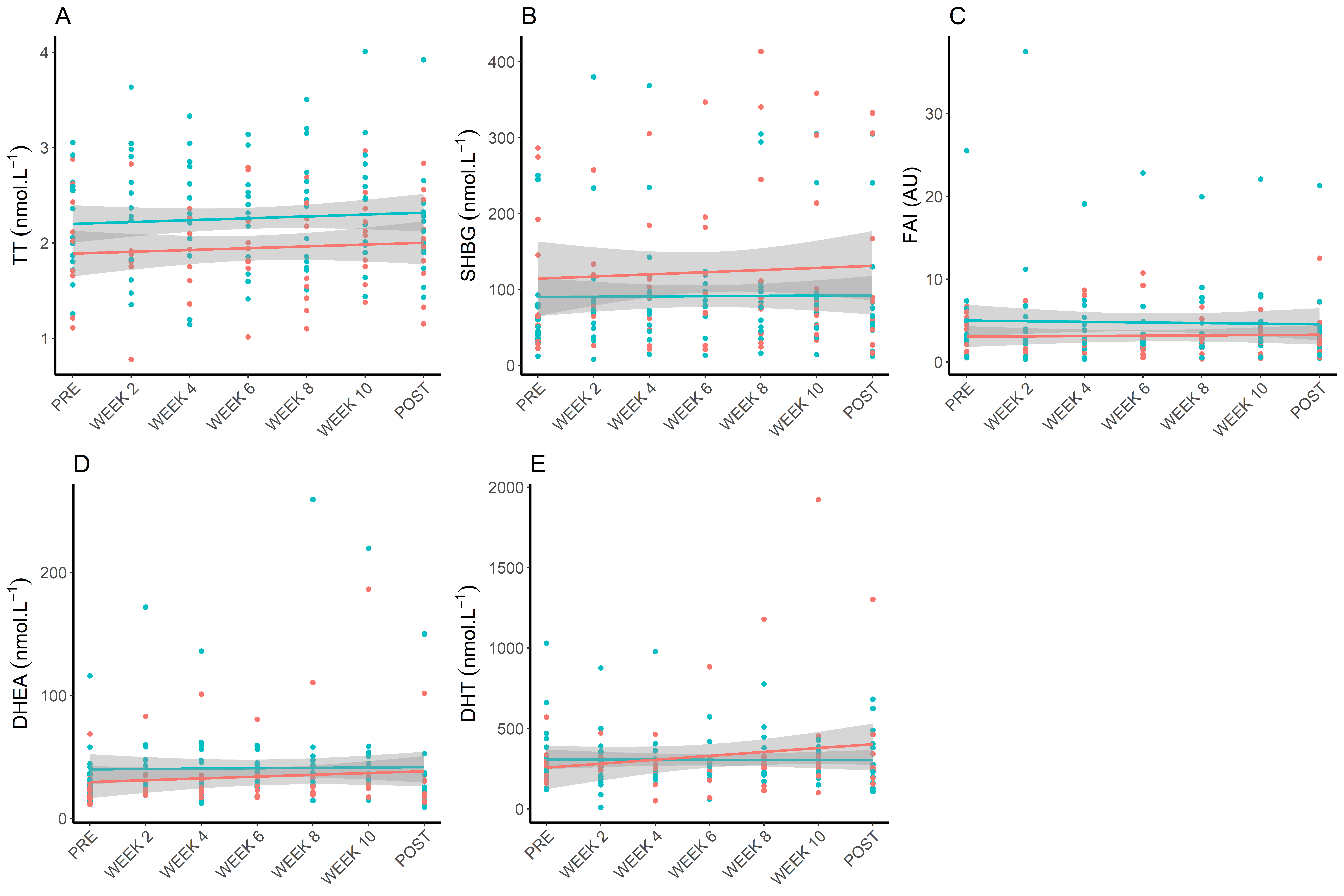

### Supplementary figure 5

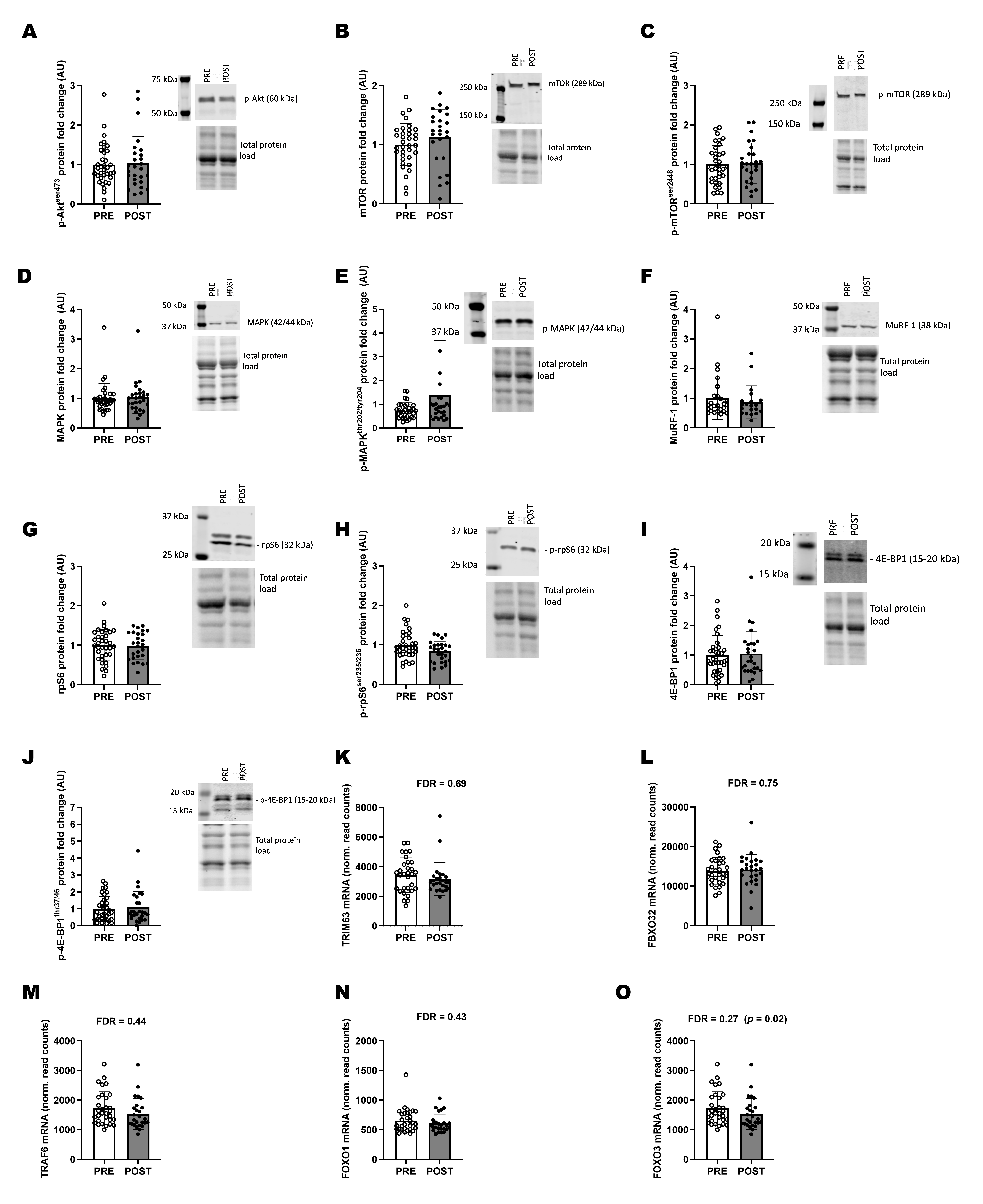

### Supplementary figure 6

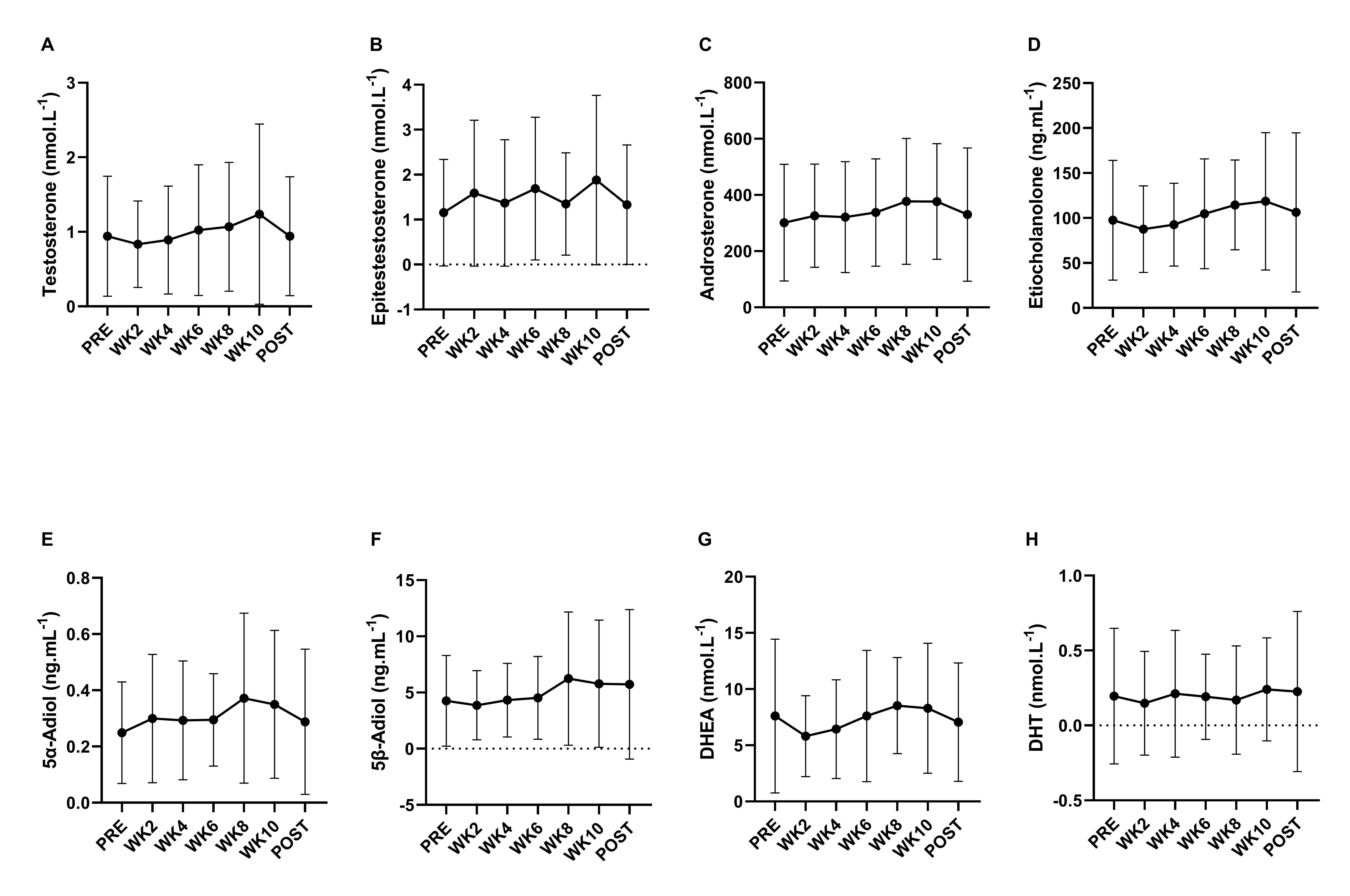

### Supplementary figure 7

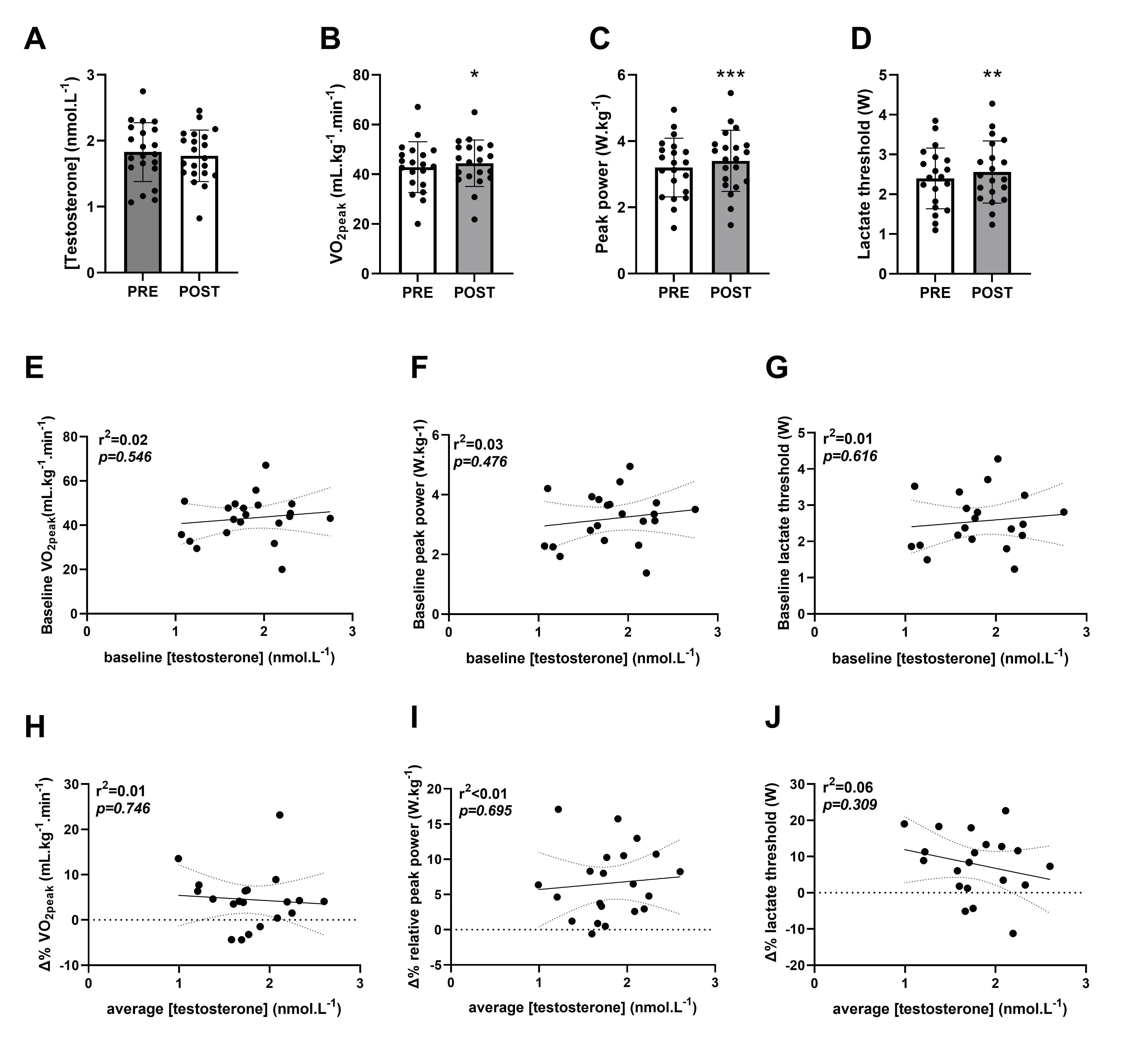

### Supplementary figure 8

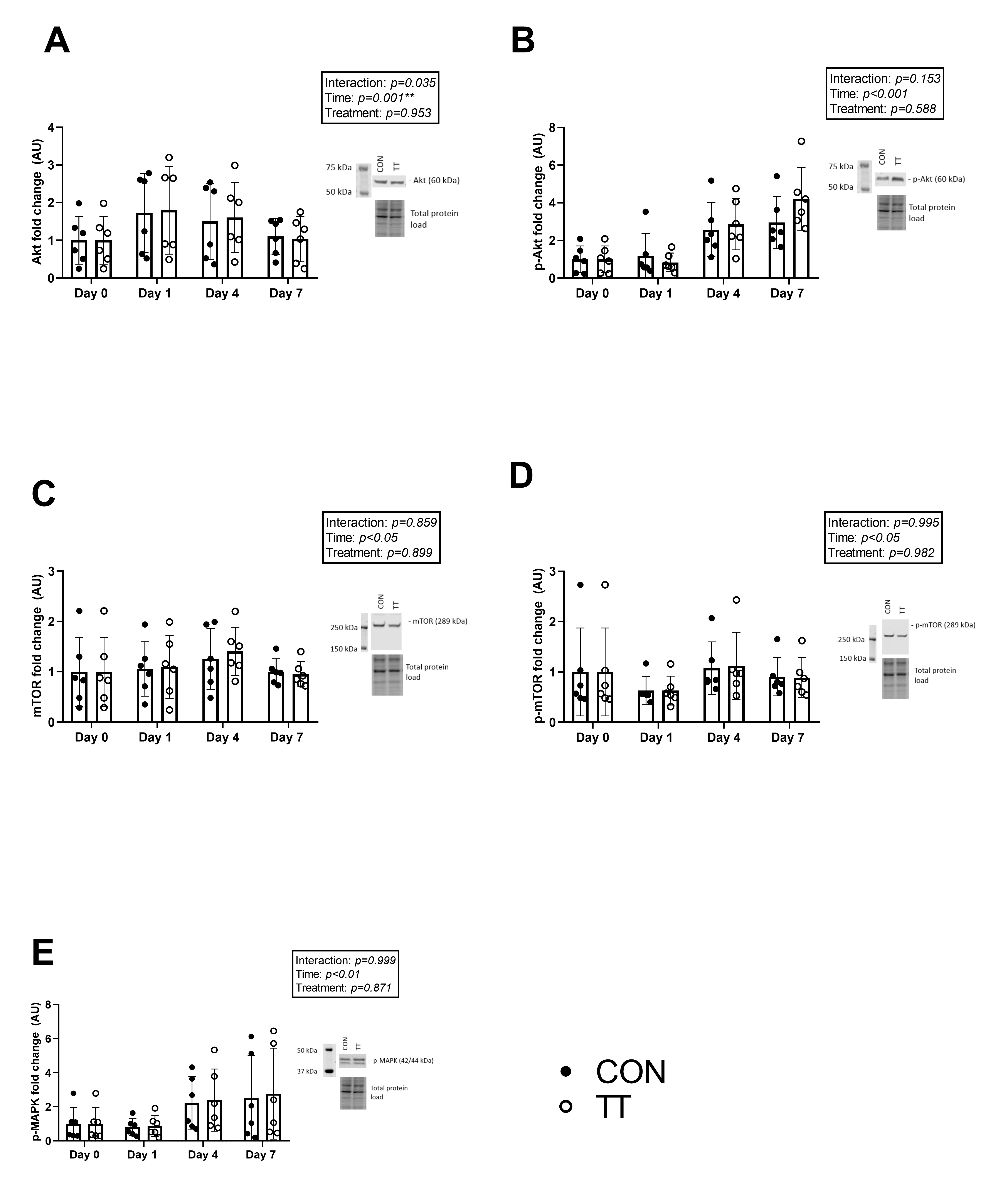
