## Supplementary table 1 for "Total testosterone is not associated with muscle mass, function or exercise adaptations in pre-menopausal females"

**Supplementary Table 1.** Summary of the menstrual phases of each participant at pre- and post-training testing timepoints. The menstrual phase was informed by a menstrual calendar collected by the participant and circulating concentrations of oestradiol (E2), progesterone, luteinising hormone (LH) and follicle stimulating hormone (FSH). Two independent researchers determined the menstrual phase for each entry and agreement was met at each case. Note: This study aimed to avoid testing and biopsy times during the late follicular (LF) phase of the menstrual cycle. Seven out of 62 (11%) testing times were taken during the late follicular phase. Statistical analyses will include the menstrual phase as a covariate to account for these times.

| **Participant** | **Timepoint** | **Menstrual calendar timepoint** | **E2 (pmol·L^-1^)** | **Progesterone (nmol·L^-1^)** | **LH (IU·L^-1^)** | **FSH (IU·L^-1^)** | **Menstrual phase (informed by calendar and hormones)** |
| --- | --- | --- | --- | --- | --- | --- | --- |
| P01 | PRE | Day 11 (out of 29) | 514.1 | 6.7 | 5.3 | 4.5 | **LF** |
| P01 | POST | Day 18 (out of 26) | 325.6 | 7.81 | 7.2 | 5.8 | EL |
| P02 | PRE | N/A | 96.6 | 3.82 | 8.2 | 14 | EF |
| P02 | POST | N/A | 169.5 | 4.337 | 1.2 | 5.7 | EF |
| P03 | PRE | N/A | 92.3 | 7.62 | 9.6 | 8.3 | ACTIVE PILL |
| P03 | POST | N/A | 28.3 | 7.3 | 2.4 | 2.9 | ACTIVE PILL |
| P04 | PRE | N/A | 106.3 | 4.56 | 1.7 | 4.9 | ACTIVE PILL |
| P04 | POST | N/A | 49.3 | 2.97 | 4.4 | 7.3 | ACTIVE PILL |
| P05 | PRE | N/A | 111.6 | 4.36 | 3.6 | 8 | ACTIVE PILL |
| P05 | POST | N/A | 127.1 | 4.58 | 3.1 | 7.5 | ACTIVE PILL |
| P06 | PRE | N/A | 30.4 | 3.54 | 0 | 0.3 | ACTIVE PILL |
| P06 | POST | N/A | 16.3 | 3.02 | 0 | 0.4 | ACTIVE PILL |
| P07 | PRE | Day 2 (out of 43) | 158.3 | 5.11 | 6.5 | 7 | EF |
| P07 | POST | Day 22 (out of 30) | 112.9 | 5.36 | 21.7 | 7.2 | EL |
| P08 | PRE | Day 31 (out of 34) | 93.8 | 10.73 | 7.4 | 7 | ML |
| P08 | POST | Day 33 (out of 35) | 526.1 | 67.83 | 6.8 | 2.2 | ML |
| P09 | PRE | Day 9 (out of 24) | 287.4 | 10.13 | 3.6 | 5.9 | **LF** |
| P09 | POST | Day 22 (out of 27) | 157.6 | 8.33 | 2.6 | 7.6 | ML |
| P10 | PRE | N/A | 95.2 | 5 | 7.1 | 13.3 | EF |
| P10 | POST | N/A | 433.7 | 64.5 | 1.1 | 3.9 | ML |
| P11 | PRE | Day 7 (out of 31) | 248 | 6.44 | 7.2 | 4.7 | **LF** |
| P11 | POST | Day 26 (out of 29) | 159.3 | 9.84 | 6.8 | 6 | ML |
| P12 | PRE | N/A | 63.2 | 9.3 | 0 | 0.1 | ACTIVE PILL |
| P12 | POST | N/A | 81.6 | 9.2 | 0 | 0 | ACTIVE PILL |
| P13 | PRE | Day 27 (out of 34) | 955.5 | 14.7 | 71.1 | 10.6 | EL |
| P13 | POST | Day 14 (out of 30) | 151 | 5.3 | 7.7 | 9.4 | EF |
| P14 | PRE | N/A | 110 | 6.6 | 4.2 | 2 | ACTIVE PILL |
| P14 | POST | N/A | 51.6 | 8.7 | 0 | 0.2 | ACTIVE PILL |
| P15 | PRE | Day 7 (out of 21) | 215.3 | 8.5 | 4.7 | 6 | **LF** |
| P15 | POST | Day 20 (out of 22) | 254.3 | 8.1 | 4.2 | 5.4 | ML |
| P16 | PRE | Day 48 (out of 84) | 162.4 | 4.4 | 2 | 5.5 | EF |
| P16 | POST | Day 22 (out of 43) | 218.7 | 5.1 | 3.1 | 7.2 | EF |
| P17 | PRE | N/A | 50.3 | 13.4 | 0.2 | 0.2 | ACTIVE PILL |
| P17 | POST | N/A | 71.3 | 10 | 0.4 | 0.4 | ACTIVE PILL |
| P18 | PRE | N/A | 329.3 | 35.9 | 1.1 | 3.4 | ML |
| P18 | POST | N/A | 765.6 | 8.4 | 10.1 | 6.1 | **LF** |
| P19 | PRE | N/A | 81.9 | 8.3 | 8.9 | 4.9 | ACTIVE PILL |
| P19 | POST | N/A | 152.2 | 7.8 | 8.7 | 5.7 | ACTIVE PILL |
| P20 | PRE | Day 7 (out of 25) | 135.7 | 12.3 | 2.8 | 3.9 | EF |
| P20 | POST | Day 15 (out of 25) | 725.3 | 10.9 | 8 | 2.6 | **LF** |
| P21 | PRE | N/A | 107.4 | 16.4 | 6.5 | 7.3 | ACTIVE PILL |
| P22 | PRE | Day 20 (out of 28) | 604.1 | 7.4 | 3.6 | 3.5 | ML |
| P23 | PRE | N/A | 183.3 | 5.2 | 6.8 | 8.8 | EF |
| P24 | PRE | N/A | 62.9 | 7.2 | 0.2 | 0.5 | ACTIVE PILL |
| P25 | PRE | Day 16 (out of 29) | N/A | N/A | N/A | N/A | EL |
| P26 | PRE | N/A | 50.7 | 12.9 | 0.2 | 0.6 | ACTIVE PILL |
| P27 | PRE | N/A | 83.5 | 8.5421 | 0.2 | 0.5 | ACTIVE PILL |
| P27 | POST | N/A | 62.1 | 9.8872 | 0.6 | 1.4 | ACTIVE PILL |
| P28 | PRE | Day 6 (of 28) | 157.2 | 5.1909 | 9.6 | 6.7 | EF |
| P28 | POST | Day 5 (of 27) | 1296.7 | 72.0561 | 8.1 | 2 | **LF** |
| P29 | PRE | N/A | 183.5 | 4.189 | 4.1 | 7 | **PLACEBO PILL** |
| P29 | PRE | N/A | 69.5 | 7.016 | 3.3 | 5.1 | ACTIVE PILL |
| P30 | PRE | Day 1 (out of 31) | 438.3 | 10.3182 | 19.9 | 8.7 | EF |
| P30 | POST | Day 28 (out of 30) | 175.1 | 5.4004 | 12.9 | 7.7 | ML |
| P31 | PRE | Day 3 (of 25) | 85.6 | 7.77 | 5.6 | 5.6 | EF |
| P31 | POST | Day 3 (of 33) | 84 | 5.8703 | 5.6 | 6.6 | EF |
| P32 | PRE | N/A | 212.2 | 5.1581 | 11.9 | 7 | EF |
| P33 | PRE | N/A | 273.1 | 29.3536 | 3.2 | 2.4 | ML |
| P34 | PRE | Day 4 (of 24) | 140.1 | 6.1849 | 4.8 | 7.6 | EF |
| P34 | POST | Day 5 (of N/A) | 112.9 | 5.0798 | 8.3 | 6.8 | EF |
| P35 | PRE | Day 29 (of 29) | 107.2 | 7.9207 | 3.3 | 8.7 | ML |
| P35 | POST | Day 13 (of 29) | 195.6 | 4.2692 | 3.2 | 5.4 | EF |
