## Supplementary table 2 for "Total testosterone is not associated with muscle mass, function or exercise adaptations in pre-menopausal females"

**Supplementary Table 2.** The resistance training program prescribed to participants (n=27). The gym-based program was followed by all participants when access to the gym was possible. A sub-cohort of participants (n=10) performed a portion of their training programs using the home-based resistance training program when access to the training facility was not possible. *last set was prescribed as many repetitions as possible (AMRAP).

| Gym-based resistance training program | | | | | | |
| --- | --- | --- | --- | --- | --- | --- |
| Exercise | **Sets** | **Reps** | **Intensity (%RM)** | **Rest between sets (s)** | **Tempo (s)** | **Volume load (reps × sets × intensity)** |
| Squat | 3 | 8 | 80 | 90 | 2, 0, 2, 0 | 19.2 |
| Leg extension | 3 | 8 | 80 | 90 | 2, 0, 2, 0 | 19.2 |
| Hamstring curl | 3 | 8 | 80 | 90 | 2, 0, 2, 0 | 19.2 |
| Seated shoulder press | 3 | 9 | 70 | 90 | 2, 0, 2, 0 | 18.9 |
| Seated row | 3 | 10 | 60 | 90 | 2, 0, 2, 0 | 18.0 |
| Seated biceps curl | 3 | 9 | 70 | 90 | 2, 0, 2, 0 | 18.9 |
| Home-based resistance training program | | | | | | |
| Exercise | **Sets** | **Reps** | **Intensity (%RM)** | **Rest between sets (s)** | **Tempo (s)** | **Volume load (reps × sets × intensity)** |
| Squats | 3 | 15* | Approximately 45% RM | 90 | 2,0,2,0 | 20.25 |
| Forward lunges | 3 | 15·leg^-1*^ | Approximately 45% RM | 90 | 2,0,2,0 | 20.25 |
| Hamstring sliders | 3 | 10* | Body weight | 90 | 2,0,2,0 | N/A |
| Seated Shoulder press | 3 | 9* | 70% RM | 90 | 2,0,2,0 | 18.9 |
| Bent over row | 3 | 9* | 70% RM | 90 | 2,0,2,0 | 18.9 |
| Seated biceps curl | 3 | 9* | 70% RM | 90 | 2,0,2,0 | 18.9 |
